# A CRISPR/Cas9 screen reveals convergent dysregulation of neurological pathways within and across glycosylation pathways

**DOI:** 10.64898/2026.10.23.761351

**Authors:** Haley Jetter, Claudia Z. Han, Sarah D. Ackerman

## Abstract

Congenital disorders of glycosylation (CDG) are a broad group of disorders characterized by deficient glycosylation. CDG are caused by mutations in over 160 different genes regulating distinct steps and types of glycosylation. Despite the diversity of CDG-causing genes, patient phenotypes often overlap, with neurological symptoms present across many CDG. However, models to study conserved mechanisms underlying neurological dysfunction across CDG types are lacking. Therefore, we leveraged CRISPR/Cas9 to knockdown 15 glycosylation genes in zebrafish spanning 7 CDG classes. Through behavioral assays and in vivo imaging, we report changes in mobility, response to stimuli, and neuronal activity in many of our models. To uncover the molecular underpinnings of these phenotypes, we performed bulk-RNA sequencing on our models. We report differential expression in genes related to neurological development and function across all tested CDG classes, which will inform future hypothesis-driven studies defining new molecular targets for therapeutic intervention in CDG.

## Introduction

Glycosylation is an important post-translational modification in which sugar groups are added to proteins and lipids. Since glycosylated proteins and lipids coat most, if not all cells, glycosylation is essential for cell-cell communication and tissue integrity, as well as protein function, structure, stability, and localization^1^. Mutations in genes in glycosylation pathways lead to congenital disorders of glycosylation (CDG). CDG are a rare, but rapidly expanding group of monogenic disorders characterized by deficient glycosylation of proteins and lipids^2,3^. As glycosylation is ubiquitous, these disorders are multisystemic, causing both neurological and non-neurological symptoms. To date, over 160 CDG-causing genes have been identified^2^. These genes can be subdivided into groups based on their enzymatic function, including those that facilitate steps in individual glycosylation pathways, such as GPI-anchor biosynthesis, N-linked protein glycosylation, and GAG biosynthesis, as well as those that facilitate synthesis, transport, or interconversion of sugars or monosaccharides essential for multiple glycosylation pathways. While CDG exhibit broad genetic and clinical diversity, neurological dysfunction, including intellectual disability, developmental delay, brain atrophy, seizures, ataxia, hypotonia, and visual impairment impact quality of life across patients with many different CDG-causing mutations (**Fig. 1a**)^4,5^.

**Figure 1:**
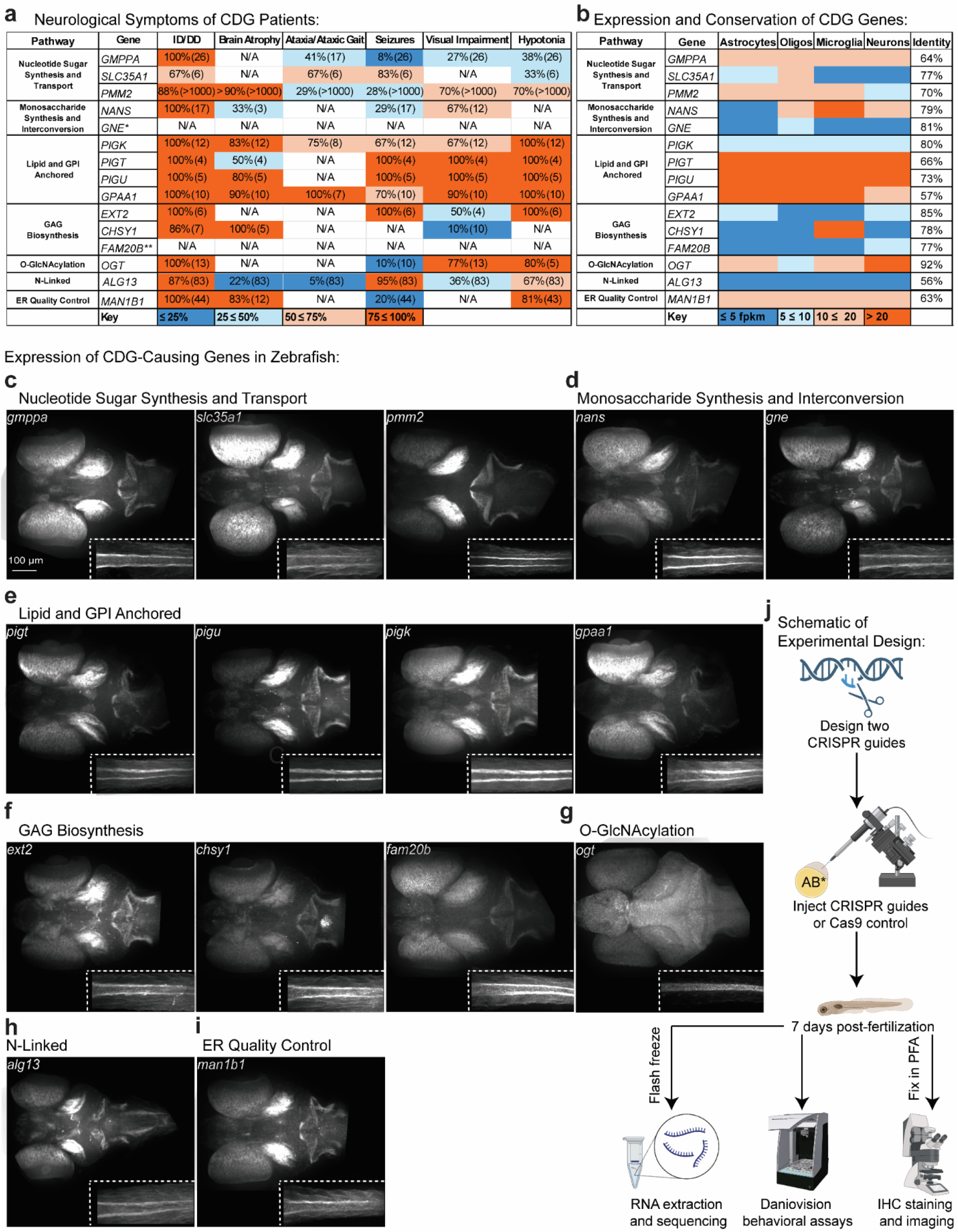
Expression and Conservation of CDG genes targeted by CRISPR/Cas9 in this study. **a:** CDG genes from seven major categories of CDG. These genes show high prevalence of neurological phenotypes in human patients. Values represent percent of reported patients with this phenotype, with the number of reported patients in parenthesis, color coded. *This CDG is characterized by muscle myopathy. The neurological symptoms listed in this table are not well characterized. **This CDG leads to perinatal lethality, thus, neurological symptoms are not characterized^91–105^. **b:** These genes show variable RNA expression in neural cell types including astrocytes, oligodendrocytes, microglia, and neurons^36^. Values represent fragments per kilobase of exon per million mapped fragments (FPKM), color-coded to denote level of expression. Identity column: protein sequence identity from zebrafish to human for each indicated gene. **c-i:** Representative images of HCR staining labeling each of the 15 genes in this study in the brain and the spinal cord (dashed inset) in WT (AB* strain) zebrafish. Images were taken at 20x. N=10 animals imaged per stain. These genes show variable expression patterns in the brain and spinal cord. **j:** Schematic of experimental design for this study. Two CRISPR guides per targeted gene were designed and injected into AB* strain WT embryos. At 7 dpf, assays including bulk RNA-sequencing, behavioral assays, and IHC imaging were performed.

Despite the prevalence of neurological dysfunction in CDG, the mechanisms underlying shared neurological phenotypes in CDG are not well-defined. One major barrier to understanding the cellular basis of CDG is a lack of models spanning all disorders^6–23^. Moreover, identifying common cellular and molecular themes across classes is hampered by the diversity of distinct animal models developed to represent any given CDG. Cross-CDG comparisons are critical, as a growing body of evidence suggests disruption of one glycosylation pathway has the potential to impact the expression of protein machinery involved in other glycosylation pathways. For example, in zebrafish, mutation of *pmm2*, which is involved in N-linked glycosylation and nucleotide sugar synthesis, increases protein O-GlcNAcylation, which, in turn, increases the abundance of enzymes essential for early stages of N-linked glycosylation^24^. This increase is protective in PMM2-CDG. Furthermore, in a yeast model of *SEC53/PMM2* deficiency, reduction of PGM1, which is important for monosaccharide interconversion, improves growth and glycosylation phenotypes^25^. Clinically, mild mutation of *ALG6,* an N-linked glycosylation gene, has higher prevalence in PMM2-CDG patients with severe symptoms than with mild symptoms, suggesting that compensation within the N-linked glycosylation pathway may modify patient outcomes^26^.

Importantly, cross-glycosylation pathway genetic interactions are observed across many CDG. In flies with inhibition of the N-glycosylation gene DPAGT1, a CRISPR enhancer/suppressor screen using RNAi in the fly eye revealed that inhibition of the mannosyltransferase DPM1, or inhibition of genes downstream of DPM1 in N-glycosylation, O-mannosylation, or GPI-anchor biosynthesis, each rescued eye area^27^. Additionally, whole genome sequencing of a family with incomplete penetrance of PIGA-CDG, a lipid and GPI-anchored CDG, revealed 4 variants in glycosylation genes as potential protective genetic modifiers^28^. Taken together, these data highlight the importance of understanding the relationships between glycosylation pathways and developing models that allow for comparison across multiple CDG classes.

Here, we leveraged zebrafish as a model system to knockdown (KD) 15 different glycosylation genes linked to distinct CDG to identify common and unique pathways disrupted across glycosylation classes. Zebrafish are the smallest genetically-tractable vertebrate model organism, and they are highly amenable to genome editing (e.g. CRISPR/Cas9 and TALENs)^29–32^. Many of the genes that cause CDG with neurological phenotypes are moderately to strongly conserved from humans to zebrafish (>50% sequence identity, **Fig. 1b**)^33^. Moreover, zebrafish motor behaviors, including seizure-like behaviors, are well-characterized and easy to assess in a laboratory setting^34,35^. Finally, the optical transparency of zebrafish allows for easy imaging of the brain during development. We find altered behavior and/or neuronal activity by quantification of phosphorylated ERK/total ERKin 11/15 of our KD models. Through bulk RNA-sequencing, we identify dysregulation of neurological pathways across all our manipulations. Ultimately, our findings will serve as a major resource to the broader community of glycosylation researchers and will be used to inform future hypothesis-driven studies defining new molecular targets for therapeutic intervention in CDG.

## Results

In this study, we performed a CRISPR-based screen targeting 15 glycosylation genes (**Fig. 1a-b**). Our gene selection criteria is that the gene causes a CDG in humans with neurological phenotypes, is highly expressed in human^36^ and fish brain cells, and is conserved from fish to humans (**Fig. 1a-i**). We designed two complementary guide RNAs (gRNAs) targeting each gene of interest. Because the teleost lineage, including zebrafish) underwent a whole genome duplication event, some genes are retained as two paralogous copies. In our gene set, *man1b1, gmppa,* and *nans* are represented by paralogs (*man1b1a/b, gmppaa/b,* and *nansa/b,* respectively). For these genes, we designed two gRNAs per paralog, resulting in four gRNAs per gene pair.. For simplicity, these genes are referenced without the *a/b* nomenclature throughout the text. We validated cutting efficiency of the gRNAs for each target using a T7 endonuclease assay^37^, and when possible, western blot (**Supp Figs. 1-2**). We then performed three assays, behavior, neuronal activity, and transcriptomic profiling, across all animals at 7 days post-fertilization (dpf), when the larval nervous system is largely mature^38–40^. To assess neurological phenotypes such as ataxia and epilepsy, we measured larval locomotion at baseline and in response to changes in light using our in-house DanioVision rig with EthoVision software. All animals were screened for changes to development prior to behavioral tests; only animals with grossly normal morphology were subjected to the DanioVision (**Supp Fig. 3,** **Fig 1j**). In parallel, we leveraged antibody staining to measure phosphorylated ERK/total ERK (pERK/tERK) as a proxy for neuronal activity^41^ in the brain and spinal cord. Finally, we used bulk RNA sequencing to measure changes in gene expression and to find conserved molecular changes across multiple CDG (**Fig. 1j**).

### Knockdown of lipid and GPI-anchored CDG genes leads to reduced mobility and neuronal activity

We measured behavioral changes following KD of four lipid and GPI-anchored CDG genes. At baseline, *pigt, pigk,* and *pigu* KD animals (e.g. “crispants”) show significantly reduced motility, and *gpaa1* crispants show a trending decrease in motility in lighted conditions. *pigt* and *gpaa1* crispants also display significantly decreased baseline motility in the dark (**Fig. 2b**). Next, we tested how changes in light impact behavior using light startle^41^ and dark transition assays. In the light startle assay, we acclimate fish to dark for 1.5 hours, followed by a 1-second light stimulus. The stimulus is repeated every 30 seconds, 5 times before returning the fish to dark for a 10-minute recovery period (**Fig. 2a**). Consistent with our baseline motility results, *pigt* and *pigu* crispants show significantly reduced response to light startle (**Fig. 2c-d**). While not statistically significant, both *pigk* and *gpaa1* crispants show a trending decrease in response to light startle (**Fig. 2e-f**). Next, for the dark transition assay, fish were acclimated to light for 10 minutes. The light was then turned off, and the animal’s response to this transition was recorded for 10 minutes (**Fig. 2a**). In wildtype, an abrupt transition from light to dark drives a startle response^42^. *pigt* and *gpaa1* crispants showed reduced response to dark transition, while *pigu* and *pigk* crispants exhibited no change in response compared to control siblings.

**Figure 2:**
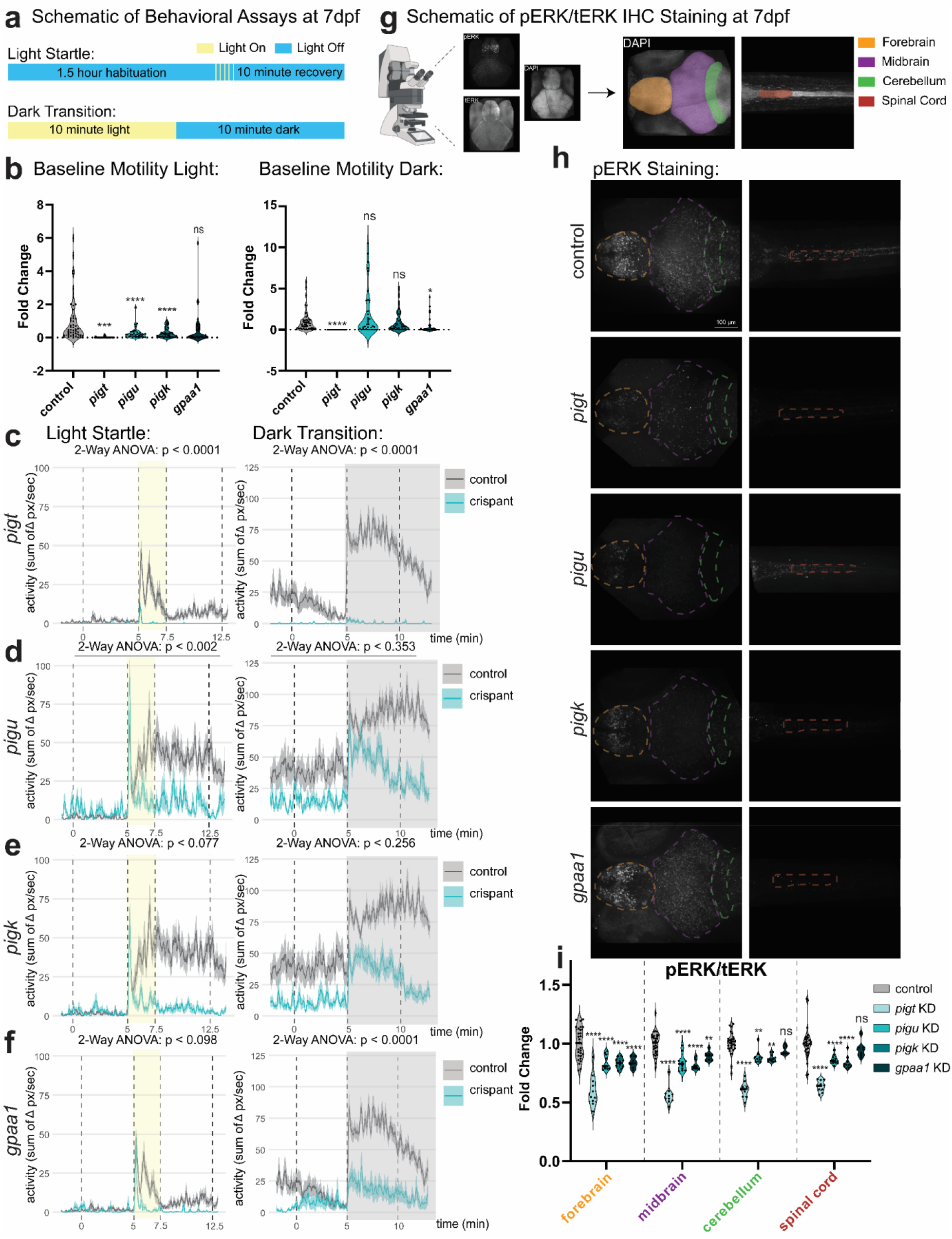
Knockdown of lipid and GPI-anchored CDG genes leads to reduced mobility and neuronal activity. **a:** Schematics of behavioral assays. Top: in the light startle assay, zebrafish were habituated to the dark for 1.5 hours, then startled with a 1-second light stimulus, repeated every 30 seconds, 5 times. Bottom: In the dark transition assay, zebrafish are habituated to the light for 10 minutes. Then, the light is turned off for 10 minutes. **b:** Baseline motility of fish in the light (left) and dark (right). Activity level was averaged over 10 minutes, and crispants (n=36 per group) were normalized to controls (n=24 per group). Statistics by one-way ANOVA, multiple comparisons by Dunnett’s multiple comparisons test. P-values vs. control in light, left to right: 0.0008, < 0.0001, < .0001, 0.1073. P-values vs. control in dark, left to right: < 0.0001, 0.1395, 0.9831, 0.0400. **c-f:** Activity traces showing response to the light startle assay (left) and dark transition assay (right) for (**c**) *pigt*, (**d**), *pigu*, (**e**) *pigk*, and (**f**) *gpaa1* crispants (n=36 per group) and controls (n=24 per group). Statistics by 2-way ANOVA. P-values shown on graphs represent time x genotype interaction. F-values for time x genotype interaction vs. control for light startle top to bottom: 67.53, 6.674, 3.066, 2.376. F-values for time x genotype interaction vs. control for dark transition top to bottom: 103.4, 0.8798, 1.322, 32.60. **g:** Schematic of pERK/tERK IHC staining. Whole fish were stained using antibody targeting total ERK, phosphorylated ERK, and DAPI, and brains and spinal cords were imaged. Forebrain (orange), midbrain (purple), cerebellum (green), and spinal cord (red), were analyzed. **h:** Representative images showing pERK staining in crispants and controls. Images from top to bottom: control, *pigt* KD*, pigu* KD*, pigk* KD*, gpaa1* KD. **i:** Quantification of pERK/tERK by region. Crispants (n=10 *pigt* KD, 10 *pigu* KD, 10 *pigk* KD, 8 *gpaa1* KD) were normalized to controls (n= 10 per group). Statistics by 2-way ANOVA, multiple comparisons by Sidak’s multiple comparisons test. P-values vs. control, left to right: forebrain: < 0.0001, < 0.0001, < 0.0001, < 0.0001; midbrain: < 0.0001, < 0.0001, < 0.0001, 0.0020; cerebellum: < 0.0001, 0.0045, 0.0015, 0.2439; spinal cord: < 0.0001, < 0.0001, < 0.0001, 0.4601. *p < .05, **p < .01, ***p < .001, ****p < .0001.

Alterations in behavior can result from changes in neuronal activity. To determine how behavior correlates to neuronal activity, we used immunohistochemical (IHC) staining with antibodies targeting phosphorylated erk (pERK) and total ERK (tERK). Since pERK labels recently active neurons, while tERK labels all neurons, the ratio of pixel intensity for pERK/tERK can be used to measure the relative activity level of neurons in various brain regions (forebrain, midbrain, and cerebellum), as well as in the spinal cord. To delineate these regions, we counterstained with DAPI to label all cells (**Fig. 2g**). Consistent with our behavioral analysis, all lipid and GPI-anchored crispants displayed evidence of reduced neuronal activity (**Fig. 2h-i**). Specifically, *pigt, pigu,* and *pigk* KD significantly reduced pERK/tERK in every region tested (forebrain, midbrain, cerebellum, and spinal cord), whereas *gpaa1* KD had significantly reduced pERK/tERK only in the forebrain and midbrain. Overall, our behavioral and cellular results showed a consistent reduction in movement and neuronal activity across all four lipid and GPI-anchored CDG genes tested.

### Knockdown of GAG Biosynthesis CDG genes alters responses to light and neuronal activity

We measured behavioral changes in three GAG Biosynthesis CDG crispants: *ext2, chsy1,* and *fam20b*. We found no changes in baseline motility in the light or in the dark compared to control siblings (**Fig. 3a**). However, these crispants did show altered responses to changes in light in the light startle and dark transition assays. *fam20b* KD resulted in reduced response to light stimuli in the light startle assay. In the dark transition assay, *ext2* KD fish displayed increased response and motility (**Fig. 3b-d**). Interestingly, *ext2* crispants displayed reduced pERK/tERK in the forebrain, despite increased motility in the dark transition assay. No changes were observed in the midbrain, cerebellum or spinal cord for any of the three GAG biosynthesis CDG crispants (**Fig 3e-f**).

**Figure 3:**
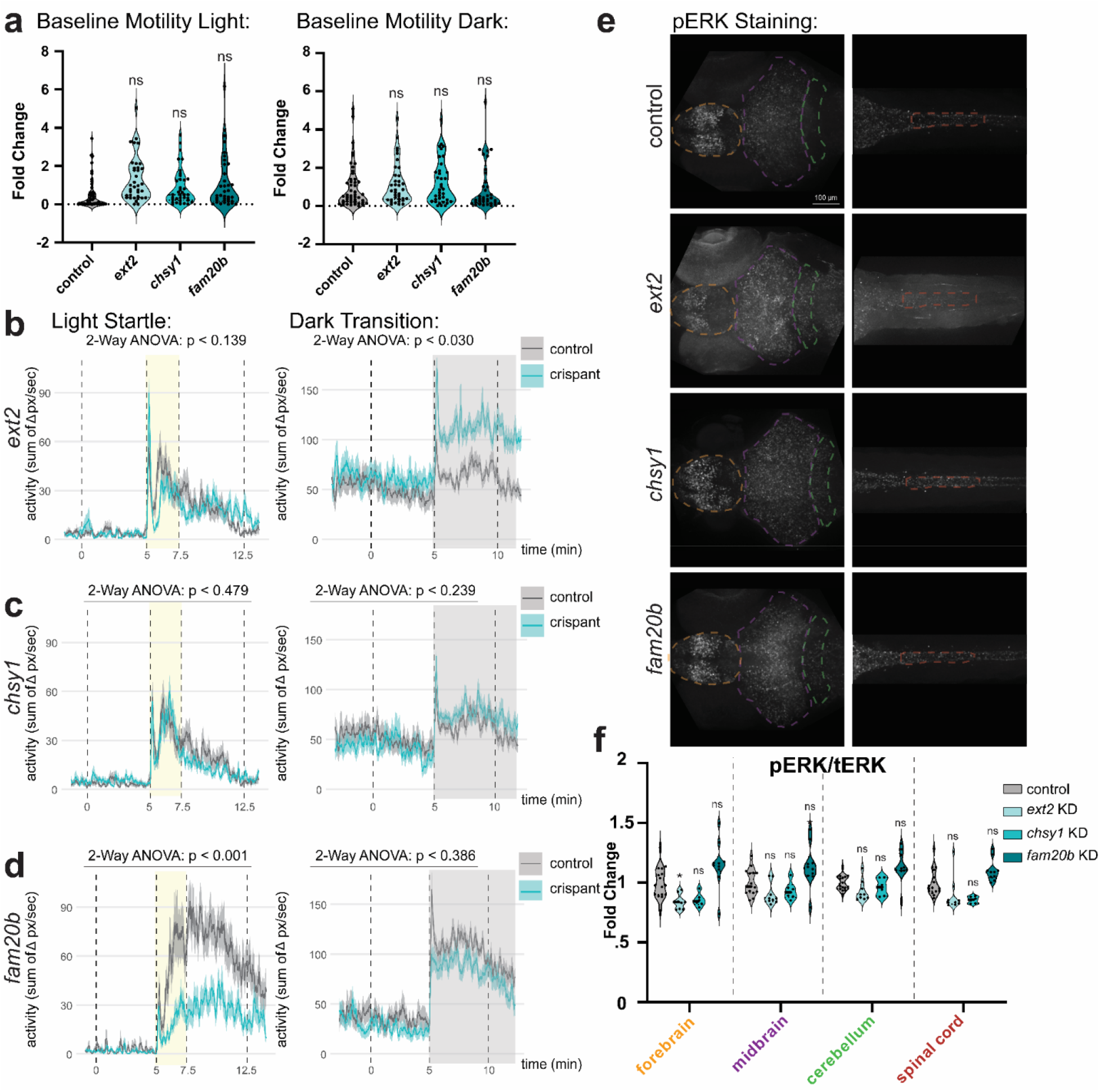
Knockdown of GAG Biosynthesis CDG genes leads to altered responses to light and neuronal activity. **a:** Baseline motility of fish in the light (left) and dark (right). Activity level was averaged over 10 minutes, and crispants (n=36 per group) were normalized to controls (n=24 per group). Statistics by one-way ANOVA, multiple comparisons by Dunnett’s multiple comparisons test. p-values vs. control, left to right. p-values vs. control in light, left to right: 0.3800, 0.9121, 0.8582. p-values vs. control in dark, left to right: 0.7946, 0.6060, 0.9629. **b-d:** Activity traces showing response to the light startle assay (left) and dark transition assay (right) for (**b**) *ext2*, (**c**), *chsy1*, and (**d**) *fam20b* crispants (n=36 per group) and controls (n=24 per group). Statistics by 2-way ANOVA. P-values shown on graphs represent time x genotype interaction. F-values for time x genotype interaction vs. control for light startle top to bottom: 2.180, 0.7413, 7.460. F-values for time x genotype interaction vs. control for dark transition top to bottom: 4.985, 1.419, 0.7633). **e:** Representative images showing pERK staining in crispants and controls. Images from top to bottom: control, *ext2* KD, *chsy1* KD, and *fam20b* KD. **f:** Quantification of pERK/tERK by region. Crispants (n=9 *ext2* KD, 6 *chsy1* KD, 10 *fam20b* KD) were normalized to controls (n= 6 control for *ext2* KD and *chsy1* KD, n = 10 control for *fam20b* KD). Statistics by 2-way ANOVA, multiple comparisons by Sidak’s multiple comparisons test. P-values vs. control, left to right: forebrain: 0.0276, 0.1207, 0.9190; midbrain: 0.4155, 0.8970, 0.9441; cerebellum: 0.8061, 0.9979, 0.9633; spinal cord: 0.3669, 0.1321, 0.9932. *p < .05, **p < .01, ***p < .001, ****p < .0001.

### Knockdown of CDG genes impacting multiple pathways leads to altered mobility, responses to light, and neuronal activity

We measured behavioral changes following KD of five CDG genes impacting multiple glycosylation pathways: three impacting nucleotide sugar synthesis and transport (*gmppa, slc35a1,* and *pmm2)* and two impacting monosaccharide synthesis and interconversion (*nans* and *gne*). We found variable changes in behavior. Only *slc35a1* KD showed changes in baseline motility, with crispants exhibiting reduced motility compared to controls in both the light and the dark (**Fig. 4a**). No crispants showed significant differences in response to light in either the light startle or dark transition assay (**Fig. 4b-f**); however, *gne* crispants showed a trending increase in response compared to controls in both assays (**Fig. 4f**). Consistently, *gne* crispants display increased pERK/tERK in the forebrain. Conversely, *gmppa* crispants showed decreased pERK/tERK in the forebrain, midbrain, and cerebellum. While not statistically significant, *slc35a1* crispants displayed a trending decrease in pERK/tERK compared to controls, consistent with the decrease in baseline motility in *slc35a1* crispants (**Fig. 4g-h**).

**Figure 4:**
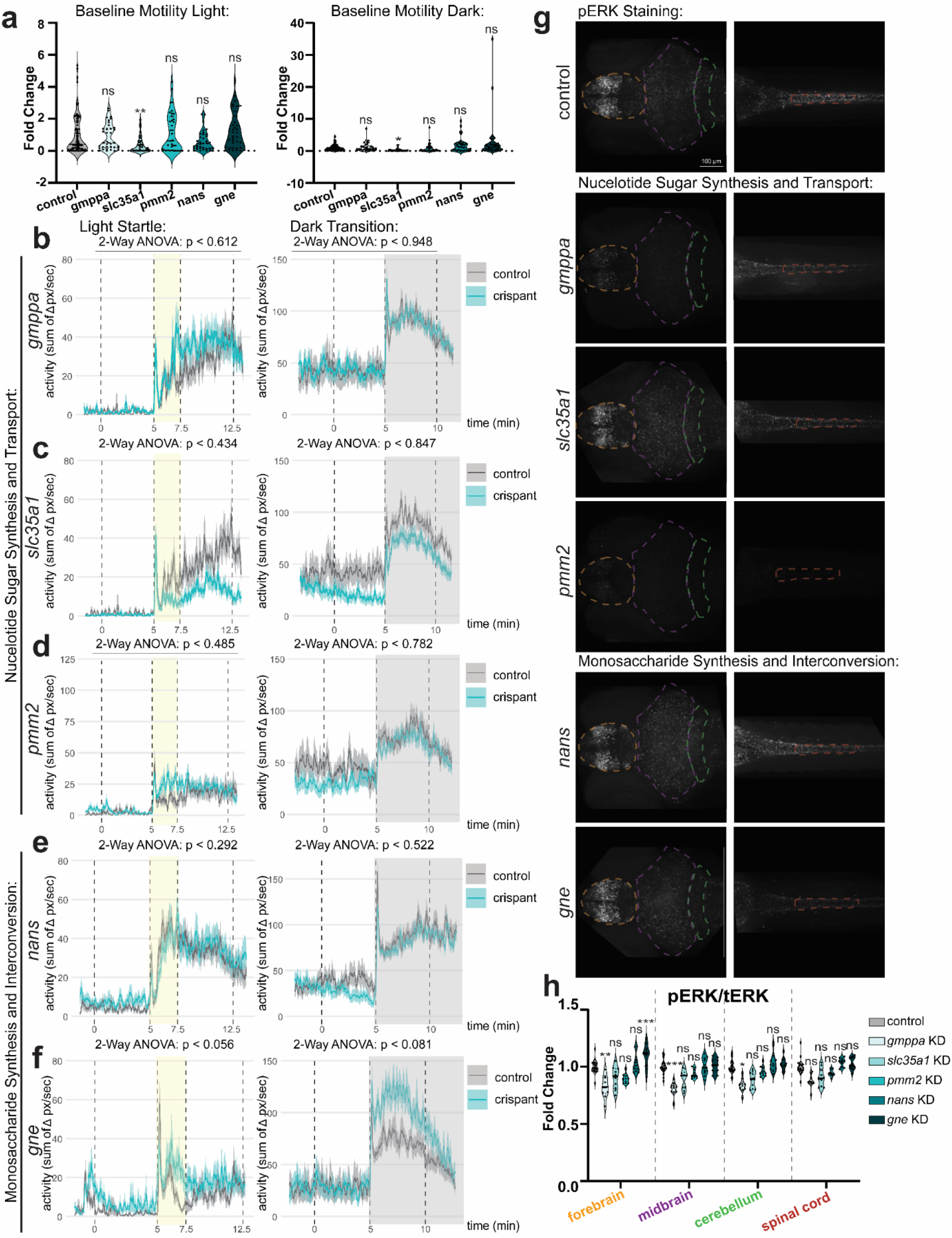
Knockdown of CDG genes impacting multiple pathways leads to altered mobility, responses to light, and neuronal activity. **a:** Baseline motility of fish in the light (left) and dark (right). Activity level was averaged over 10 minutes, and crispants (n=36 per group) were normalized to controls (n=24 per group). (Statistics by one-way ANOVA, multiple comparisons by Dunnett’s multiple comparisons test for *gmppa, slc35a1,* and *nans*. Statistics by Welch’s t-test for *pmm2* and *gne*. P-values vs. control in light, left to right: 0.7996, 0.0088, 0.5704, 0.1189, 0.5486. P-values vs. control in dark, left to right: 0.8289, 0.0156, 0.8437, 0.6747, 0.0867). **b-f:** Activity traces showing response to the light startle assay (left) and dark transition assay (right) for (**b**) *gmppa*, (**c**) *slc35a1,* (**d**) *pmm2*, (**e**) *nans*, and (**f**) *gne* crispants (n=36 per group) and controls (n=24 per group). Statistics by 2-way ANOVA. P-values shown on graphs represent time x genotype interaction. F-values for time x genotype interaction vs. control for light startle top to bottom: 0.4946, 0.6642, 0.7284, 1.229, 2.974. F-values for time x genotype interaction vs. control for dark transition top to bottom: 0.004347, 0.03765, 0.07771, 0.4153, 3.156. **g:** Representative images showing pERK staining in crispants and controls. Images from top to bottom: control, *gmppa* KD, *slc35a1* KD, *pmm2* KD, *nans* KD, and *gne* KD. **h:** Quantification of pERK/tERK by region. Crispants (n=10 per group) were normalized to controls (n= 10 per group). Statistics by 2-way ANOVA, multiple comparisons by Sidak’s multiple comparisons test. P-values vs. control, left to right: forebrain: 0.0026, 0.2705, > 0.9999, 0.9832, 0.0005; midbrain: 0.0007, 0.1551, > 0.9999, > 0.9999, > 0.9999; cerebellum: 0.0145, 0.2000, > 0.9999, 0.9700, 0.9718; spinal cord: 0.1166, 0.4988, 0.9969, 0.9008, 0.9313. *p < .05, **p < .01, ***p < .001, ****p < .0001.

### Knockdown of other CDG genes leads to variable changes in mobility, responses to light, and neuronal activity

Finally, we knocked down 1 gene related to N-linked glycosylation (*alg13*), 1 gene related to O-GlcNAcylation (*ogt*), and 1 gene related to ER quality control (*man1b1*). None of these crispants showed changes in baseline motility in the light or the dark (**Supp. Fig. 4a**). Consistently, KD of *alg13* and *man1b1* did not lead to any changes in response to light in the light startle or dark transition assay (**Supp. Fig. 4b,d**). However, *man1b1* KD zebrafish showed a significant increase in pERK/tERK in the forebrain, midbrain, and cerebellum (**Supp. Fig. 4e-f**). *ogt* crispants showed a statistically significant decrease in response to both light startle and dark transition (**Supp. Fig. 4c**). Since ERK is a target of O-GlcNAcylation, pERK/tERK staining cannot be used as a proxy of neuronal activity in *ogt* crispants. In fact, the phosphorylation and O-GlcNAcylation sites in ERK are known to compete^43,44^. Accordingly, reduction of Ogt-mediated O-GlcNAcylation leads to increased pERK/tERK, likely due to reduced competition for phosphorylation sites (**Supp. Fig. 4e-f**).

### Bulk RNA sequencing reveals altered expression of genes essential for neural structure and function across all CDG classes

Overall, we identified statistically significant changes in baseline and light-evoked motility in 8/15 of our crispant populations relative to control, which frequently coincided with changes in pERK/tERK staining in one or more brain regions. To explore the molecular basis of these changes, we measured gene expression via bulk RNA sequencing. RNA was collected from pools of whole fish at 7 dpf that were siblings of the animals analyzed in our DanioVision behavior rig (**Figs. 2-4, Supp. Fig. 4**). Crispants across all CDG pathways showed significant, differentially expressed genes (DEGs) compared to control siblings (**Fig. 5a-g**). However, the number of DEGs varied based on CDG class. For example, KD of all lipid and GPI-anchored genes displayed at least 291 upregulated and 172 downregulated significant DEGs (**Fig. 5c**). Conversely, KD of our sole N-linked glycosylation (*alg13*) gene resulted in few DEGs (24 DEGs total, **Fig. 5f**), despite high cutting efficiency of both *alg13* CRISPR guides (**Supp. Fig. 1a,e**).

**Figure 5:**
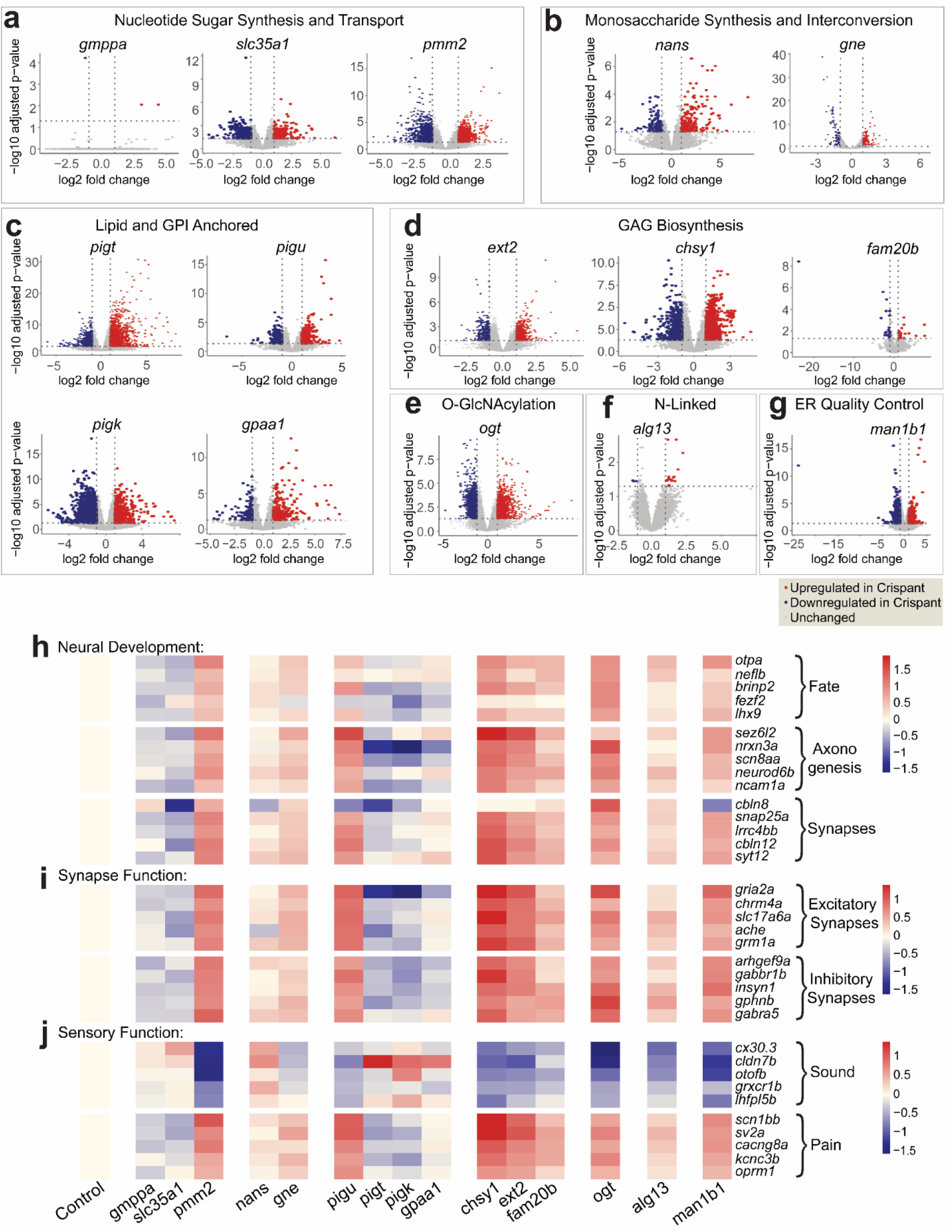
Neurological pathways are dysregulated across all tested CDG classes. **a-g:** Volcano plots showing significantly up-(red) and downregulated (blue) genes in each manipulation, grouped by glycosylation gene function. Upregulated genes have adjusted p-value < .05 and log2(fold change) ≥ 1, and downregulated genes have adjusted p-value < .05 and log2(fold change ≤ -1. **h-j:** Heatmaps displaying log2(fold change) of selected genes related to neurological development and function compared to control, colored to represent level of expression, and subdivided by glycosylation gene function. **h:** Genes related to neural development, including those essential for neuron fate, axonogenesis, and synapse formation. **i:** Genes critical for excitatory and inhibitory synapse function. **j:** Genes related to sensory function, such as perception of sound and pain.

To assess dysregulated pathways across manipulations, we performed gene ontology (GO) enrichment analysis of significantly up- and downregulated genes (adjusted p-value < .05, log2(fold change) ≥ 1 or ≤ -1) using Metascape^45^ (**Supp. Figs. 5-8**). We found enrichment for neural-related pathways in 13/14 crispants with sufficient DEGs to analyze using the platform.

To further characterize changes in gene expression across neural development, we selected representative genes important for different aspects of neuron development and function for closer analysis. Neural development can be broken into three essential processes. First, cells must undergo fate specification to differentiate into neurons. Next, neurons undergo axonogenesis, in which they initiate axon formation and extend their axon to the correct target location. Finally, neurons undergo synaptogenesis, in which they form and organize synapses to wire into circuits. Genes related to each of these three steps, including fate-driving transcription factors (*fezf2*^46^ and *lhx9*^47^), axon guidance cues (*ncam1a*^48^), and synaptic adhesion proteins (l*rrc4bb*^49^) were dysregulated in our dataset, suggesting broad changes in neural circuit assembly (**Fig. 5h**). Notably, *ncam1a* is known to be polysialylated. Loss of *ncam1a* or of the polysialyltransferase, *st8sia2* in zebrafish leads to splitting of the normally tightly-fasciculated axon bundle in the posterior commissure^48^. Given the role for the polysialylation of *ncam1a* in axon development and structure, our data suggests that disruption of Ncam1a glycosylation may drive compensatory transcriptional upregulation of *ncam1a* transcripts. We next assessed expression of genes related to synapse function, revealing broad dysregulation of genes essential for both excitatory neurotransmission through glutamate (*gria2a*^50^) and acetylcholine (*chrm4a*^51^), as well as genes that promote the function of inhibitory GABAergic synapses (*gabbr1b*^52^*, gabra5*^53^, and *arhgef9a*^54^) (**Fig. 5i**). Since neural circuits drive behavior and sensory perception of stimuli, we also assessed the expression of genes related to the perception of sound and pain. We observed a generalized downregulated of genes related to perception of sound and increased expression of genes relation to pain perception (**Fig. 5j**). This is consistent with increased sensory modulation disorders in patients with autistic features^55,56^, which are common in CDG^57,58^.

Importantly, our crispant data shows strong similarities with published mutant data across several CDG models. First, we compared the DEGs from our *pmm2* KD zebrafish to DEGs from *pmm2* global mutant zebrafish and *PMM2* mutant human cortical organoids^10^. We assessed GO-terms for all DEGs that were dysregulated in the same direction between our dataset and the dataset of interest. We detected a statistically significant overlap in concordantly downregulated DEGs between our 7 dpf *pmm2* KD fish and published 5 dpf *pmm2* KO fish (**Supp. Fig. 9a**). These DEGs were enriched for neurological function, including “olfactory signaling pathway” and “adenylate cyclase-inhibiting serotonin receptor signaling pathway” (**Supp. Fig. 9a**). Similarly, concordant genes between our *pmm2* KD fish and *PMM2* mutant human cortical organoids were enriched for “regulation of neuron projection development” and “chemical synaptic transmission” (**Supp. Fig. 9b**). We also compared our *ogt* KD zebrafish to *Ogt* conditional knockout mouse astrocytes^59^ and *Ogt* induced knockout mouse embryonic stem cells^60^. Concordant genes between our *ogt* KD fish and *Ogt* iKO mouse embryonic stem cells were enriched for “regulation of membrane potential, glutamate receptor signaling pathway,” and “synapse organization” (**Supp. Fig. 9d**). Thus, KD or knockout of glycosylation genes across many distinct pathways caused profound changes to genes that drive each stage of nervous system development.

### Bulk RNA sequencing reveals altered expression of genes related to cellular stress and injury and immune system function across all CDG classes

Cellular stress and injury is a common mechanism in CDG^61–68^. We observed altered pathway enrichment for ER processes and cell death in 7/14 of our crispants analyzed via Metascape (**Supp. Figs. 5-8**). To further characterize changes in gene expression related to cellular stress and injury, we selected five representative genes associated with ER stress and five associated with apoptosis to analyze. Interestingly, while we observed significant increases in cell stress in three of our four Lipid and GPI-Anchored crispants, we observed strong downregulation of these pathways in most of our KD conditions (**Supp. Fig. 10a**).

Changes in ER stress and apoptosis can lead to immune system activation^69^, and interesting, altered immune system function is also common across CDG^70–72^. Consistently, GO-terms associated with these pathways were enriched in 12/14 crispants analyzed using Metascape. (**Supp. Figs. 5-8**). We also saw robust changes in genes broadly related to immune system function across all tested CDG classes (**Supp. Fig. 10b**), which mirrored the trends observed for cell stress (**Supp. Fig 10a**). We selected five genes from four categories of immune function: type 1 antiviral response, interferon-gamma, inflammasome, and macrophage activation (**Supp. Fig. 10b**). Interestingly, we found strong dysregulation of inflammasome-related genes, which are essential for detecting infection or cell damage and triggering cell death, and macrophage activation-related genes, which are critical for phagocytosis and activating adaptive immunity (**Supp. Fig 10b**).

### Jaccard analysis reveals overlap between crispants in different CDG classes

Given the striking overlap in the GO terms impacted in our 15 crispant conditions, we aimed to define which conditions showed the most similar DEGs. Accordingly, we compared DEGs across each of our KD conditions using Jaccard analysis, and performed GO-term analysis (Metascape) on highly-concordant crispant pairs (**Fig. 6, Supp. Fig. 11**). Five KD conditions displayed high similarity: *pmm2, chsy1, man1b1, ext2,* and *pigu.* Notably, of these five genes, only two genes are from the same glycosylation pathway—*chsy1* and *ext2*, which both regulate GAG biosynthesis. These data suggest that changes in enzymes across distinct steps of glycosylation can drive similar transcriptomic responses (**Fig. 6b-c**). Consistent with our analysis of GO-terms for individual CDG causing genes, shared upregulated genes between highly concordant pairs of crispants displayed enrichment for pathways associated with neural development (**Fig. 6d-e, Supp. Fig. 11**). Downregulated genes were commonly enriched for immune system and apoptosis (**Fig 6d-e, Supp. Fig. 11**).

**Figure 6:**
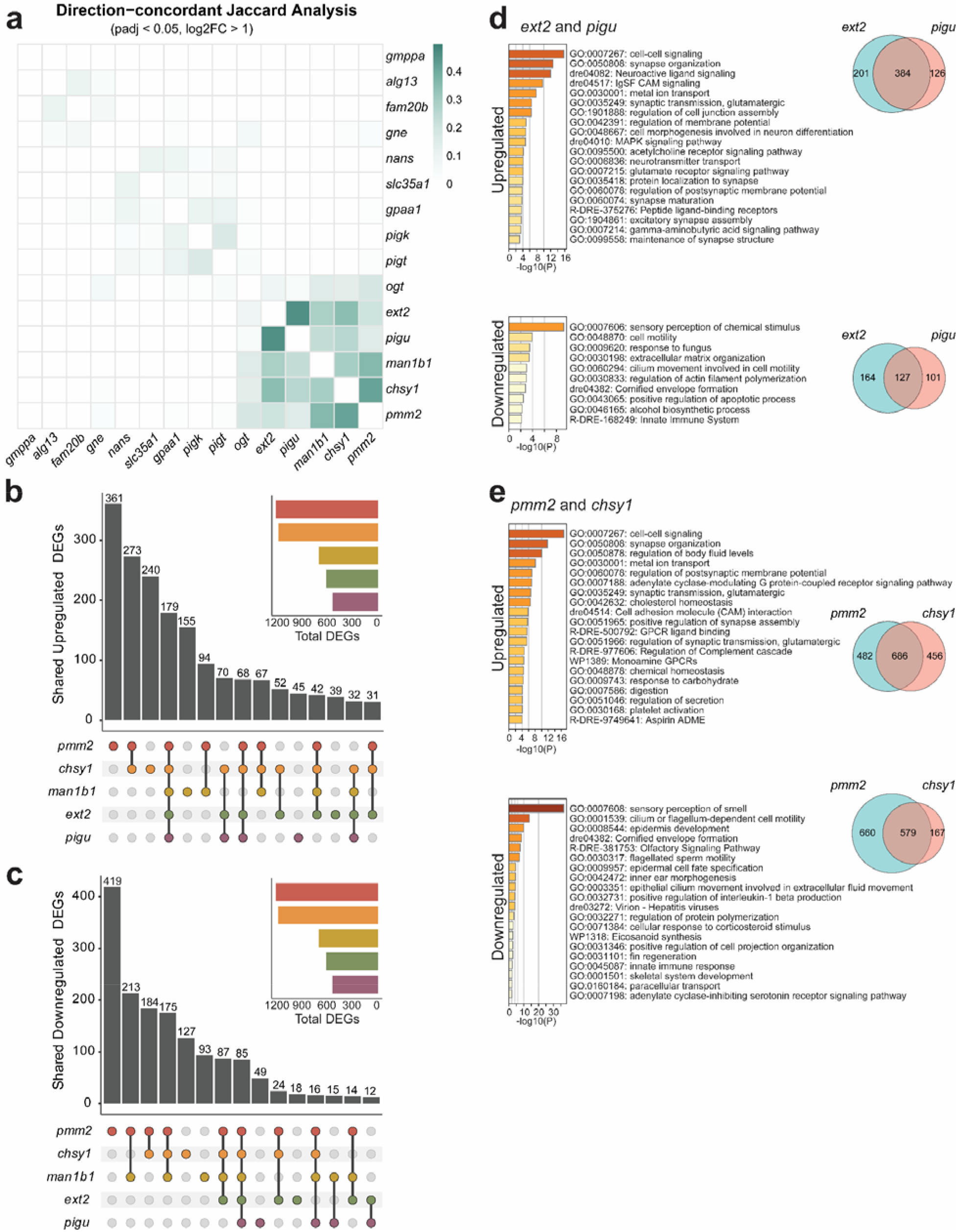
Knockdown of distinct CDG-causing genes leads to overlapping DEGs enriched for neural development, immune system function, and cellular stress and injury. **a:** Heatmap displaying Jaccard Analysis, color coded to reflect degree of overlap. **b:** upset plot showing the number of concordantly upregulated (adjusted p-value < .05, log2(Fold Change) ≥ 1) of five highly overlapping genes: *pmm2, chsy1, man1b1, ext2,* and *pigu.* c: upset plot showing the number of concordantly downregulated (adjusted p-value < .05, log2(Fold Change) ≤ -1) of five highly overlapping genes: *pmm2, chsy1, man1b1, ext2,* and *pigu.* d: The pair of genes with the highest overlap is *ext2* and *pigu*. **Left:** GO-term analyses (Metascape) of concordantly upregulated (top, adjusted p-value < .05, log2(fold change) ≥ 1) and downregulated (bottom, adjusted p-value > .05, log2(fold change) ≤ -1) genes in this pair. **Right:** Venn diagrams of concordantly upregulated (top) and downregulated (bottom) genes in this pair. **e:** The pair of genes with the second highest overlap is *pmm2* and *chsy1*. **Left:** GO-term analyses of concordantly upregulated (top) and downregulated (bottom) genes in this pair. **Right:** Venn diagrams of concordantly upregulated (top) and downregulated (bottom) genes in this pair.

## Discussion

Mutation of over 160 genes can lead to CDG^2^; however, identifying conserved cellular and molecular themes across type is hampered by the diversity of distinct animal models that have been generated to represent individual CDG^6–23^. Here, we leveraged zebrafish to generate CRISPR/Cas9 KDs of 15 distinct CDG-causing genes spanning 7 glycosylation gene classes. We assessed behavior, neuronal activity, and gene expression in each KD condition. Behavioral and pERK/tERK alterations were common across our KD animals that span seven categories of glycosylation genes, implicating neurological dysfunction as a common theme in CDG. Indeed, this is consistent with human patients, as neurological symptoms are common sources of discomfort in patients^4,5^.

Most strikingly, KD of all four lipid and GPI-anchored glycosylation genes we tested (*pigt, pigu, pigk,* and *gpaa1*) led to a profound reduction in motility both at baseline and in response to light and dark stimuli. Each of our lipid and GPI-anchored crispants also showed significant reductions in pERK/tERK across brain tissues. Interestingly, all four of our tested lipid and GPI-anchored genes belong to the GPI transamidase complex. Unlike many CDG genes, which are responsible for building and elongating glycans or synthesizing sugars necessary for glycosylation, the GPI transamidase complex catalyzes the attachment of GPI anchors to target proteins and lipids^73^. Clinically, patients with lipid and GPI-anchored CDG can be categorized into two groups depending on the function of the mutated gene: synthesis, which includes enzymes mediating assembly of the GPI precursor backbone, or transamidase+remodeling^74^. Patients in the transamidase+remodeling group have higher prevalence of every neurological symptom recorded than those in the synthesis group, including sleep disturbance, neurodevelopmental abnormality, behavioral abnormality, neurodevelopmental delay, and intellectual disability^74^. Together, these data indicate a critical role for the GPI transamidase complex in movement and neuronal activity, and suggest that compensation may be more difficult at late glycosylation steps, such as transferring glycans, compared with earlier glycan building steps.

Changes in glycosylation can have broad transcriptional impact. RNA can be indirectly glycosylated^75^. Like glycosylated proteins, many glycosylated RNAs are found at the cell surface and are important for cell-cell interactions, engage in canonical adhesion pathways, and are essential for functional regulation of the immune system. In addition, mutation of glycosylation genes can drive compensatory changes in the transcripts of diverse glycosylation enzymes^24–28^. Finally, altered glycosylation of proteins in the ER and golgi can activate the unfolded protein response pathway, which drives substantial changes in gene transcription as the system tries to reestablish proteostasis^76,77^. Thus, we performed bulk RNA sequencing of whole animals for all 15 CDG KD models to explore common and divergent gene expression changes. To make the results of this assay an easily-accessible and understandable resource for CDG patients and caregivers, as well as researchers, we created a website, daniosugar.com. On this site, users can search for any gene of interest and view its log2(fold change) expression compared to control in each of our 15 manipulations.

Via GO-term^45^ analysis we identified core cellular and molecular pathways that are upregulated or downregulated in each of our KDs. Strikingly, in 13/14 manipulations, we found enrichment for neural-related pathways. Sweeping changes in genes that span each stage of neuronal and neural circuit development, including in genes underlying neuronal fate specification, axon formation and extension, and synapse formation and organization, were observed in almost all crispants relative to sibling controls. We also saw broad changes in neuronal function, including genes essential for excitatory and inhibitory synapses, as well as for sensory processing of sound, vision, and pain. Both visual impairment, such as Retinitis pigmentosa and myopia, and hearing impairments, such as sensorineural hearing loss, have been previously reported in CDG patients^78,79^. These changes are conserved across CDG class and across datasets (**Supp. Fig. 9-10**), providing avenues for future studies of neurological dysfunction in many different CDG.

Since glycosylation is ubiquitous in the human body, CDG are multisystemic disorders. Consistent with previous studies, our data reveals common immune system and cellular stress and injury-related themes across our distinct crispant classes^61–72^. Emerging data indicate strong relationships between immune system state and brain function^80^. Common DEGs represent strong candidates to be investigated in future mechanistic studies to test the relationship between immune system dysfunction and neurological phenotypes in CDG.

## Limitations of the study

In sum, our work highlights the necessity for comparable CDG models to determine conserved mechanisms for neurological dysfunction across CDG class, though several key limitations must be addressed. Patients with CDG have partial loss-of-function genetic mutations^1,2^. While our CRISPR/Cas9 KD system models loss of function in CDG-causing genes more accurately than complete gene knockout, we did not generate humanized alleles to exactly model patient genetics. Additionally, since the study was performed in F0 crispants, it is possible that findings are impacted by CRISPR guide efficiency, as our KDs were mosaic.

Given the number of distinct KD conditions tested, our study was performed at only one timepoint (7 dpf). It is therefore possible that phenotypes arising later in development may be missed, or that maternal deposition of wildtype mRNA attenuates some phenotypes at this early timepoint. Future studies using stable CRISPR KO models and performing experiments at multiple timepoints throughout development will be informative.

In this study, we used pERK/tERK as a proxy of neuronal activity. Since different neuronal subtypes can be phosphorylated at variable rates, this approach may mask important activity differences between neuronal subtypes. Functional studies, including *in vivo* calcium imaging or electrophysiology will help to analyze neuronal activity more directly and across neuronal subtypes.

Finally, while changes in glycosylation can have broad transcriptional impact, glycosylation more immediately impacts proteins and lipids. Thus, glycoproteomics and lipidomics will be necessary in future studies to causally link neurological targets to CDG symptoms.

## Methods

### Fish Husbandry

Zebrafish (*Danio rerio*) were maintained in the Washington University Zebrafish Consortium facility (<u>Home | Zebrafish Facility | Washington University in St. Louis</u>). All experiments were performed in compliance with Washington University’s institutional animal protocols, as well as IACUC guidelines. Embryos were collected from pair-wise or harem matings and reared at 28° C in embryo medium (5 mM NaCl, 0.17 mM KCl, 0.33 mM CaCl_2_, 0.33 mM MgSO_4_). Larvae were maintained in temperature and light-controlled incubators within the lab matching the fish facility (14/10 light-dark cycle, 28°C). Embryos were fed living rotifers at 5 dpf,and harvested for experiments at 7 dpf. All experiments were performed in a WT (AB*) background.

### CRISPR Design and Injection

Two CRISPR guides were designed per targeted gene using CHOPCHOP^81–83^ (**Supplementary Table 1**). Parameters are set to target each gene in *Danio rerio* (danRer/GRCz11) using CRISPR/Cas9 for knock-out. When possible, one guide was designed to target exon 1, and the second guide was designed to target any downstream exon. Integrated DNA Technologies’ CRISPR-Cas9 gRNA checker was used to validate on-target and off-target effects. crRNA was annealed to tracrRNA via manufacturer’s instructions to make gRNAs. Injection mixes were prepared using the following concentrations: 5 micromolar gRNA, 5 micromolar Cas9 protein (IDT Cat # 1081059), and 1:10 phenol red dye in water. Cas9 control injection mix was prepared in the same manor, but with no gRNA. Prior to microinjection, the complex solution was incubated at 37°C for 5 minutes and then placed at room temperature. 1 nL was injected using a microinjector and glass needle into one-cell stage AB* strain WT embryos.

### T7 Endonuclease Assays

T7 endonuclease assays were performed as previously described to assess CRISPR/Cas9 injection efficiency^37^. In brief, PCR was performed to amplify the target region using gDNA isolated from injected embryos and uninjected AB* strain WT controls at 1 dpf. Primers were designed using CHOPCHOP (**Supplementary Table 2**). Amplified DNA from WT and injected embryos were annealed. Samples were then incubated at 37°C with T7 endonuclease I enzyme (New England BioLabs Cat # M0302L) for 15 minutes, run on a 2% agarose gel, and imaged. Images were analyzed using FIJI/ImageJ, and editing efficiency was calculated as Σ(cleaved band intensity)/(uncleaved band intensity) * 100.

### Protein Isolation and Western Blots

Pools of 10-15 whole fish per sample were flash frozen in 500 µl of NP40 lysis buffer at 7 dpf. NP40 lysis buffer was made using the following concentrations: 10mM Tris/Cl (pH7.5), 150mM NaCl, 0.5mM EDTA, 0.5% igepal (NP40 substitute), and 1x protease inhibitor cocktail (Thermo Scientific REF A32955). Protein was isolated from pools of fish as previously described^8^. In short, we sonicated embryos using 5 quick pulses at 40% power followed by a 15-minute incubation on ice, 3 times. We then centrifuged samples at 15,000g for 15 minutes at 4°C and saved the supernatant. Samples were quantified via Pierce BCA Protein Assay Kit (ThermoFisher Cat. 23225). Western blots were also run as previously described^8^. Briefly, we loaded 20µg of sample per well of a 4-20% Mini-PROTEAN TGX gel (Cat # 4561094) and ran at 50V for 30 minutes, 75V for 30 minutes, then 100V for 1 hour. Samples were then transferred to a nitrocellulose membrane in the cold at 100V for 2 hours. Membranes were blocked in 5% milk in 1x tris-buffered saline with .1% Tween 20 (TBST) for 1 hour at room temperature, shaking. Membranes were then incubated with primary antibodies rabbit anti-Pmm2 (ProteinTech Cat # 10666-1-AP) and rabbit anti-Beta Actin (GeneTex Cat # GTX637675) diluted in 1x TBST with 3% bovine serum albumin (BSA) at 4°C overnight, shaking. The blot was rinsed 3-5x quickly with 1x TBST to remove milk. Blots were rinsed for 10 minutes 3x in 1x TBST, then incubated with secondary antibody, anti-rabbit HRP-linked antibody, (Cell Signaling Technology Cat # 7074S) diluted in 1x TBST with 3% BSA for 1 hour at room temperature. Samples were rinsed for 10 minutes 3x in 1x TBST, then incubated with enhanced chemiluminescence (Millipore Cat. # WBKLS0100) for 5 minutes in the dark, then imaged. Blots were quantified using FIJI/ImageJ to determine relative expression of each gene compared to the loading control as previously described^84^.

### Behavioral Assays

Fish were loaded 1 per well into a round-well 96-well plate. In each plate, rows A and B contained Cas9 only-injected control fish, rows C, D, and E contained one CRISPR/Cas9 experimental group (*e.g.* crispant), and rows F, G, and H contained a second crispant group. All behavioral experiments were performed using the Daniovision behavioral rig between 3:30 and 7:00pm at 7 dpf. In the light and dark baseline motility assays, fish were habituated to light or dark for 80 minutes, then the average activity level of fish (% pixel change per frame in the well) was measured for 10 minutes in light or dark, respectively. In the light startle assay, fish were acclimated to dark for 90 minutes, then startled with a 1-second light stimulus. The stimulus was repeated every 30 seconds 5 times, as in previous studies^41^. Activity level was measured 5 minutes prior to the first stimulus, during the stimuli, and 5 minutes after the final stimulus. In the dark transition assay, fish were acclimated to light for 10 minutes, then switched to dark. Activity level was measured 5 minutes prior to and 5 minutes following the switch to dark.

### Immunohistochemical Staining and Imaging

Immunohistochemical (IHC) staining was performed as previously published^85^. In brief, fish were fixed in 4% paraformaldehyde (PFA) in 1x phosphate buffered saline (PBS) overnight at 7 dpf. Fish were washed out of PFA into .1% triton in 1x PBS, then de-pigmented in 50mM KOH, 1.5% H2O2 in sterile water for 10 minutes, two times^41^. Fish were washed out of depigmentation solution into .1% triton in 1x PBS. Next, fish were digested in 20mg/ml Prot-K diluted 1:1000 in 1x PBS for 75 minutes, then washed in .1% triton in 1x PBS Fish were post-fixed in 4% PFA supplemented with .1% triton in 1x PBS for 20 minutes, then washed in .1% triton in 1x PBS to remove fixative. Samples were blocked overnight in buffer containing .1% triton, 1% normal donkey serum, 1% normal goat serum, and 1% BSA in 1x PBS. Samples were then incubated overnight in primary antibody solution that included mouse anti-tERK (1:500, Cell Signaling Technology Cat # 4696S) and rabbit anti-pERK (1:500, Cell Signaling Technology Cat # 4370S) in 1x PBS supplemented with .1% Triton and DMSO (1:50) to improve antibody penetration. Samples were then washed overnight with 1x PBS + .1% triton. Next, samples were incubated overnight in secondary antibody, including 1:400 anti-mouse RRX (Invitrogen Cat. #R6393), 1:400 anti-rabbit 488 (Jackson Immuno Research Labs Cat. #NC0174015), and 1:10,000 DAPI in 1x PBS supplemented with .1% Triton and DMSO (1:50) to improve antibody penetration. Finally, samples were washed overnight in .1% triton in 1x PBS. Post-staining, fish were mounted dorsally in 1% low melt agar and brains and spinal cords were imaged at 20x (.8NA) using a custom-built spinning disc confocal from Intelligent Imaging Innovations (3i).

### Hybridization Chain Reaction (HCR) Staining and Imaging

We performed HCR as previously described (**doi:** https://doi.org/10.64898/2026.06.23.734083) following the manufacturer’s protocol (Molecular Instruments). HCR probe sequences can be found in **Supplementary Table 3.** For staining, 7 dpf, AB* strain WT fish were fixed in 4% PFA in 1x PBS at 4°C overnight. Fish were washed out of fixative and depigmented as described above. Fish were washed out of depigmentation solution into .1% Tween 20 in 1x PBS (PBTween). Next, fish were digested in 1:1000 Prot-K (20 mg/ml) in PBS for 75 minutes, then washed in PBTween. Fish were post-fixed in 4% PFA in PBTween for 20 minutes, then washed in PBTween. Dehydration washes were then performed using 25%, 50%, then 75% methanol in PBTween for 15 minutes each before being stored in 100% methanol at -20°C. Rehydration washes were performed with 75%, 50%, then 25% methanol in PBTween for 15 minutes each, followed by two washes in 100% PBTween. Samples were incubated with hybridization buffer for 1 hour at 37°C. Probes were diluted in hybridization buffer, then added to samples overnight at 37°C. Samples were then washed with hybridization probe wash buffer 4x for 15 minutes each at 37°C. Next, samples were washed with 5x .1% Tween-20 in saline-sodium citrate (SSCT) for 5 minutes 2x at room temperature, then incubated with amplification buffer for 1 hour at room temperature. HCR amplifiers were prepared by aliquoting hairpin 1 and hairpin 2 (H1, H2) into strip tubes, bringing the tubes to 95°C, then cooling the tubes at room temperature in the dark for 30 minutes. H1 and H2 were then mixed and added to amplification buffer. Samples were incubated with the hairpin and amplification buffer mixture overnight at room temperature in the dark. 4x 15-minute washes with 5x SSCT were performed. Finally, fish were mounted dorsally in 1% low melt agar and imaged using our 3i spinning disc confocal as described above. Controls were treated identically to samples, except no probes were added.

### RNA Extraction, Isolation, and Bulk Sequencing

Three pools of 20 fish per condition (crispant or day-matched, Cas9-injected control) were flash frozen in Qiagen’s Buffer RLT (GTIN 04053228006114, REF 74104). Samples were homogenized using a cordless pestle motor, then RNA was extracted and isolated following the Qiagen RNeasy Mini Kit (GTIN 04053228006114, REF 74104). Samples were submitted to the Washington University Genome Technology Access Center for bulk RNA-sequencing. QC was performed to ensure RIN >8 for each sample. Takara-Clontech SMARTer was used for library construction. Illumina Sequencing on a NovaSeq X Plus (300 cycles) with 30 M targeted reads was used for sequencing.

### RNA Sequencing Analysis and Visualization

Raw paired-end RNA-seq reads were processed using fastp (v0.23.4) to remove Illumina TruSeq adapter sequences, trim poly-G tails, and filter low-quality reads. Reads were quality-filtered using a Phred quality threshold of 20 and reads shorter than 50bp were discarded. Trimmed reads were aligned to the zebrafish reference genome (danRer11) using STAR (v2.7.11b), allowing a maximum of 10 mapping loci per read. The gene expression raw counts were quantified by HOMER’s^86^ analyzeRepeats command with the option “-condenseGenes -count exons -noadj”. TPM (transcript per kilobase million) were quantified for all genes matching accession number to raw counts. Differentially expressed genes were assessed with DESeq2^87^ at p-adj (adjusted pvalue) < 0.05 and FC (fold change) > 2 where indicated. Orthologous genes between zebrafish and mouse and human were identified using Ensembl BioMart^88^.

Pairwise transcriptional similarity between crispants was assessed using the Jaccard similarity index. Upregulated and downregulated DEGs were analyzed separately. Pairwise Jaccard similarity was calculated as the number of shared DEGs divided by the total number of unique DEGs across two models (intersection/union). Gene-set intersections across models were visualized using UpSet plots^89^.

Heatmaps were generated with the pheatmap packages in R and other plots were made with ggplots2 in R with colors reflecting the scores/expression values, including z-scores, as noted in each figure.

## Statistical Analysis

Behavioral analysis was performed using 36 crispants and 24 sibling controls per condition. For baseline activity analysis, statistics were performed by two-tailed one-way ANOVA with multiple comparison’s by Dunnett’s multiple comparisons test. Unless otherwise noted, for light startle assays, statistics were done by two-tailed two-way ANOVA with three timepoints: average activity for five minutes before startle, average activity during startle, and average activity for five minutes after startle. For dark transition assays, statistics were performed by two-tailed, two-way ANOVA with two timepoints: average activity for five minutes before startle and average activity for five minutes after startle. For pERK/tERK staining assays, 6-10 mutants per condition were compared to 6-10 sibling controls (exact n for each group is noted in figure legends). Statistics were performed by two tailed, two-way ANOVA, with multiple comparisons by Sidak’s multiple comparisons test. For western blot, three samples, each containing a pool of 15 fish, were run per condition. Crispant pools were compared to pools of sibling controls. Statistics were run by two-tailed unpaired t-test. Gene expression differences were calculated with DESeq2^87^ with Benjamini-Hochberg multiple testing correction^90^. Significance of gene set overlap was calculated using the one-tailed Fisher exact test, p < 0.05.

## Supporting information

Jetter2026_SupplementaryFigsAndTables

## Data Availability

Data from this study can be explored and visualized through an interactive website at daniosugar.com. Outside datasets used in this paper can be accessed at GSE1693471, GSE252758, and GSE251886.

## Author Contributions

HJ and SDA conceived and designed the study. HJ performed all experiments. SDA assisted with CRISPR injections for all experiments. HJ and CZH analyzed the data. HJ, SDA, and CZH generated figures, wrote and edited the manuscript collaboratively.

