## Supplementary material for "A CRISPR/Cas9 screen reveals convergent dysregulation of neurological pathways within and across glycosylation pathways": Jetter2026_SupplementaryFigsAndTables

**Jetter et al. 2026 Submission**

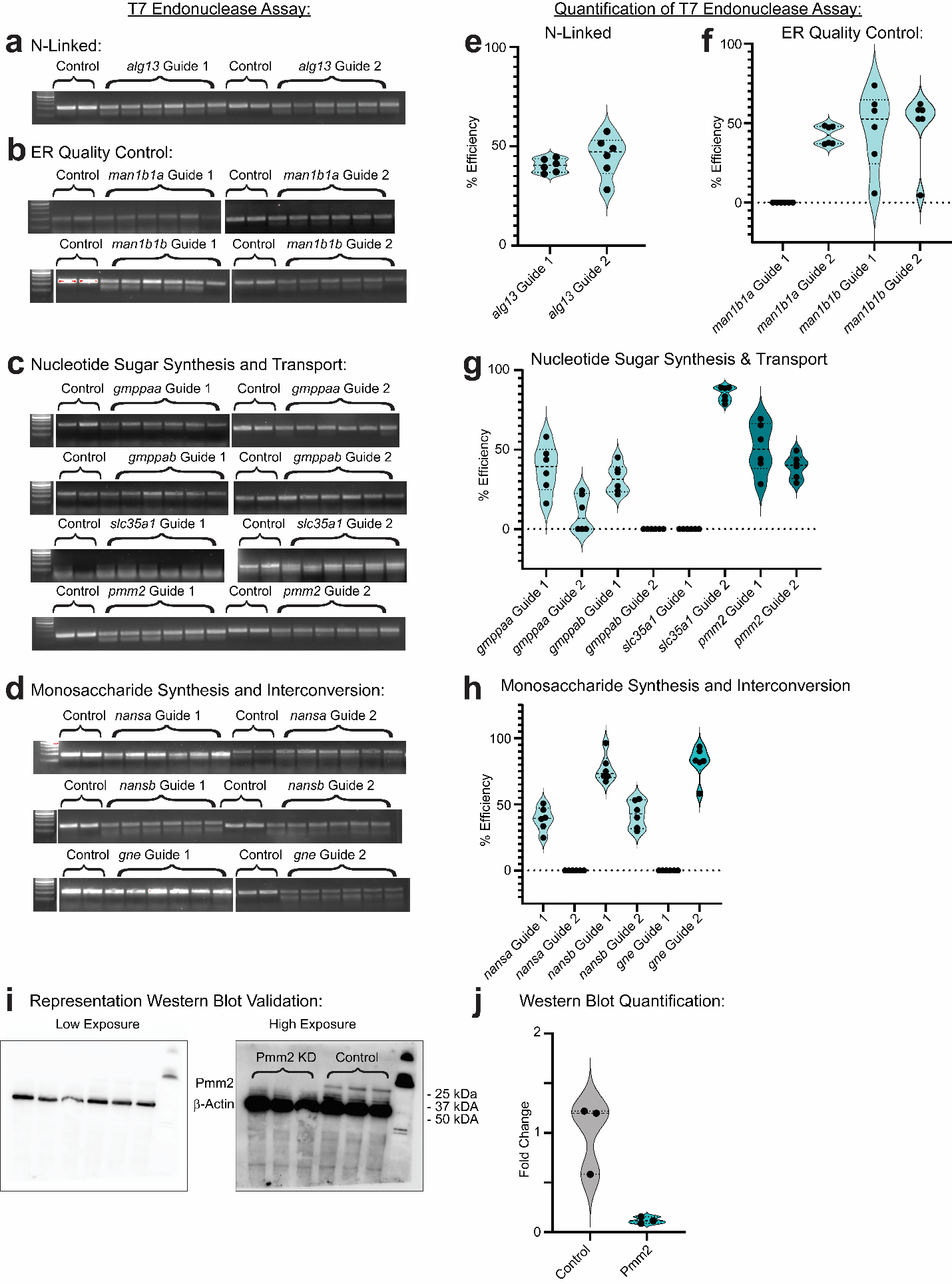

**Supplementary Figure 1: Knockdown efficiency of CRISPR guides targeting CDG-causing genes in the N-linked, ER quality control, nucleotide sugar synthesis and transport, and monosaccharide synthesis and interconversion classes.**

**a-d:** Gel electrophoresis images of T7 endonuclease assays for each CRISPR guide, separated by glycosylation gene class. (**a**) KD of *alg13;* member of the N-linked glycosylation pathway. (**b**) KD of *man1b1a* and paralog *man1b1b*; ER quality control gene. (**c**) From top to bottom, KD of paralogs *gmppaa* and *gmppab*, *slc35a1*, and *pmm2*; enzymes involved in nucleotide sugar synthesis and transport. (**d**) From top to bottom, KD of paralogs *nansa* and *nansb*, KD of *gne*; enzymes required for monosaccharide synthesis and interconversion. **e-h**: Quantification of T7 endonuclease assays, showing cutting efficiency of each guide. n=6 crispants and 2 controls per guide. Band intensity was quantified in FIJI/ImageJ and cutting efficiency was calculated as Σ(cleaved band intensity)/(uncleaved band intensity) * 100. **i:** Western blots showing protein KD in *pmm2* crispants*.* **j:** Quantification of western blot assay. Crispants (n=3 pools of 15 fish) were normalized to controls (n=3 pools of 15 fish). Band intensity was quantified in FIJI/ImageJ and normalized to the intensity of the corresponding housekeeping (Beta-Actin) band. Statistics by unpaired t-test (p = 0.0136). *p < .05, **p < .01, ***p < .001, ****p < .0001.

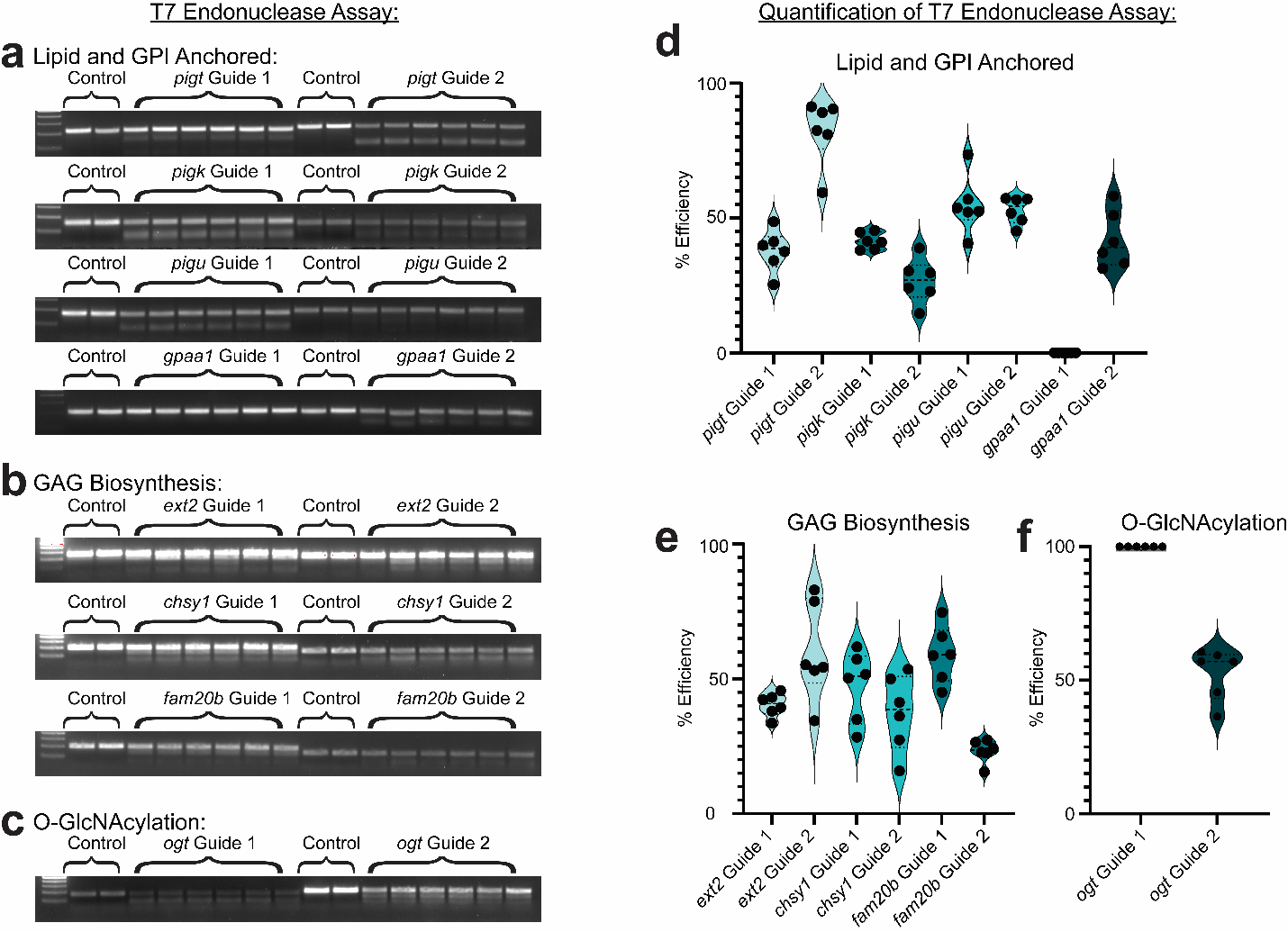
**Supplementary Figure 2: Knockdown efficiency of CRISPR guide RNAs targeting CDG-causing genes in the lipid and GPI-anchored, GAG biosynthesis, and O-GlcNAcylation classes.**

**a-c:** Gel electrophoresis images of T7 endonuclease assays for each CRISPR guide, separated by glycosylation gene class. (**a**) From top to bottom, KD of *pigt, pigk, pigu,* and *gpaa1*; enzymes required for glycosylation of lipid and GPI-anchored proteins. (**b**) From top to bottom, KD of *ext2*, *chsy1*, and *fam20b*; genes required for GAG biosynthesis. (**c**) KD of *ogt*, required for O-GlcNAcylation. **d-f**: Quantification of T7 endonuclease assays, showing cutting efficiency of each guide. n=6 crispants and 2 controls per guide. Band intensity was quantified in FIJI/ImageJ and cutting efficiency was calculated as Σ(cleaved band intensity)/(uncleaved band intensity) * 100.

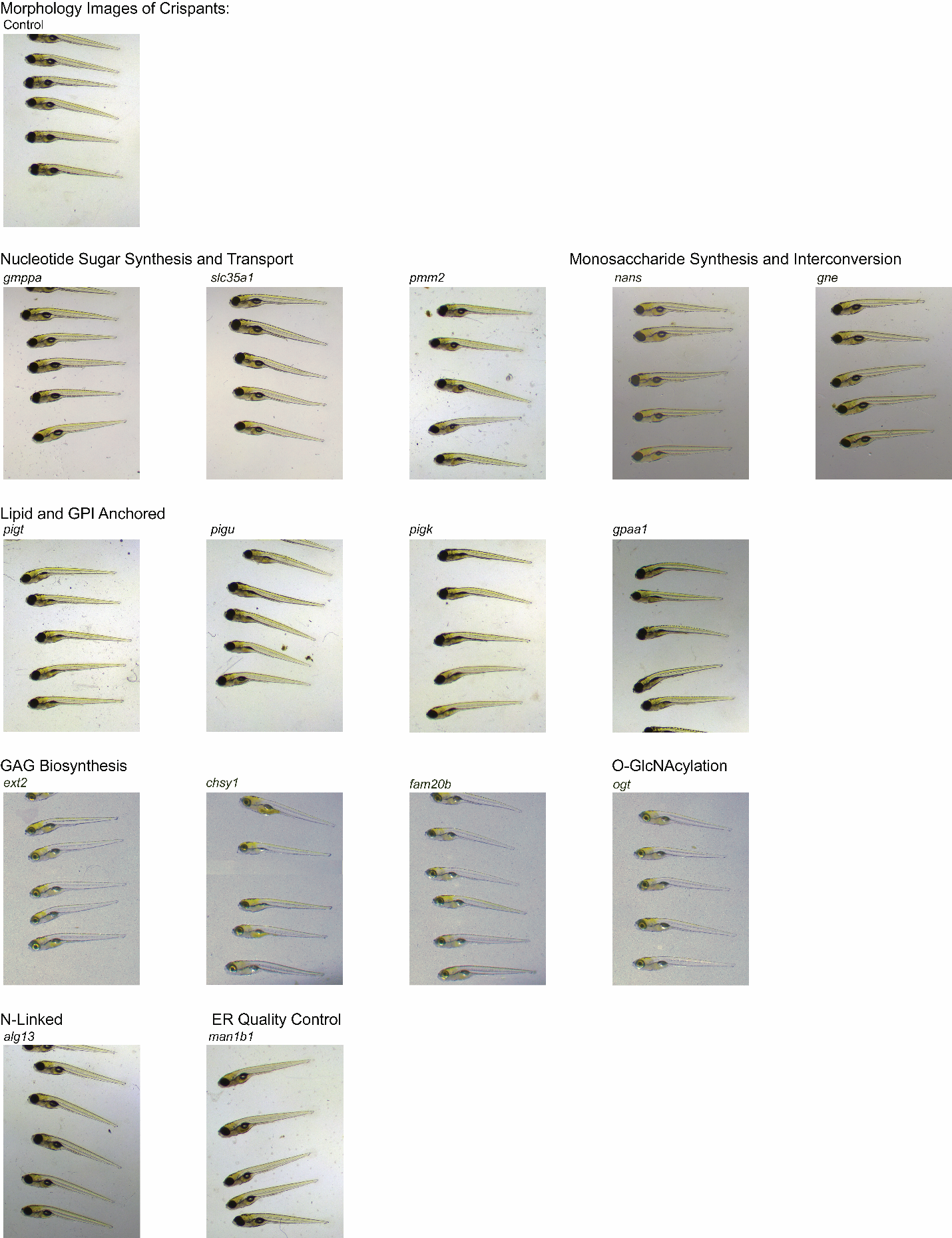

**Supplementary Figure 3: Crispant fish show minimal gross morphological changes.**

Representative whole-body images of crispant (n=5 per condition) and control (n=5) zebrafish, separated by glycosylation gene class. Fish were mounted in 1% low melt agar and imaged using bright-field microscopy.

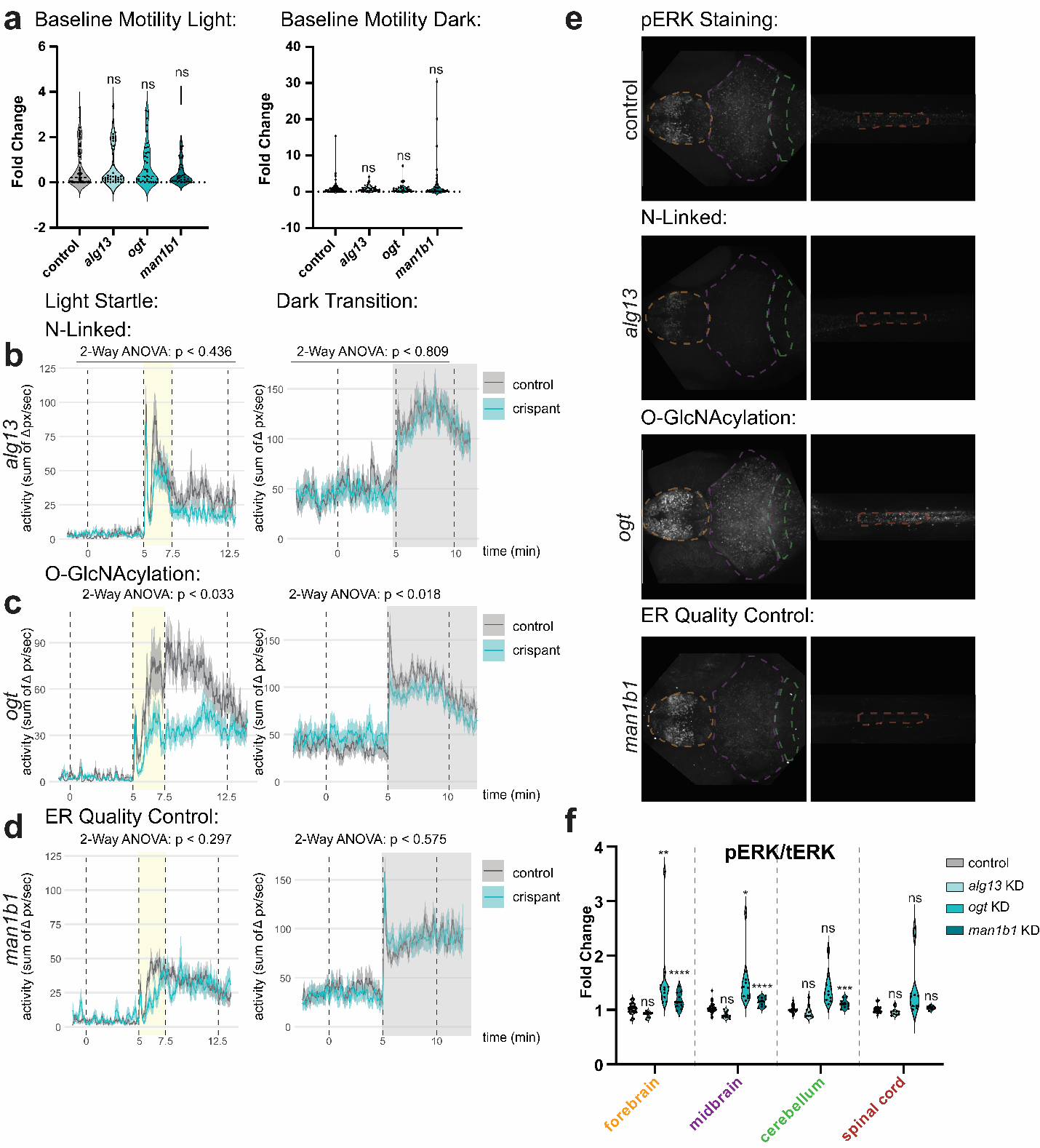

**Supplementary Figure 4: Knockdown of N-linked, O-GlcNAcylation, and ER quality control CDG genes lead to variable changes in zebrafish behavior and neuronal activity.**

**a:** Baseline motility of fish in the light (left) and dark (right). Activity level was averaged over 10 minutes, and crispants (n=36 per group) were normalized to controls (n=24 per group). Statistics by one-way ANOVA, multiple comparisons by Dunnett’s multiple comparisons test. P-values vs. control in light, left to right: 0.3649, 0.7140, 0.1117. P-values vs. control in dark, left to right: 0.8970, 0.9417, 0.2156. **b-e:** Activity traces showing response to the light startle assay (left) and dark transition assay (right) for (**b**) *alg13*, (**c**) *ogt*, and (**d**) *man1b1* crispants (n=36 per group) and controls (n=24 per group). Statistics by 2-way ANOVA. p-values shown on graphs represent time x genotype interaction. F-values for time x genotype interaction vs. control for light startle top to bottom: 0.7675, 3.631, 1.229. F-values for time x genotype interaction vs. control for dark transition top to bottom: 0.05952, 6.004, 0.3183). **f:** Representative images showing pERK staining in crispants and controls. From top to bottom: control, *alg13* KD, *ogt* KD, and *man1b1* KD. **g:** Quantification of pERK/tERK by region. Crispants (n=10 per group) were normalized to controls (n = 10 per group). Statistics by 2-way ANOVA, multiple comparisons by Sidak’s multiple comparisons test. P-values vs. control, left to right: forebrain: 0.4313, 0.0051, < 0.0001; midbrain: 0.2215, 0.0335, < 0.0001; cerebellum: 0.9887, 0.1305, 0.0005; spinal cord: 0.9966, 0.0893, 0.7351. *p < .05, **p < .01, ***p < .001, ****p < .0001.

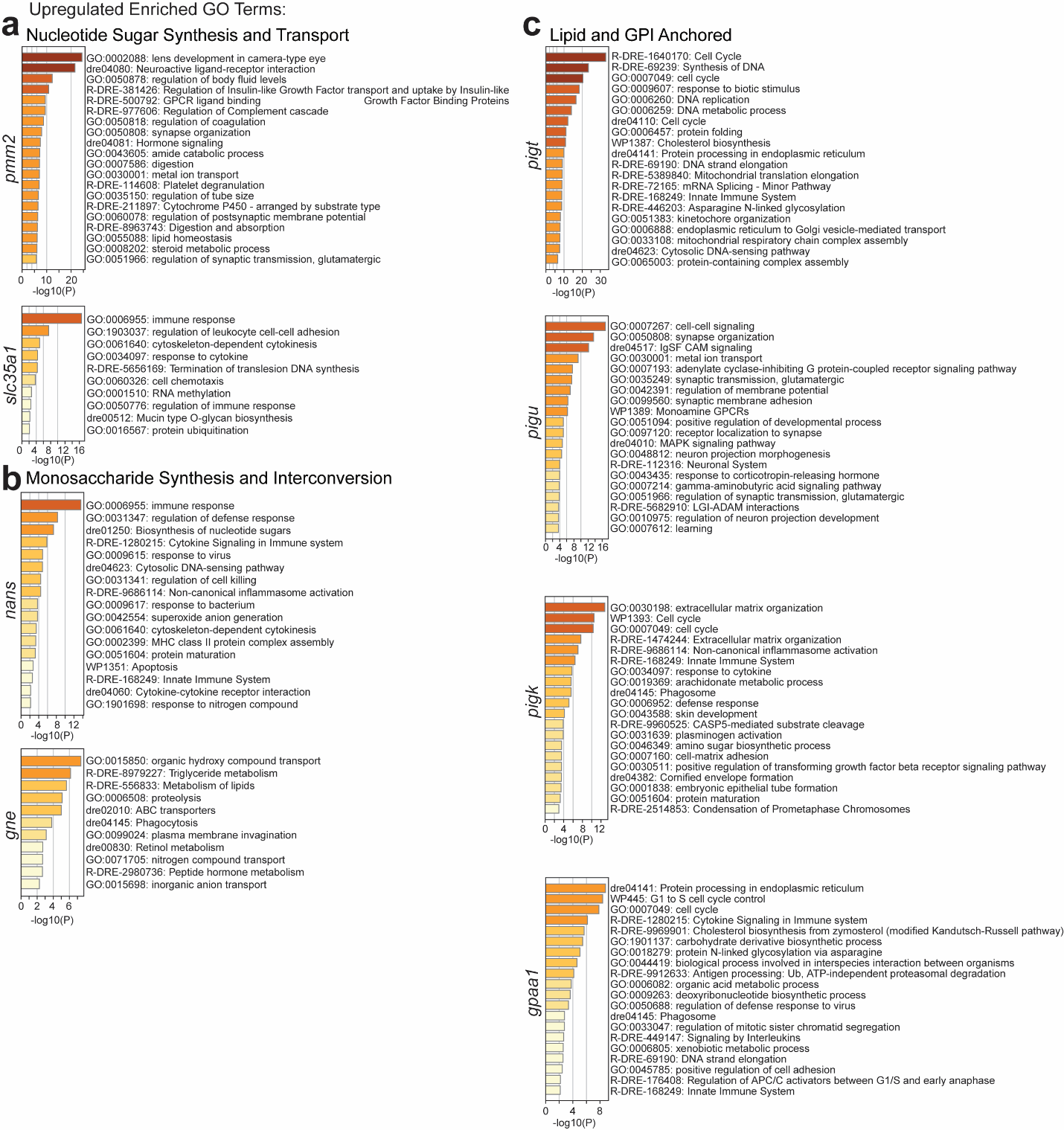

**Supplementary Figure 5: Upregulated GO-terms following knockdown of CDG-causing genes in the nucleotide sugar synthesis and transport, monosaccharide synthesis and interconversion, and lipid and GPI-anchored classes.**

GO-term analysis (Metascape) of significantly upregulated genes (adjusted p-value < .05; log2(fold change) ≥ 1) following KD of genes in (**a**) the nucleotide sugar synthesis and transport glycosylation pathway, (**b**) monosaccharide synthesis and interconversion glycosylation pathway, and (**c**) the lipid and GPI-anchored glycosylation pathway. n=3 pools of 20 fish per condition.

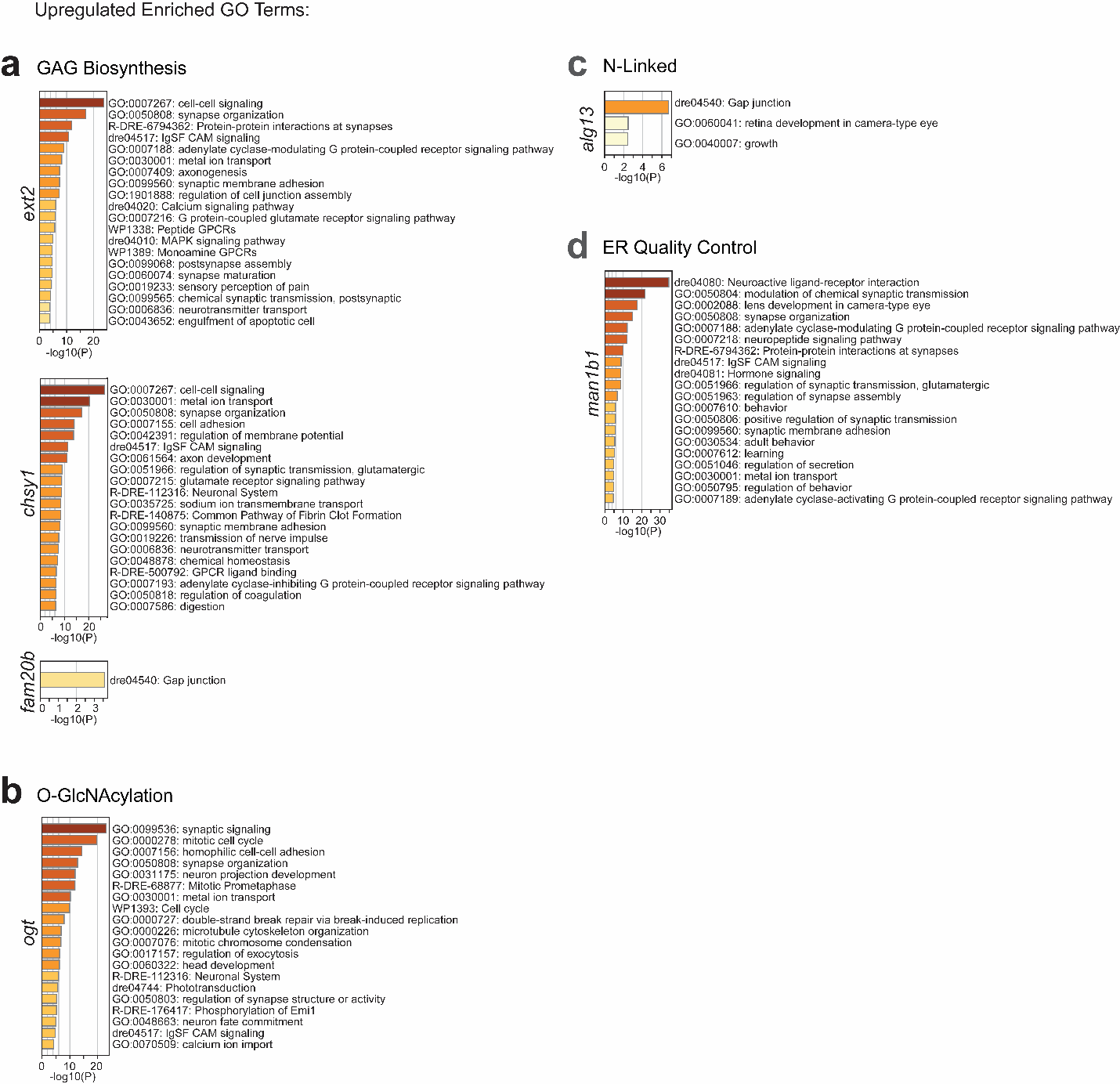

**Supplementary Figure 6: Upregulated GO-terms following knockdown of CDG-causing genes in the GAG biosynthesis, O-GlcNAcylation, N-linked, and ER quality control classes.**

GO-term analysis (Metascape) of significantly upregulated genes (adjusted p-value < .05; log2(fold change) ≥ 1) following KD of genes in the (**a**) GAG biosynthesis glycosylation pathway, (**b**) O-GlcNAcylation glycosylation pathway, (**c**) N-linked glycosylation pathway, and in (**d**) ER quality control. n=3 pools of 20 fish per condition.

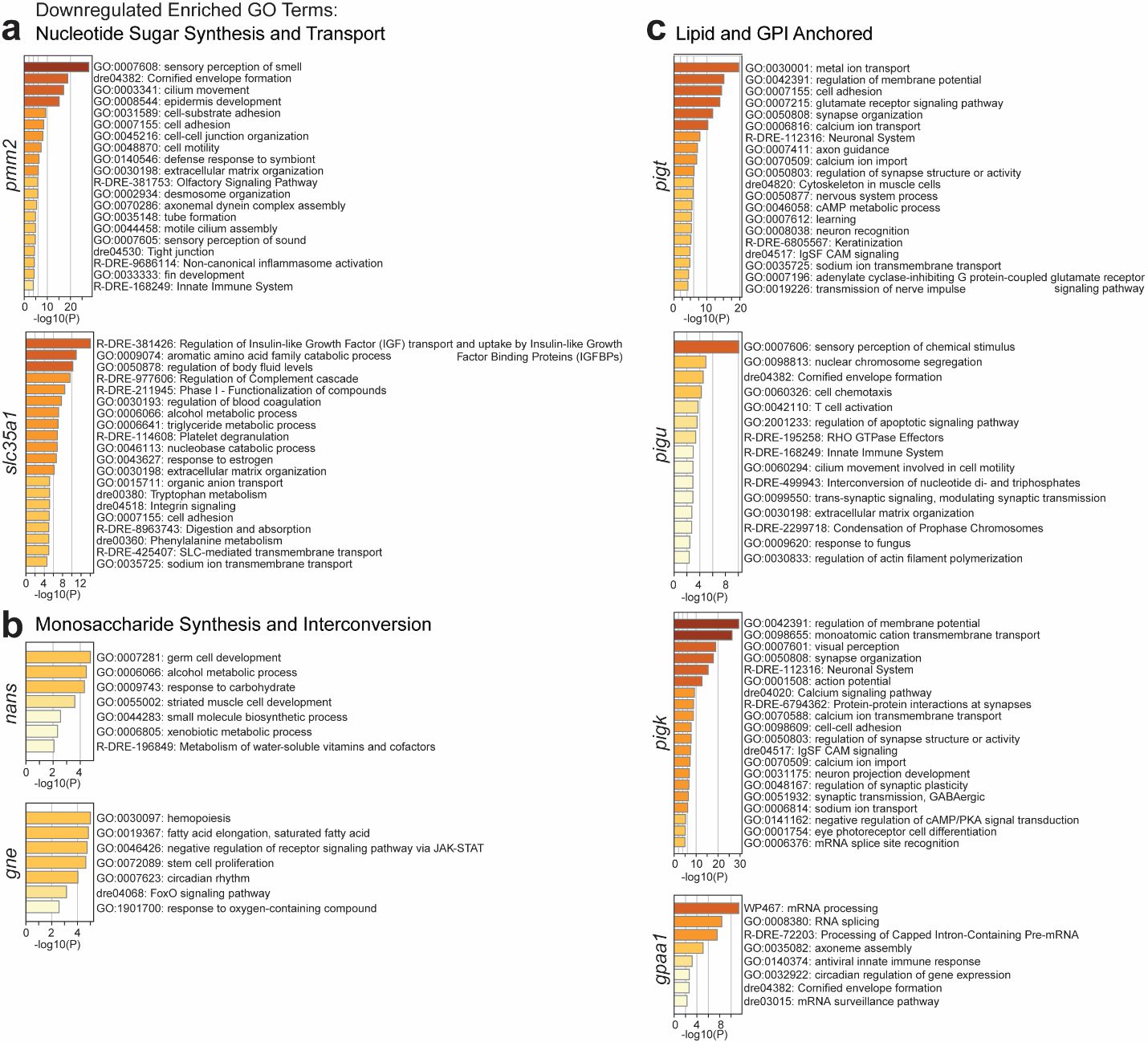

**Supplementary Figure 7: Downregulated GO-terms following knockdown of CDG-causing genes in the nucleotide sugar synthesis and transport, monosaccharide synthesis and interconversion, and lipid and GPI-anchored classes.**

GO-term analysis (Metascape) of significantly downregulated genes (adjusted p-value < .05; log2(fold change) ≤ -1) following KD of genes in the (**a**) nucleotide sugar synthesis and transport glycosylation pathway, (**b**) monosaccharide synthesis and interconversion glycosylation pathway, and (**c**) the lipid and GPI-anchored glycosylation pathway. n=3 pools of 20 fish per condition.

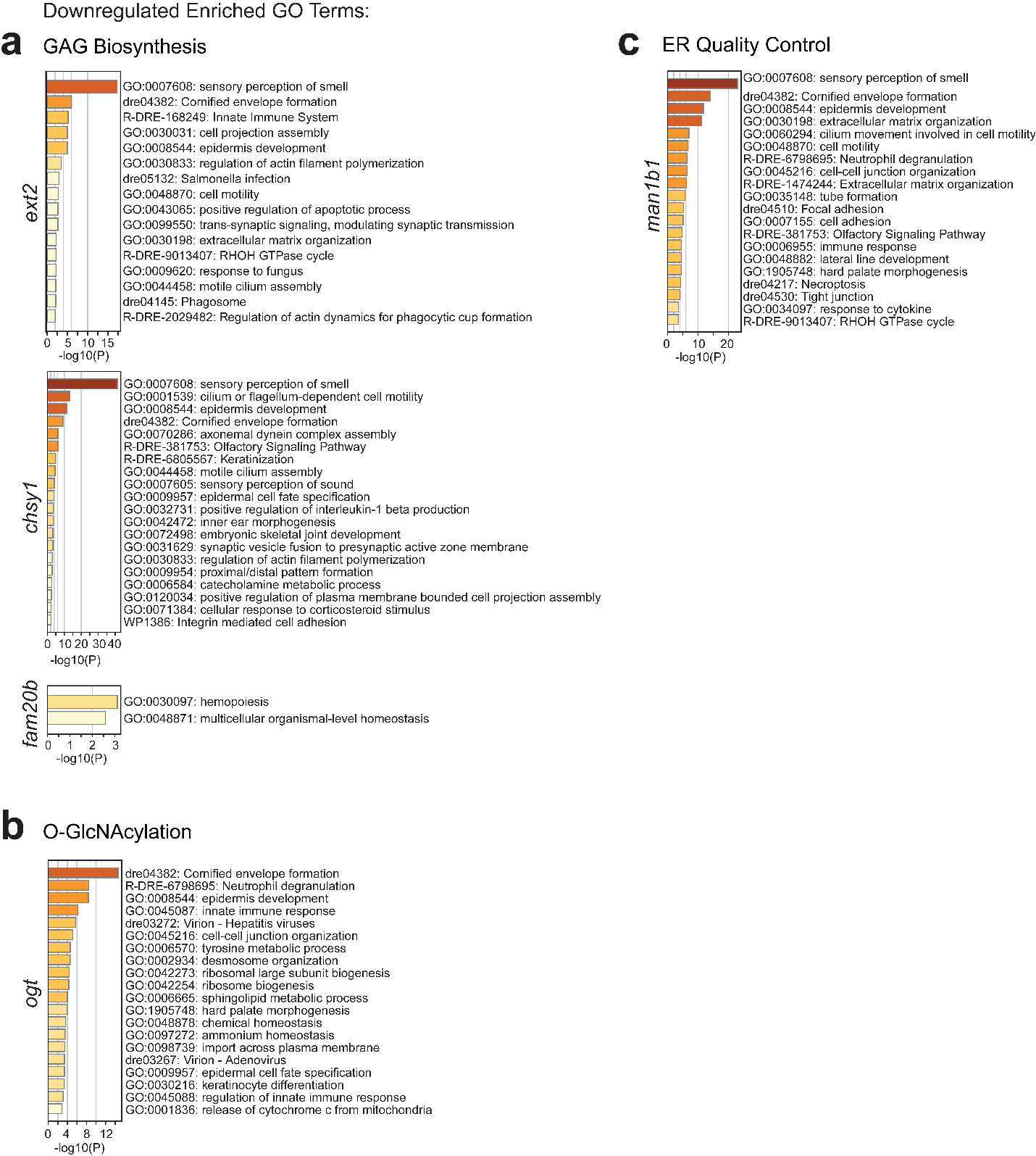

**Supplementary Figure 8: Downregulated GO-terms following knockdown of CDG-causing genes in the GAG biosynthesis, O-GlcNAcylation, N-linked and ER quality control classes.**

GO-term analysis (Metascape) of significantly downregulated genes (adjusted p-value < .05; log2(fold change) ≤ -1) following KD of genes in the (**a**) GAG biosynthesis glycosylation pathway, (**b**) O-GlcNAcylation glycosylation pathway, and in (**c**) ER quality control. n=3 pools of 20 fish per condition.

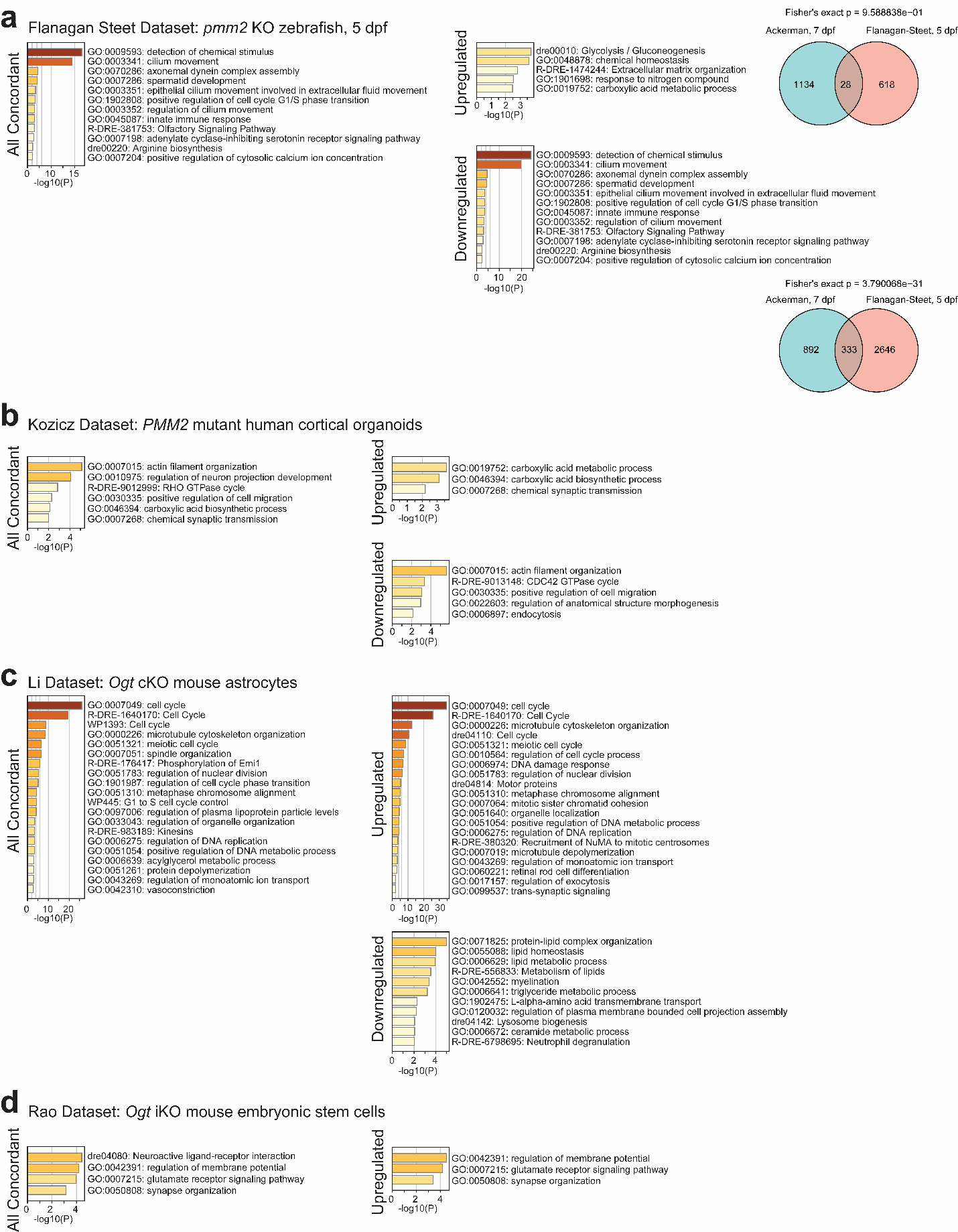

**Supplementary Figure 9: *pmm2* and *ogt* knockdown zebrafish show overlapping DEGs with published datasets.**

**a**: **Left:** GO-term analysis (Metascape) of all concordant significant (adjusted p-value < .05) DEGs in our *pmm2* KD zebrafish and the Flanagan-Steet lab’s *pmm2* KO zebrafish model. N=361 concordant genes. **Right:** GO-term analysis (Metascape of all significantly upregulated (log2(fold change) ≥ 1, top) and downregulated (log2(fold change) ≤ -1) genes shared in both datasets. Venn diagrams show the number of overlapping upregulated (top) and downregulated (bottom) DEGs. The number of overlapping downregulated DEGs is statistically significant (Fisher’s exact p = 3.790068e-31). Concordant genes are enriched for immune system function and neural function. **b**: **Left:** GO-term analysis (Metascape) of all concordant significant (adjusted p-value < .05) DEGs in our *pmm2* KD zebrafish and the Kozicz lab’s *PMM2* mutant human cortical organoids. N=61 concordant genes. **Right:** GO-term analysis (Metascape) of all significantly upregulated (log2(fold change) ≥ 1, top) and downregulated (log2(fold change) ≤ -1, bottom) genes shared in both datasets. Concordant genes are enriched for metabolism and neural development and function. **c**: **Left:** GO-term analysis (Metascape) of all concordant significant (adjusted p-value < .05) DEGs in our *ogt* KD zebrafish and the Li lab’s *Ogt* cKO mouse astrocytes. N=342 concordant genes. **Right:** GO-term analysis (Metascape) of all significantly upregulated (log2(fold change) ≥ 1, top) and downregulated (log2(fold change) ≤ -1, bottom) genes shared in both datasets. Concordant genes are enriched for cell cycle, metabolism, and synaptic signaling. **d**: **Left:** GO-term analysis (Metascape) of all concordant significant (adjusted p-value < .05) DEGs in our *ogt* KD zebrafish and the Rao lab’s *Ogt* cKO mouse embryonic stem cells. N=44 concordant genes. **Right:** GO-term analysis (Metascape) of all significantly upregulated (log2(fold change) ≥ 1) genes shared in both datasets. Concordant genes are enriched for neural development and function.

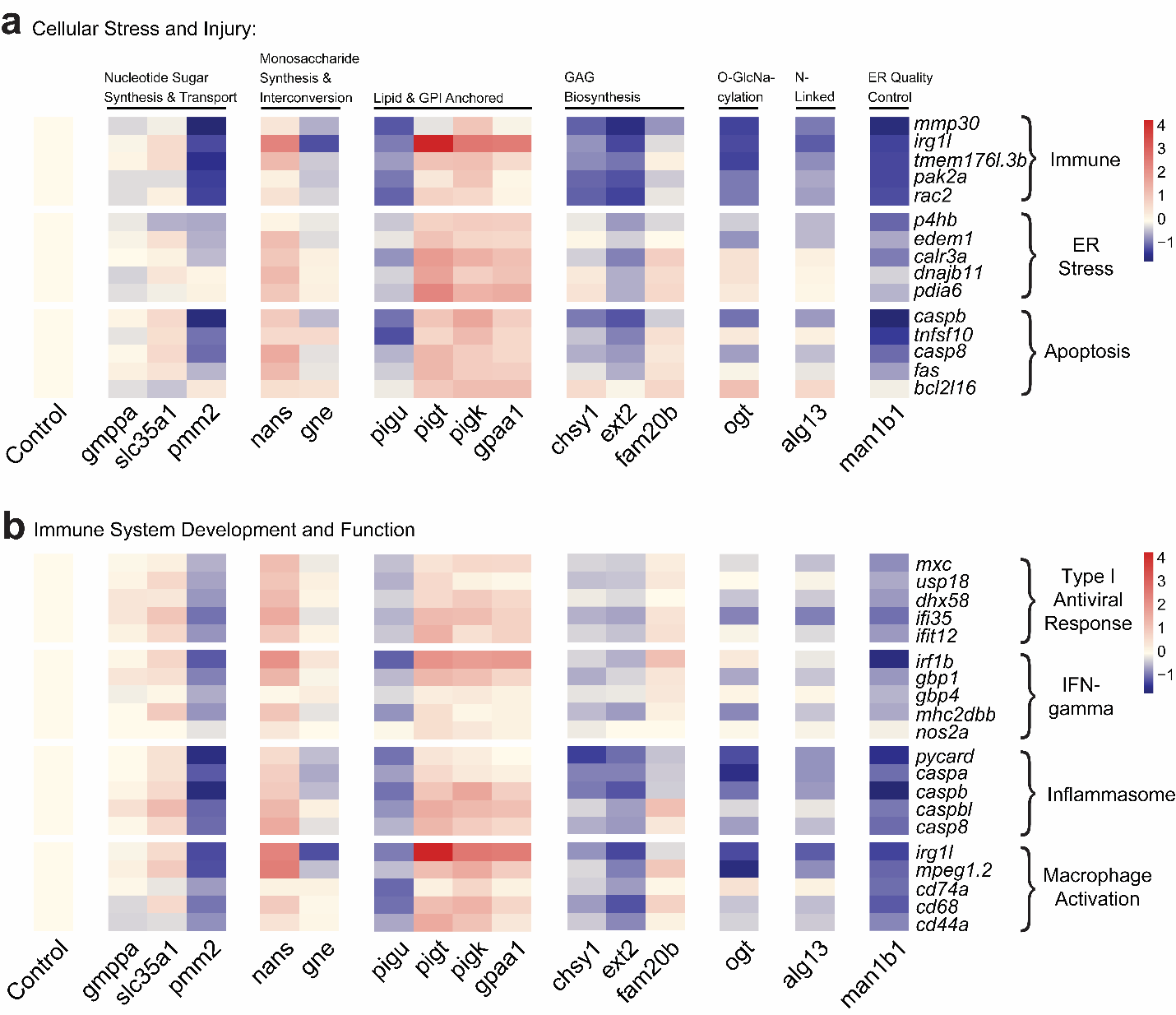

**Supplementary Figure 10: Genes related to cellular stress and injury and immune activation are dysregulated across all tested CDG classes.**

Heatmaps displaying log2(fold change) of selected genes related to cellular stress and injury and immune system function compared to control, colored to represent level of expression, and subdivided by glycosylation gene class. **a:** Genes related to immune system, ER stress, and apoptosis. **b:** Genes essential for specific categories of immune system activation and function including type 1 antiviral response, interferon-gamma, inflammasome, and macrophage activation.

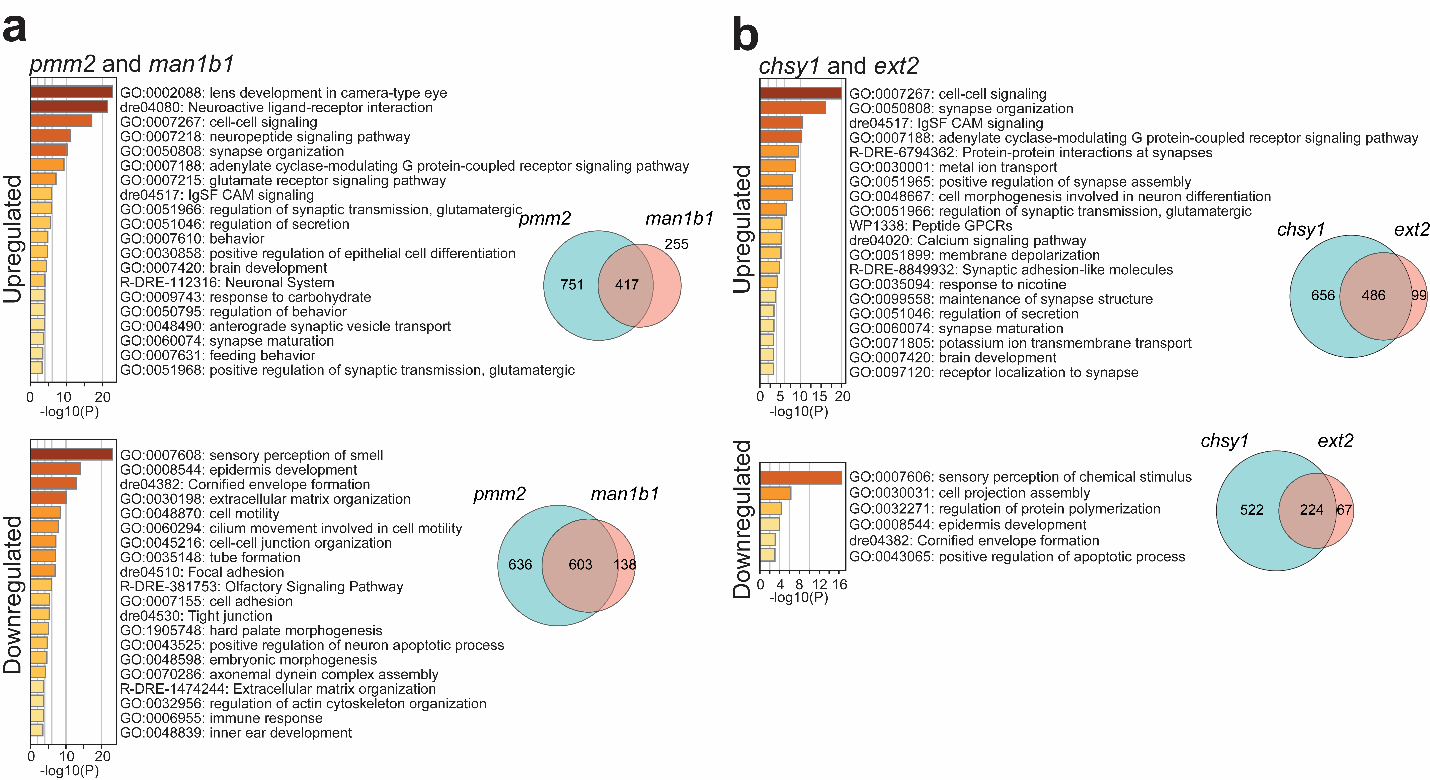

**Supplementary Figure 11: GO-term analysis of concordant genes between highly congruent pairs of knockdown zebrafish.**

GO-term analysis (Metascape) for pairs of knockdown zebrafish with Jacard Index > .4. **Left:** GO-term analyses of concordantly upregulated (adjusted p-value > .05, log2(fold change) ≥ 1, top) and downregulated (adjusted p-value > .05, log2(fold change) ≤ -1, bottom) genes in each pair. **Right:** Venn diagrams of concordantly upregulated (top) and downregulated (bottom) genes in each pair.

| **Gene Name** | **Guide 1 Sequence** | **Guide 2 Sequence** |
| --- | --- | --- |
| alg13 | TGTAAGCATGGACGAGTACC | GGAGGATTTTCCATACCCAG |
| chsy1 | GCTCCGGAAGAGCCATGACG | TCATTTGCACAATCTCACGA |
| ext2 | CCGTAAGCTGGACATCGTGA | TCAGGGAAGTACGGTTCACG |
| fam20b | CGCTACTGACGACACATGTG | AAGAGGCCCTGATAACACGT |
| gmppaa | CTTATCGGAGGCCCCCAAAA | CTTGTAAGGTAGCACCATGA |
| gmppab | ATCCATTCAGTACAGTAACG | AGAAGATCCAGTGAGTGCCA |
| gne | TGAACTACAGATAATCGCGT | CGTGGCCACTTGCAACCGAG |
| gpaa1 | GTCAAGGCTTTTCTGCGATT | TGCTGAGGGCTACACCCATG |
| man1b1a | GCGGTTATGGGAGAAACGCG | TTGGGATGAAGTGGATAATC |
| man1b1b | GACGACAAATCTGAGAAGCC | GTCACACTAACCTCAGAGGG |
| nansa | GAAATCGGACAGAATCATCA | TCCCGCTGGACACTACCATG |
| nansb | GAGATCGGACAGAACCATCA | AATACCGCATATCTTTACCT |
| ogt | TCCGCCATTTCGGTGTAGAC | AAATCCTACAACCCCTTACG |
| pigk | GTACCGGTGACGAACGATAA | TTCATCATAGACACGTGCCA |
| pigt | CCCAACCATGGCTTCTACGT | AGGCTACATTCACTACCAGC |
| pigu | CGACACCACCTCGACCCTCT | CGAGAACATCACCTGACGCA |
| pmm2 | GGCAGATGTTGTACACGTGG | CGTGTGAAAGTCGGAGTGGT |
| slc35a1 | GGTAAGCGACGACTGACAAG | TGACCAAGGCAGCTCGACAG |

**Supplementary Table 1: Sequences for CRISPR guide RNAs used in this study.**

Two CRISPR guides per gene were designed using CHOPCHOP. The sequence of each CRISPR guide is listed in this table.

| **Gene Name** | **Guide 1 Forward Primer Sequence** | **Guide 1 Reverse Primer Sequence** | **Product Size (Base Pairs)** |
| --- | --- | --- | --- |
| alg13 | GGAAAACCGTGAAGCTCTGTAT | CAACAATTTCGAGATTTGGTCA | 226 |
| chsy1 | CGTTAATTGCTCCTGACTACCC | TAAACCAAGATCTGCGACTCCT | 276 |
| ext2 | CTTTCCTGTCAGCTGTCCTTTT | CAATTCCAAAACACACTCGCTA | 212 |
| fam20b | CAGCATCAGATTCAGCAGTACC | AACAGGCTAACAGTGAAACCGT | 185 |
| gmppaa | CATCAAAACGATTCACTGAAGG | TTATGATGTCAATCCGAACTGC | 209 |
| gmppab | TAAACATCATCGTCCCAGAATG | CGGATTATTTCACGTCCAGTTT | 242 |
| gne | ATCAAGCTCATCTCATTTGGCT | TAAAGGTACATGTAACGTGGCG | 188 |
| gpaa1 | CTGTGGCTTTGTTTTCTGTTTG | AAGACGCCTGAACTAAATTGGA | 217 |
| man1b1a | AACATTGCACTGAACACACACA | CACTGAGTAGGTGACAGCAGGA | 197 |
| man1b1b | CTAACACGCCTCCTTACAGCTT | GATCACTCTCACTCCCGAAAAC | 199 |
| nansa | GTGACTGTCTGTTGACTCCTCG | CTCTCACCTTTGCCATTTTGAT | 207 |
| nansb | TAAAATGCCTGTCGAGTTTGAA | ATGGTTTTGGGATTTCCCTATT | 197 |
| ogt | TTTTGCTCAACTTGTCTCATGG | TAATGAAACATACCGATTTGCG | 178 |
| pigk | TACTCAAACCGGAAGTTTAGCG | CACTCACCTCAACATAATTGCTG | 168 |
| pigt | AGATCACGGTGCAGTTTGATAG | AAACTGCTAAACAATGGCTGGT | 234 |
| pigu | GCTCCTTTAACACTCCTTCTGC | CTGCAGGAACAAACCATCACTA | 183 |
| pmm2 | TACAATAACAACGTGGCCCATA | CTCAGTGGCAGTCAGTTCATTC | 197 |
| slc35a1 | GATAACCGACCATCATTACCGT | TCAGAACACACACAAAACCACA | 244 |
| **Gene Name** | **Guide 2 Forward Primer Sequence** | **Guide 2 Reverse Primer Sequence** | **Product Size (Base Pairs)** |
| alg13 | TGGTCAGCATGTCTCTTCTCAT | TAGGTAGAGTGAAGTGCCGGTT | 203 |
| chsy1 | CTTACCAGTATCGGTTGCACAG | AAAATGTACTTTCCCGTGAGGA | 226 |
| ext2 | TTCTGAGGGGCTATTTTGTGTC | AAGACCACAGAGAAGAGGGTGA | 156 |
| fam20b | AATGTGATTGACTCTTGATGCG | GGCCAAAAGTAAAAAGGGAGAC | 252 |
| gmppaa | CAGTTAGGTCCGAATGTGTCAA | GGGTTTAAATGGCAGACAAAAA | 161 |
| gmppab | GACATCATCAACTGTGGCATCT | ATCTGTCCAGTGAGAAGTGCAA | 151 |
| gne | ACTATCTGAGAATGCAGCGAGC | TTCTGCTCAAGCTTTCTTACCC | 222 |
| gpaa1 | CCGTATCACCCAGCTCTAGTCT | TAGTGCCTGAAACTGTTGATGC | 215 |
| man1b1a | TCTGTCTGTTTGTTCTCCAGGT | TAACATCTGGTTGGACGTCTTG | 149 |
| man1b1b | TGTGTTTGTGTGTCTTGGTTGA | TTTTGGAGGGAAAACAGACATT | 252 |
| nansa | CAAGACTGTAGAGCCCCAAAAT | GATGCAAAAGTTCTGATTGTGC | 265 |
| nansb | GCTCCAACCCTTGTTTTATCAT | ATGGTTTTGGGATTTCCCTATT | 150 |
| ogt | TGCTGCCTCAAGTCTGTTACAT | CAAGACTTCCCAATGACATGAA | 200 |
| pigk | TCAGTTTCCTAAAAGCAGCCTC | TTACTGAAAGTGAGTCCTCCCC | 159 |
| pigt | AACCAAACACACACACCAACAA | AGGGGTGTATTCAGTCCATTTG | 255 |
| pigu | TTGAACACAGCCACTGAAAGTT | TCAGTCCAGTGAAAGAAGGTCA | 202 |
| pmm2 | TTTTTGTCTTCAGAAAGCCACC | CCACTCTGTCAATCACTGGAAA | 218 |
| slc35a1 | GTGTGTGTGATCTGCTCTGTGA | TGTGTGTGTGTGTGTGATGATG | 264 |

**Supplementary Table 2: Sequences for primers used in T7 endonuclease assays.**

One primer pair was designed around the expected cut site of each CRISPR guide used in the study using NCBI primer blast. The forward primer and reverse primer sequences, as well as the expected product size in base pairs, are listed in this table.

| **Pool name** | **Sequence** |
| --- | --- |
| alg13_b1 | ACAATCTCTTGGAAGGTCGCTGCCATAGAAGAGTCTTCCTTTACG |
| alg13_b1 | GAGGAGGGCAGCAAACGGAAGGGGTCATAAGAGTAGGGTGGAGGA |
| alg13_b1 | TTGGGCTGTAAGGAGTTTGGCGGGGTAGAAGAGTCTTCCTTTACG |
| alg13_b1 | GAGGAGGGCAGCAAACGGAATGGGAGCTCCCAGTGGATGAAGAAG |
| alg13_b1 | GGCAGCCGGGGCGGAGTTGATTGGTTAGAAGAGTCTTCCTTTACG |
| alg13_b1 | GAGGAGGGCAGCAAACGGAAAGAAGAGGAGAGGACAGATGACTTC |
| alg13_b1 | AACTGTAGGTAGCGACACCCTCTTGTAGAAGAGTCTTCCTTTACG |
| alg13_b1 | GAGGAGGGCAGCAAACGGAAGCGACCCCGCACGAGAACTCCCTTC |
| alg13_b1 | GTAGATTCTGCTGCTGCTGCATACATAGAAGAGTCTTCCTTTACG |
| alg13_b1 | GAGGAGGGCAGCAAACGGAAGATGGACTCTGATAACCTGCTTCTG |
| alg13_b1 | CTCAGAGTATTCACCTTGATCACCATAGAAGAGTCTTCCTTTACG |
| alg13_b1 | GAGGAGGGCAGCAAACGGAAAGCACTCCTCTCATCAGGGTCCTCC |
| alg13_b1 | GTTTGCCTGAGGGAGGAATTGGACTTAGAAGAGTCTTCCTTTACG |
| alg13_b1 | GAGGAGGGCAGCAAACGGAAGAACTCTTGGCACCCAGTAAGCAGT |
| alg13_b1 | CTAGAGACTGCCTGTCCCGATATTGTAGAAGAGTCTTCCTTTACG |
| alg13_b1 | GAGGAGGGCAGCAAACGGAAGAGAAGGCACCCTCATCAGTCTCTT |
| alg13_b1 | GCTTCCCTCCATGTCTTGGGTCTCCTAGAAGAGTCTTCCTTTACG |
| alg13_b1 | GAGGAGGGCAGCAAACGGAATCCTGGCTCAGGAACACAGCTTAGA |
| alg13_b1 | GCATCTGCCTTCGACTCCTACGAGATAGAAGAGTCTTCCTTTACG |
| alg13_b1 | GAGGAGGGCAGCAAACGGAATGAATGAGTAGTTATCCAAACTCTG |
| alg13_b1 | CTAGGATAATAGGCCATGTCTGGTCTAGAAGAGTCTTCCTTTACG |
| alg13_b1 | GAGGAGGGCAGCAAACGGAAATAAGATGGTGCCTGTTCAGGAATC |
| alg13_b1 | ATACCCACGCGGCTGTCTAGGTGGATAGAAGAGTCTTCCTTTACG |
| alg13_b1 | GAGGAGGGCAGCAAACGGAAGTGGGGTCGGTAGTGTTCATAAATC |
| alg13_b1 | CTGGACTGTGCACCCCACGAGGCTTTAGAAGAGTCTTCCTTTACG |
| alg13_b1 | GAGGAGGGCAGCAAACGGAATGAAGAACCGTCGCCGTCCTCGAAG |
| alg13_b1 | GGGTACTGCTCTGCATGGCTGTATGTAGAAGAGTCTTCCTTTACG |
| alg13_b1 | GAGGAGGGCAGCAAACGGAATTTCCCTTTCTGCTTGGTGTTATGT |
| alg13_b1 | CACTGGCTTCAGATTGGTCAAAGAATAGAAGAGTCTTCCTTTACG |
| alg13_b1 | GAGGAGGGCAGCAAACGGAAAAGGTGCTTTTCTCCAAGCTCCTCA |
| alg13_b1 | TGGAATGAGTCCCAACTTCTTGAATTAGAAGAGTCTTCCTTTACG |
| alg13_b1 | GAGGAGGGCAGCAAACGGAAAGGCATTGTAGTACTTTCCTTTTGG |
| alg13_b1 | TTATCTCCTAAAAAGTACTGTCTGCTAGAAGAGTCTTCCTTTACG |
| alg13_b1 | GAGGAGGGCAGCAAACGGAAGCAAACACCATGTAATCGGTCTTCT |
| alg13_b1 | TTGCCACACGTCAAACTCCACGTTGTAGAAGAGTCTTCCTTTACG |
| alg13_b1 | GAGGAGGGCAGCAAACGGAAGTACAGCTCTGGCTCTAAGGCCTTC |
| alg13_b1 | CTAGTTTCCCAGACCCTTCTGTGACTAGAAGAGTCTTCCTTTACG |
| alg13_b1 | GAGGAGGGCAGCAAACGGAATTTGGTCCTCAGCTCCAACTCTAGG |
| alg13_b1 | TGTCCCCTCGACTCCCAATCCTCTCTAGAAGAGTCTTCCTTTACG |
| alg13_b1 | GAGGAGGGCAGCAAACGGAAAAAAAGTCACTCTGAGAACGATCTT |
| alg13_b1 | AACTGTGAGTGTGGCAGATGGTTGGTAGAAGAGTCTTCCTTTACG |
| alg13_b1 | GTCTTCGCTGCACATAGAATGGCTGTAGAAGAGTCTTCCTTTACG |
| alg13_b1 | GAGGAGGGCAGCAAACGGAATCGGTATCGTCGTCCCCCTCCTTTG |
| alg13_b1 | GAGGAGGGCAGCAAACGGAATGACCCGGGGCTGTGATATAGTTGA |
| alg13_b1 | GTATTTCATCATCCTCAACTCCCAGTAGAAGAGTCTTCCTTTACG |
| alg13_b1 | GAGGAGGGCAGCAAACGGAACACGAGTGTACAACAGTTCATACAG |
| alg13_b1 | GCATCTACTGGATAGCTCTTTGGGTTAGAAGAGTCTTCCTTTACG |
| alg13_b1 | GAGGAGGGCAGCAAACGGAAACAATGTCATAGTGGCCATTGTTTG |
| alg13_b1 | AGGCAAATTGTCTTCCTCTGCAATCTAGAAGAGTCTTCCTTTACG |
| alg13_b1 | GAGGAGGGCAGCAAACGGAAAGTTGGTGGCTTACCTGGATACCTG |
| alg13_b1 | TGTACAGTAAAGACAGTGCCTTTATTAGAAGAGTCTTCCTTTACG |
| alg13_b1 | GAGGAGGGCAGCAAACGGAATCACTTGACCTACGGTTTCCTTGGG |
| alg13_b1 | TACTTCTCAAAAGATCCTTCAACAATAGAAGAGTCTTCCTTTACG |
| alg13_b1 | GAGGAGGGCAGCAAACGGAAGGTTCAAAGTTGCATCTGTTCGCCC |
| alg13_b1 | CTTTCGTATTCTTTGATGGTAATTCTAGAAGAGTCTTCCTTTACG |
| alg13_b1 | GAGGAGGGCAGCAAACGGAAAGAGTAATAAAGCTGCTCAGAAACA |
| alg13_b1 | CATCTCTCGCCATCATTTTTCGGTATAGAAGAGTCTTCCTTTACG |
| alg13_b1 | GAGGAGGGCAGCAAACGGAAGTCCAATGCTGGCCAGGTACTCGTC |
| alg13_b1 | GTCTAAAAGCGAAAAAACTGTGAAGTAGAAGAGTCTTCCTTTACG |
| alg13_b1 | GAGGAGGGCAGCAAACGGAACGAAAGTTTCCTTTCGGGCTTGAAA |
| alg13_b1 | GGGACAGTAAGTGGTGTGGTGGAACTAGAAGAGTCTTCCTTTACG |
| alg13_b1 | GAGGAGGGCAGCAAACGGAAGGAATGTGGGGCAGCACCAGGAGTC |
| alg13_b1 | ATGTTGTGGAGAGCACAGGCATGTATAGAAGAGTCTTCCTTTACG |
| alg13_b1 | GAGGAGGGCAGCAAACGGAAGGTCATAGGTAGAGTGAAGTGCCGG |
| alg13_b1 | ACCACAGGATAGAACACCGTGCCAGTAGAAGAGTCTTCCTTTACG |
| alg13_b1 | GAGGAGGGCAGCAAACGGAAACCACTGTAGGAGGATTTTCCATAC |
| alg13_b1 | GGGCATGTCTCCTTGTGGGTGCGGGTAGAAGAGTCTTCCTTTACG |
| alg13_b1 | GAGGAGGGCAGCAAACGGAACATGCGGACAGAATCAGGGTAGGCC |
| alg13_b1 | CGGGAGGAGCTGTGCTCTGGTAAGATAGAAGAGTCTTCCTTTACG |
| alg13_b1 | GAGGAGGGCAGCAAACGGAAGGAATTGAGCATCTGCAGCTTGATG |
| alg13_b1 | TGCACCATAGAGTGCCAGTAACTCTTAGAAGAGTCTTCCTTTACG |
| alg13_b1 | GAGGAGGGCAGCAAACGGAATGATACCACTGAACACCCAAATTAT |
| chsy1_B2 | CATCGCTCTCTGTAAACATTCACAAAAATCATCCAGTAAACCGCC |
| chsy1_B2 | CCTCGTAAATCCTCATCAAAAAACATTTACCATCTTGCCAGACAC |
| chsy1_B2 | TTTTTTTCTGGCGAGGTCCACAGCAAAATCATCCAGTAAACCGCC |
| chsy1_B2 | CCTCGTAAATCCTCATCAAACTGTTTTTGAGGTGCCATAAATATA |
| chsy1_B2 | CCTCATTGATTCATTATTGGAGTTCAAATCATCCAGTAAACCGCC |
| chsy1_B2 | CCTCGTAAATCCTCATCAAAGATCTTTCGGAAGCTGTGGTTGTTT |
| chsy1_B2 | GCTTTGGATCCCAGGCACATTTTGTAAATCATCCAGTAAACCGCC |
| chsy1_B2 | CCTCGTAAATCCTCATCAAATGCTTGGGATCAAGATTGGGGTCGC |
| chsy1_B2 | ACAGCTTGATCCCGGATTGGACAAAAAATCATCCAGTAAACCGCC |
| chsy1_B2 | CCTCGTAAATCCTCATCAAATGTTGAAAAGGTCAACGTCCTCCAG |
| chsy1_B2 | TCCTTTGTAGACGCAGACAATACCAAAATCATCCAGTAAACCGCC |
| chsy1_B2 | CCTCGTAAATCCTCATCAAAACCATAGTGTCTCCACAGGCCTGTT |
| chsy1_B2 | ACAACCTTTGGGTCATACTGGCTGAAAATCATCCAGTAAACCGCC |
| chsy1_B2 | CCTCGTAAATCCTCATCAAAATAATAGGAAAGTATGTCTGTTCCC |
| chsy1_B2 | GTAAGTCCACGTCACAGTAGAAGAGAAATCATCCAGTAAACCGCC |
| chsy1_B2 | CCTCGTAAATCCTCATCAAAAAGAATCATTGGTAAAGTGGGCCGA |
| chsy1_B2 | GATCTCCATATCGGCTTTGGGGTACAAATCATCCAGTAAACCGCC |
| chsy1_B2 | CCTCGTAAATCCTCATCAAACATGCGATACTCCCTCATGAGCTCG |
| chsy1_B2 | TGTATGGGGGCTAACAAAGAACTGAAAATCATCCAGTAAACCGCC |
| chsy1_B2 | AGAAGCTTGACATTCTGGTTGGGAAAAATCATCCAGTAAACCGCC |
| chsy1_B2 | CCTCGTAAATCCTCATCAAAAGGCAGATCTTCTCGAAGTTGGCCA |
| chsy1_B2 | CCTCGTAAATCCTCATCAAATATGAAAACGCCCCAGAGAAAAACA |
| chsy1_B2 | TGATCTTCTTCTCTTTGGGTTCGTGAAATCATCCAGTAAACCGCC |
| chsy1_B2 | CCTCGTAAATCCTCATCAAAGTTCGATGCCTGGGCTGCTCAACTT |
| chsy1_B2 | GGAGTCTTGGTTGATCCTGGCAGCCAAATCATCCAGTAAACCGCC |
| chsy1_B2 | CCTCGTAAATCCTCATCAAAGACTCGAGCGTCCATCTCTTCTTCT |
| chsy1_B2 | CTTACGGGCACTGTCATAGTCTTCCAAATCATCCAGTAAACCGCC |
| chsy1_B2 | CCTCGTAAATCCTCATCAAATTGTGCTTCTTGTAGAGCAGCAGCA |
| chsy1_B2 | CATATTGGATCTCTTTGAAGTCAATAAATCATCCAGTAAACCGCC |
| chsy1_B2 | CCTCGTAAATCCTCATCAAACTCTTCCGCGTGTCTTGGCATTGGC |
| chsy1_B2 | CTGCGAGGAGTCAATGCCTCGACGGAAATCATCCAGTAAACCGCC |
| chsy1_B2 | CCTCGTAAATCCTCATCAAAAGGCTGTCCATCTGAGGATGAGAAA |
| chsy1_B2 | TGGGGATGAAAGCGCATGAAGGATGAAATCATCCAGTAAACCGCC |
| chsy1_B2 | CCTCGTAAATCCTCATCAAAGGCATGCCCAGTTGAAGGTCTTCCC |
| chsy1_B2 | GGAGCTGAATGGTGCGGTGGCGCAGAAATCATCCAGTAAACCGCC |
| chsy1_B2 | CCTCGTAAATCCTCATCAAACTGCAATCTTCCGGCTCAACATGTA |
| chsy1_B2 | TATGGCTCTGTGGATCTTACTGTTGAAATCATCCAGTAAACCGCC |
| chsy1_B2 | CCTCGTAAATCCTCATCAAAGAGATCACGGATGTAGCCCTTCTTG |
| chsy1_B2 | TCCTGCGCAGCACCTCGCGGCTCATAAATCATCCAGTAAACCGCC |
| chsy1_B2 | CCTCGTAAATCCTCATCAAATAACACCAGGGCCGCCCATGCAGAA |
| chsy1_B2 | CATGCCCGTCTGGCCCAGAAAGATGAAATCATCCAGTAAACCGCC |
| chsy1_B2 | CCTCGTAAATCCTCATCAAACTCGCTGCTGTTGAGACTCCTTAGG |
| chsy1_B2 | ATGAACCACTCGAACTGGTCCAAATAAATCATCCAGTAAACCGCC |
| chsy1_B2 | CCTCGTAAATCCTCATCAAATGGTCATGCATGTATTTGAGCATCA |
| chsy1_B2 | CCACAATGGGGATGGGGATGGTGGTAAATCATCCAGTAAACCGCC |
| chsy1_B2 | CCTCGTAAATCCTCATCAAACAGAACCTTCACTGGAGAAAAACTC |
| chsy1_B2 | GCGGTTATTCAGGTACTTCTGCGCGAAATCATCCAGTAAACCGCC |
| chsy1_B2 | CCTCGTAAATCCTCATCAAACATGACGCCCACAAACAGAAAGCTG |
| chsy1_B2 | AGCTACTAGTCGCTTTAATTTGTCAAAATCATCCAGTAAACCGCC |
| chsy1_B2 | CCTCGTAAATCCTCATCAAAGTTTTGTTTATTAAACGCCGCCCGT |
| chsy1_B2 | GGTCACCGCTCCGGTCCCCATAAATAAATCATCCAGTAAACCGCC |
| chsy1_B2 | CCTCGTAAATCCTCATCAAACCAACCGCTGCCATCTAATCACACA |
| chsy1_B2 | GATTTCGAGCCCGAGTCGGTGGGTAAAATCATCCAGTAAACCGCC |
| chsy1_B2 | CCTCGTAAATCCTCATCAAACAGGAGCAATTAACGGTTTGTTGCG |
| chsy1_B2 | CTACAGTACATTACATGTTTTTTCCAAATCATCCAGTAAACCGCC |
| chsy1_B2 | CCTCGTAAATCCTCATCAAACTCTACTGCCTTGCATTTGGACTCA |
| chsy1_B2 | AAAAACGAACCGAAAAGAAGCTGTTAAATCATCCAGTAAACCGCC |
| chsy1_B2 | CCTCGTAAATCCTCATCAAATCAATGGCCCTTAATTGTCAGAATT |
| ext2_b3 | ACTGTTTACCAGTGGATTGTGATACTTCCACTCAACTTTAACCCG |
| ext2_b3 | GTCCCTGCCTCTATATCTTTCTCAGAGTGTTCACTTTTCTCACAC |
| ext2_b3 | GCTTGTGACCAGTCCTCATAGAAATTTCCACTCAACTTTAACCCG |
| ext2_b3 | GTCCCTGCCTCTATATCTTTTCCACTGCTTTGATGCGTCGCAACT |
| ext2_b3 | AGAGAAATCCCGGAGTGCCCGTAGCTTCCACTCAACTTTAACCCG |
| ext2_b3 | GTCCCTGCCTCTATATCTTTGTCTGAGAGAAAAAAAATAAGTGGT |
| ext2_b3 | CAATGTTGGGGAAACTTTTGAGTTTTTCCACTCAACTTTAACCCG |
| ext2_b3 | GTCCCTGCCTCTATATCTTTCTGGGAAGTCGTCTTTGTAGAGTAC |
| ext2_b3 | ATGGTGCCGAACACAGAGGCGAACTTTCCACTCAACTTTAACCCG |
| ext2_b3 | GTCCCTGCCTCTATATCTTTTTTATGCACTCAGACCTCTCCACCA |
| ext2_b3 | TGTGCACTCCGGGCATTTAAATTTCTTCCACTCAACTTTAACCCG |
| ext2_b3 | GTCCCTGCCTCTATATCTTTCCGAGGAGTAACCTTTATTGGTGCT |
| ext2_b3 | CAATATCCTCACAGTTCATGTGAGCTTCCACTCAACTTTAACCCG |
| ext2_b3 | GTCCCTGCCTCTATATCTTTCCACCCAGTTCTTGATGTCTCCAGG |
| ext2_b3 | TGGTAGAAAGCAGCACCAGTCAAGATTCCACTCAACTTTAACCCG |
| ext2_b3 | GTCCCTGCCTCTATATCTTTATGGATACTTCATTAGTCCATTCAG |
| ext2_b3 | TAAATGGAGCCGACCAGGATATCCATTCCACTCAACTTTAACCCG |
| ext2_b3 | GTCCCTGCCTCTATATCTTTCAGCCTGTCAGGGAACTCACGCCAT |
| ext2_b3 | ACATGATGATGTCATCGTCAATGGCTTCCACTCAACTTTAACCCG |
| ext2_b3 | GTCCCTGCCTCTATATCTTTGGACAGCTTCTGTCTCGATCTCATC |
| ext2_b3 | AGAAAAGGATGATTGCTTGTATCACTTCCACTCAACTTTAACCCG |
| ext2_b3 | TTGGTGCGGACCACTTTAAGAGGCATTCCACTCAACTTTAACCCG |
| ext2_b3 | GTCCCTGCCTCTATATCTTTGCGACTTTGGGCCACAGCGACTCTT |
| ext2_b3 | GTCCCTGCCTCTATATCTTTAGGAGAAGAGTCACTTGTTGAAACC |
| ext2_b3 | CAGTAGTTTGGCCAAACTGGGCACCTTCCACTCAACTTTAACCCG |
| ext2_b3 | GTCCCTGCCTCTATATCTTTGGAGATCTCAGTGATGACCCTAAAA |
| ext2_b3 | CTGTGAACCCAGGAGAGCGTGGCGGTTCCACTCAACTTTAACCCG |
| ext2_b3 | GTCCCTGCCTCTATATCTTTTGAGAGGAAGGAAAAGTGGACTGTT |
| ext2_b3 | CATTCTTCATAGGTATGAGCTGCATTTCCACTCAACTTTAACCCG |
| ext2_b3 | GTCCCTGCCTCTATATCTTTGGGTAAATGCGATCATTAATGATCT |
| ext2_b3 | GAAGTAGGCTTCCCAAAACCAGCGATTCCACTCAACTTTAACCCG |
| ext2_b3 | GTCCCTGCCTCTATATCTTTCTGTCTCTGCATCTCCTCCACTTGT |
| ext2_b3 | CTGGCAACTTTTCTTCTGGAATGACTTCCACTCAACTTTAACCCG |
| ext2_b3 | GTCCCTGCCTCTATATCTTTCAGACGCCCTTTTCCAGTCCAGCAC |
| ext2_b3 | ATGACAGGGACACATCCCGCTTGCATTCCACTCAACTTTAACCCG |
| ext2_b3 | GTCCCTGCCTCTATATCTTTACGTCACTGAGCGTGGCTTGGCCTA |
| ext2_b3 | CTCCTGTAAAATTTGTGGGTAGTCGTTCCACTCAACTTTAACCCG |
| ext2_b3 | GTCCCTGCCTCTATATCTTTCACCTGTCCTTTATAGCAGCGCTTT |
| ext2_b3 | ATTTGTCTAACAGGAGCAGGGCCTCTTCCACTCAACTTTAACCCG |
| ext2_b3 | GTCCCTGCCTCTATATCTTTCATTCTCATCCTTCAGCCTTTCCAG |
| ext2_b3 | GATGACAGAATGAAGTAGCGCCGAGTTCCACTCAACTTTAACCCG |
| ext2_b3 | GTCCCTGCCTCTATATCTTTCCGGGTTGCCTCTCAGGAAGGTCCA |
| ext2_b3 | GACATCATAACCTTGTCTGTAGGTCTTCCACTCAACTTTAACCCG |
| ext2_b3 | GTCCCTGCCTCTATATCTTTTGTGGAGAAGCCTCCCCCAGCCAGC |
| ext2_b3 | TGTAGTCAGGAGGTCCTCCAGGAAGTTCCACTCAACTTTAACCCG |
| ext2_b3 | GTCCCTGCCTCTATATCTTTTATTAAACAGCAGGTGGTTCATGCC |
| ext2_b3 | GCAGTTTCTCGAATACGCAGAGAGTTTCCACTCAACTTTAACCCG |
| ext2_b3 | GTCCCTGCCTCTATATCTTTTGATTCAGCACATCAATAGAAGGTA |
| ext2_b3 | GAAGTCACTGTCCGAGATGGCGCTCTTCCACTCAACTTTAACCCG |
| ext2_b3 | GTCCCTGCCTCTATATCTTTCAGGTCATTGTATTCTCGAGACAGG |
| ext2_b3 | ACTCCATTCGGCTGAAGACTCAATGTTCCACTCAACTTTAACCCG |
| ext2_b3 | GTCCCTGCCTCTATATCTTTGTGAGGCCAGAACTGGAACATCCCT |
| ext2_b3 | GGGTGATGTAGTAGATCCGGTGTTTTTCCACTCAACTTTAACCCG |
| ext2_b3 | GTCCCTGCCTCTATATCTTTTCTTCATGCGGGGTATGAGAGCAGG |
| ext2_b3 | ATGAGCCTGTCCAGACAAAACCACTTTCCACTCAACTTTAACCCG |
| ext2_b3 | GTCCCTGCCTCTATATCTTTACGATGTCCAGCTTACGGCCTCAAA |
| fam20b_b2 | AAAGAAGCCTGCGCATCAAGAGTCAAAATCATCCAGTAAACCGCC |
| fam20b_b2 | CCTCGTAAATCCTCATCAAACACATTTGTACTTCCCTGGTTTGTC |
| fam20b_b2 | TACGGATGAGGCAGGTTCATCCGGTAAATCATCCAGTAAACCGCC |
| fam20b_b2 | CCTCGTAAATCCTCATCAAATCGATCAGTGTGTTTTCCGAGCCCT |
| fam20b_b2 | CACTGCCGGACGGTGGCTAATACTCAAATCATCCAGTAAACCGCC |
| fam20b_b2 | CCTCGTAAATCCTCATCAAATTGAGGCGCCGGTCGAGCGCAGTGA |
| fam20b_b2 | CCTTTCAGGAGGTTCAGTCTGTTCCAAATCATCCAGTAAACCGCC |
| fam20b_b2 | CCTCGTAAATCCTCATCAAAGTGGACACGCGGACCATGCAGCACT |
| fam20b_b2 | GCGAGAATACTCCTCTCATCCAAAGAAATCATCCAGTAAACCGCC |
| fam20b_b2 | CCTCGTAAATCCTCATCAAAGGGTTCCCAAAACTTTTGGCATTAT |
| fam20b_b2 | AGCATACTTGCTCCGCCATCGTCCTAAATCATCCAGTAAACCGCC |
| fam20b_b2 | CCTCGTAAATCCTCATCAAAAAGCTCTCGTAGTGATGGCGGTCAG |
| fam20b_b2 | AGATAATCGAAGATGGAAGTGTCGAAAATCATCCAGTAAACCGCC |
| fam20b_b2 | CCTCGTAAATCCTCATCAAAACATCCAGCAGTCGCGGCCCAGCAT |
| fam20b_b2 | ATTTTCTTTACAGCCTCGCAGTAACAAATCATCCAGTAAACCGCC |
| fam20b_b2 | CCTCGTAAATCCTCATCAAATCATCATACTCCCACCTGGCGAGTT |
| fam20b_b2 | TATGTGCGTCCCCATGGGTGTCTGTAAATCATCCAGTAAACCGCC |
| fam20b_b2 | CCTCGTAAATCCTCATCAAATTCTGTAAGGGCCAGACGTCAGGGA |
| fam20b_b2 | TTTACCATCTTCTCTGAACATCAGTAAATCATCCAGTAAACCGCC |
| fam20b_b2 | TTGCCATAGAAGCATGTATTATTCCAAATCATCCAGTAAACCGCC |
| fam20b_b2 | CCTCGTAAATCCTCATCAAATGCATCAGGAAAGTGCTCAGCAGTT |
| fam20b_b2 | CCTCGTAAATCCTCATCAAATAACATCCGTACAACTCTATATCAT |
| fam20b_b2 | ACTGGTTTTATCTCCGTGCGCAGATAAATCATCCAGTAAACCGCC |
| fam20b_b2 | CCTCGTAAATCCTCATCAAAATAAACCTCCCCACCACCAGAGGCG |
| fam20b_b2 | CCCAGAATCCTGTCCAAGTGGAAAGAAATCATCCAGTAAACCGCC |
| fam20b_b2 | CCTCGTAAATCCTCATCAAAGCGATTTCTGCATTATGGCGATCGT |
| fam20b_b2 | GGCTCTCCCTCCACCACATAGTCTCAAATCATCCAGTAAACCGCC |
| fam20b_b2 | CCTCGTAAATCCTCATCAAAACGTACCTCTTGGGCTTAAACACAA |
| fam20b_b2 | CCGTCGAGTACCAGCAGGGCTTTTAAAATCATCCAGTAAACCGCC |
| fam20b_b2 | CCTCGTAAATCCTCATCAAATGGGTTCCTTTATATCCCACGTCTG |
| fam20b_b2 | TGAGCGGTGGCCAGAGCATTCAGCAAAATCATCCAGTAAACCGCC |
| fam20b_b2 | CCTCGTAAATCCTCATCAAAGCAGCTAACTCAGGTGTTTCGTCTG |
| fam20b_b2 | CCTTTGGCCAGGGAAAGGCTGGCCTAAATCATCCAGTAAACCGCC |
| fam20b_b2 | CCTCGTAAATCCTCATCAAAATCTTTCGGAAGGCCCGGAGGTCGC |
| fam20b_b2 | GCTGATCCCTCTCCTCCATCCAGTAAAATCATCCAGTAAACCGCC |
| fam20b_b2 | CCTCGTAAATCCTCATCAAAAAGACCTTGGCCAGGCCGAGGAGGA |
| fam20b_b2 | CAGAGCACCACCACACGCTGCTTCAAAATCATCCAGTAAACCGCC |
| fam20b_b2 | CCTCGTAAATCCTCATCAAATTCATCACCTCCCACCACCACACCG |
| fam20b_b2 | TATGCCTGGGACATGGCACTGCCCGAAATCATCCAGTAAACCGCC |
| fam20b_b2 | CCTCGTAAATCCTCATCAAAGTCTCACCAACACCAGCACCACCCG |
| fam20b_b2 | TGGACACACTTCCTTAAATCCTCCTAAATCATCCAGTAAACCGCC |
| fam20b_b2 | CCTCGTAAATCCTCATCAAATGTTTGATTTGGAGTTCAGCCTGGT |
| fam20b_b2 | TCTGCTGCTACTGTCTGCACAAGTCAAATCATCCAGTAAACCGCC |
| fam20b_b2 | CCTCGTAAATCCTCATCAAAGCATCTCCCAGCAGATTTGATCTCC |
| fam20b_b2 | ATTAATATCCACCCGCCTGTCGGTCAAATCATCCAGTAAACCGCC |
| fam20b_b2 | CCTCGTAAATCCTCATCAAACCTCCATTCATTACTGGTACTGCTG |
| fam20b_b2 | TTGTTAATCCATCATGGTCACTTTGAAATCATCCAGTAAACCGCC |
| fam20b_b2 | CCTCGTAAATCCTCATCAAAGTTATCAGTCTAATAAGGCCAAATG |
| fam20b_b2 | GTCTCAGAATTGTCCATCCATCTTCAAATCATCCAGTAAACCGCC |
| fam20b_b2 | CCTCGTAAATCCTCATCAAAGCGGATGGCCAAAAGTAAAAAGGGA |
| fam20b_b2 | CGACAGTATCTGCTGTAGTGTTATAAAATCATCCAGTAAACCGCC |
| fam20b_b2 | CCTCGTAAATCCTCATCAAACCACCTACGTGTTATCAGGGCCTCT |
| fam20b_b2 | GAAAAAAATCGACTCCCAAGGCTTTAAATCATCCAGTAAACCGCC |
| fam20b_b2 | CCTCGTAAATCCTCATCAAAATGGCCTTCGGGTCTTCGGCTCTAT |
| gmppaa_B3 | GCACTCTGCTCAGCTTTCATACAGGTTCCACTCAACTTTAACCCG |
| gmppaa_B3 | CCAACACGGCTGGTCTGCAATTACATTCCACTCAACTTTAACCCG |
| gmppaa_B3 | TTGCCCCGAGCCAACAGCAAATAAATTCCACTCAACTTTAACCCG |
| gmppaa_B3 | TCTTTGTGCGGCAGGACGATAGAGTTTCCACTCAACTTTAACCCG |
| gmppaa_B3 | GTCCCTGCCTCTATATCTTTAGTATGATGACCTCAGACGGGATGT |
| gmppaa_B3 | GTCCCTGCCTCTATATCTTTCTTTCCATCTCTGAAGAGCGTCTCG |
| gmppaa_B3 | TGGCATATGGGTCATTAGGATTGGGTTCCACTCAACTTTAACCCG |
| gmppaa_B3 | ATGGTGCTTTCCCAACCCACAATGCTTCCACTCAACTTTAACCCG |
| gmppaa_B3 | GTCCCTGCCTCTATATCTTTTTCAGCACGCAGCTGTGATCTTGTA |
| gmppaa_B3 | ACACATTCGGACCTAACACAGCAGTTTCCACTCAACTTTAACCCG |
| gmppaa_B3 | GTCCCTGCCTCTATATCTTTCAGCCGTTAGACTGCTCCTCTAACA |
| gmppaa_B3 | GTCCCTGCCTCTATATCTTTGTGCTGGAAAATCTCCGGAGTAAAC |
| gmppaa_B3 | GAGGATGACAAAGCTGTAGGTGTCTTTCCACTCAACTTTAACCCG |
| gmppaa_B3 | GTCCCTGCCTCTATATCTTTATGCTCCTTCTGGAAGTCCAGCATC |
| gmppaa_B3 | GAAACTCTGAGCACACGTCAGCATTTTCCACTCAACTTTAACCCG |
| gmppaa_B3 | GTCCCTGCCTCTATATCTTTCCAGTTCCCAGAGCAGCATATTCCT |
| gmppaa_B3 | GTCCCTGCCTCTATATCTTTGGAAACAGAGGTTTGGGCACTTCAA |
| gmppaa_B3 | GTCCCTGCCTCTATATCTTTTCCGATAAGAATGACAGCCTTCAGC |
| gmppaa_B3 | GTCCCTGCCTCTATATCTTTACCAACAGGCTGTAAAATGCTGCTG |
| gmppaa_B3 | GTCCCTGCCTCTATATCTTTCACATCTGGTAAATCCATCTGCACT |
| gmppaa_B3 | GCCCAAACTACTCACAATAGTGGAATTCCACTCAACTTTAACCCG |
| gmppaa_B3 | ATGAAGGATGATAGACTCCCTCACATTCCACTCAACTTTAACCCG |
| gmppaa_B3 | GTCCCTGCCTCTATATCTTTAACTCCTGCACCGATCGTTACGCCT |
| gmppaa_B3 | GTCCCTGCCTCTATATCTTTGATCAATGTTGGCAGTGGGATGTAT |
| gmppaa_B3 | GTCCCTGCCTCTATATCTTTTTAGTAGCTAACCTCTCTGGATGTG |
| gmppaa_B3 | GTCCCTGCCTCTATATCTTTAATCGCAGATCCAGCAGATTTGATC |
| gmppaa_B3 | TCCAGACGGATCACCTCTGCTCGATTTCCACTCAACTTTAACCCG |
| gmppaa_B3 | CTGGTTCTTCTGGAAAACTGATCCATTCCACTCAACTTTAACCCG |
| gmppaa_B3 | CACAGTTGATGATGTCGCTCACAAATTCCACTCAACTTTAACCCG |
| gmppaa_B3 | GTCCCTGCCTCTATATCTTTTGCCAGGCTTCTCTACATAATGCAA |
| gmppaa_B3 | GTCCCTGCCTCTATATCTTTTGACAAAAAATGCATCTGGACCTCC |
| gmppaa_B3 | GTCCCTGCCTCTATATCTTTTCATATTTGGAAGCTTAGAGCAGGC |
| gmppaa_B3 | GCCTAGAATTGTAATGGAAGGAGTTTTCCACTCAACTTTAACCCG |
| gmppaa_B3 | CCTCTGGTCTTTGCTCCTCCCTCTGTTCCACTCAACTTTAACCCG |
| gmppaa_B3 | TTGATTCAGATACAAACGGCTTGCATTCCACTCAACTTTAACCCG |
| gmppaa_B3 | ATCGGTCTGTTTTGTACACGTAAAGTTCCACTCAACTTTAACCCG |
| gmppaa_B3 | GTTTGCTCATTCTCAACAATACAGCTTCCACTCAACTTTAACCCG |
| gmppaa_B3 | GTCCCTGCCTCTATATCTTTTAATTCAGGGACTGTTTTCTGTTGG |
| gmppaa_B3 | TGATCTCTGAAATGGTAGATTCCTCTTCCACTCAACTTTAACCCG |
| gmppaa_B3 | GTCCCTGCCTCTATATCTTTCTTCAGTGAATCGTTTTGATGCATG |
| gmppaa_B3 | GTCCCTGCCTCTATATCTTTGCAGTCACAGAAACATCAATTATTC |
| gmppaa_B3 | TCTTTTTATCCAGCTTTGGCATCATTTCCACTCAACTTTAACCCG |
| gmppaa_B3 | GTCCCTGCCTCTATATCTTTACAGTATGATCTGGTTCTTAAAGCT |
| gmppaa_B3 | GTCCCTGCCTCTATATCTTTCACTGGGAGTTCCCTCCACTCTGGC |
| gmppaa_B3 | GTCCCTGCCTCTATATCTTTTGCCCTGCCCTGCCAGAGCAGTGAA |
| gmppaa_B3 | GATAAAACCCAATGAGTAAGATCTCTTCCACTCAACTTTAACCCG |
| gmppaa_B3 | TGCTGCAGCATTGGCACTCCGGCCATTCCACTCAACTTTAACCCG |
| gmppaa_B3 | CCGAAAGCGGGTCCCTTTTTGGGGGTTCCACTCAACTTTAACCCG |
| gmppaa_B3 | AACAAGCTGAAATAAGTGTCTGAGATTCCACTCAACTTTAACCCG |
| gmppab_b1 | GAGGAGGGCAGCAAACGGAAGTCTCACTGTCTATCTTGGCATACG |
| gmppab_b1 | TTAGGGTCACTTGGAGTTCCTTCCATAGAAGAGTCTTCCTTTACG |
| gmppab_b1 | GAGGAGGGCAGCAAACGGAATAGATGCCACAGTTGATGATGTCGC |
| gmppab_b1 | TGACGGTGGAACTGGAGCATTTCTATAGAAGAGTCTTCCTTTACG |
| gmppab_b1 | GAGGAGGGCAGCAAACGGAAAAGGGGAACTCTGAGCATACATCTG |
| gmppab_b1 | ATGAAGCGGTTCAGCTCATCGTTGGTAGAAGAGTCTTCCTTTACG |
| gmppab_b1 | GGAACTTGGGCACAGGCTTCAATGTTAGAAGAGTCTTCCTTTACG |
| gmppab_b1 | GAGGAGGGCAGCAAACGGAATGTTGCAGCATTGGAACTCCTGCTA |
| gmppab_b1 | GAGGAGGGCAGCAAACGGAACGGAACCTGGTACCTTTTTGAGGAC |
| gmppab_b1 | TGTGTCAGTGCGTGTAAGGCTGGTGTAGAAGAGTCTTCCTTTACG |
| gmppab_b1 | GAGGAGGGCAGCAAACGGAAGGAATATTCACATTGCAGCCAAGGA |
| gmppab_b1 | ACTCTTACACCTCCTCCAATTGTCATAGAAGAGTCTTCCTTTACG |
| gmppab_b1 | GAGGAGGGCAGCAAACGGAACCTTTGCCAATGGAAACATTAGGAC |
| gmppab_b1 | TGGTTTGTGGCCAGTCTCTCCGGATTAGAAGAGTCTTCCTTTACG |
| gmppab_b1 | TTTTTCTGTCCTGCTAGGGCTGTAATAGAAGAGTCTTCCTTTACG |
| gmppab_b1 | GAGGAGGGCAGCAAACGGAAATGTCCTGTTCCAGCCGAACCACCT |
| gmppab_b1 | CTGCCATGAGTGAGCTCTTCCTGGATAGAAGAGTCTTCCTTTACG |
| gmppab_b1 | TGGGCAAAAATATCTGGGGTGAACATAGAAGAGTCTTCCTTTACG |
| gmppab_b1 | CTGGGTTTCTCTACAAAATGGAGCATAGAAGAGTCTTCCTTTACG |
| gmppab_b1 | GAGGAGGGCAGCAAACGGAAGTTCCAAGTAGGACTCCACAATGAT |
| gmppab_b1 | GGTTTGGGCACCTCAAATGACAGGGTAGAAGAGTCTTCCTTTACG |
| gmppab_b1 | GAGGAGGGCAGCAAACGGAATTCCGAAAATCGCAACAGCGAGCCC |
| gmppab_b1 | GAGGAGGGCAGCAAACGGAAAAATCCATTCAGTACAGTAACGCGG |
| gmppab_b1 | ATAGAGTTTCGAATGATGACCTCTGTAGAAGAGTCTTCCTTTACG |
| gmppab_b1 | GAGGAGGGCAGCAAACGGAACTTGCCCACTTCCCCACTGTGCTGT |
| gmppab_b1 | ATACTGTTCAACACACAACAGTGGTTAGAAGAGTCTTCCTTTACG |
| gmppab_b1 | GAGGAGGGCAGCAAACGGAATGTAACACGGCTCCATGTAAAATGA |
| gmppab_b1 | GAAGGATCAATGTTTGCAGTAGGATTAGAAGAGTCTTCCTTTACG |
| gmppab_b1 | GAGGAGGGCAGCAAACGGAAGTCTGATGATACTGTTTGAGGTACA |
| gmppab_b1 | TATATTGCTGACCCTGCAGACTTTATAGAAGAGTCTTCCTTTACG |
| gmppab_b1 | GAGGAGGGCAGCAAACGGAATGACTCCAGAAGTGCTGTGTTTTAT |
| gmppab_b1 | GAGGAGGGCAGCAAACGGAATTCTCTTGTTGGTTCCTCTGGAAAA |
| gmppab_b1 | TTCAGAGATTGTGTTCGATTTGCTGTAGAAGAGTCTTCCTTTACG |
| gmppab_b1 | GAGGAGGGCAGCAAACGGAAATTTGGTCACGAAAGTGATAGATCC |
| gmppab_b1 | CCCAGAGCAGCAAACTCCTGCAGGTTAGAAGAGTCTTCCTTTACG |
| gmppab_b1 | GAGGAGGGCAGCAAACGGAACTGATTGGGATTTTAAACTCCTGCT |
| gmppab_b1 | GAGGAGGGCAGCAAACGGAATTTCTGCCCAAAAAAACACTATGTC |
| gmppab_b1 | ATCTGCTTCCATATGTAGACACATTTAGAAGAGTCTTCCTTTACG |
| gmppab_b1 | GAGGAGGGCAGCAAACGGAACCCATTGTGCAATATTTATTCCCAA |
| gmppab_b1 | AGTGTCTTTTAGAGAATGATCTGGTTAGAAGAGTCTTCCTTTACG |
| gmppab_b1 | GAGGAGGGCAGCAAACGGAATTGAAGCTCCTGTTGAGATCTTTAT |
| gmppab_b1 | GGGGTCAGACCTCCGTCCCTGAACATAGAAGAGTCTTCCTTTACG |
| gmppab_b1 | GAGGAGGGCAGCAAACGGAATCATTTGTCTCATGGTTTTCCACAA |
| gmppab_b1 | AAAAAAGCAGCCGGGCCCCCGGACATAGAAGAGTCTTCCTTTACG |
| gmppab_b1 | GAGGAGGGCAGCAAACGGAAAATGACCTAATCTCTCATCATTTTG |
| gmppab_b1 | AAATTTATCAGGAAAATCCCCAGTTTAGAAGAGTCTTCCTTTACG |
| gmppab_b1 | GAGGAGGGCAGCAAACGGAACTCCAAGTACCTAAACATTACATAA |
| gmppab_b1 | GAGGAGGGCAGCAAACGGAATGATAAAAGCCTATTAGCATGATCT |
| gmppab_b1 | ATTTAACCTTAACAGACCAGTTTCATAGAAGAGTCTTCCTTTACG |
| gne_b1 | GCCTTGGTTCTTTGAGTCTGTGAGTTAGAAGAGTCTTCCTTTACG |
| gne_b1 | GAGGAGGGCAGCAAACGGAATATTTCAGTAAGTTCTGCGGGTGGT |
| gne_b1 | CCAGAACCCCAGACAGAATGACCAGTAGAAGAGTCTTCCTTTACG |
| gne_b1 | GAGGAGGGCAGCAAACGGAAACGGGTTGATCATGTGCAGGATGTT |
| gne_b1 | CAGTCTCTGCTTTGGAGTTGCCGAGTAGAAGAGTCTTCCTTTACG |
| gne_b1 | GAGGAGGGCAGCAAACGGAAGAGCAGCGTTGATCAGATGAATGGC |
| gne_b1 | GCAGCAAGTCCTCGTCATGGAGTCTTAGAAGAGTCTTCCTTTACG |
| gne_b1 | GAGGAGGGCAGCAAACGGAATGGCTTCTCTCTGCAGAGCGAGACC |
| gne_b1 | ATTCTGGTCCTTCCAGCGACACCACTAGAAGAGTCTTCCTTTACG |
| gne_b1 | GAGGAGGGCAGCAAACGGAATGTGTCCCAGTTCAGCTGCGCAAAA |
| gne_b1 | CTCCAATACCGGTTCCAGTGATAATTAGAAGAGTCTTCCTTTACG |
| gne_b1 | GAGGAGGGCAGCAAACGGAATAACAAAGTTCTCCACGCCTTTGCC |
| gne_b1 | GTTTACACATGGTCAGAATCAGCTCTAGAAGAGTCTTCCTTTACG |
| gne_b1 | GAGGAGGGCAGCAAACGGAATCCTCTCCTCAAACGTCTTGGGATT |
| gne_b1 | CGATGCCGACACGCAGGTTTGTTCCTAGAAGAGTCTTCCTTTACG |
| gne_b1 | GAGGAGGGCAGCAAACGGAACCAGGTCGACAGAAAGTGCGCTCTG |
| gne_b1 | TTACTGGAGGAAAACAGAACTTCTTTAGAAGAGTCTTCCTTTACG |
| gne_b1 | GAGGAGGGCAGCAAACGGAAGCAGAGGCTCTGAGAGGTCGATGGT |
| gne_b1 | CATAGATCTTGGAGCAGGGGTAGCGTAGAAGAGTCTTCCTTTACG |
| gne_b1 | GAGGAGGGCAGCAAACGGAATTCCAAACTGCAGCTCGAGGGCATG |
| gne_b1 | CATTTTCCCCTGTTTCTCTGCCTGTTAGAAGAGTCTTCCTTTACG |
| gne_b1 | GAGGAGGGCAGCAAACGGAAGCCGAGTGCCCAGATTTATGACAGG |
| gne_b1 | TTCCGATCATGCAGCCTGCGTGAGCTAGAAGAGTCTTCCTTTACG |
| gne_b1 | GAGGAGGGCAGCAAACGGAACCAGCTGGATGAACTGATCGAACGG |
| gne_b1 | CCTTCCGCCGCATCACACGAACCATTAGAAGAGTCTTCCTTTACG |
| gne_b1 | GAGGAGGGCAGCAAACGGAACCTTACTGCCTGCGTCAATATTCGG |
| gne_b1 | CCAGCATTAACTCGTAGATCTTGATTAGAAGAGTCTTCCTTTACG |
| gne_b1 | GAGGAGGGCAGCAAACGGAAAGTTCTTGATATCTGTGGTAACAGG |
| gne_b1 | CCCCGATCCACGACTTGATGATGTCTAGAAGAGTCTTCCTTTACG |
| gne_b1 | GAGGAGGGCAGCAAACGGAACATAATCGTCTCGTTTGTAGGCGGA |
| gne_b1 | GAGAGTGATCTTCGCACATGGAGATTAGAAGAGTCTTCCTTTACG |
| gne_b1 | GAGGAGGGCAGCAAACGGAAGGTGTCTCTCGGCAGAGAGCGTGCA |
| gne_b1 | TGGAGTCGTCGATGGTTCCGCTCACTAGAAGAGTCTTCCTTTACG |
| gne_b1 | GAGGAGGGCAGCAAACGGAACTCCACCCTCCAGATGCAATATCCG |
| gne_b1 | CAAAACGGTCACCATGAACCAGCAGTAGAAGAGTCTTCCTTTACG |
| gne_b1 | GAGGAGGGCAGCAAACGGAATGTCGGGCGCGAGACGTTGGAGGAC |
| gne_b1 | TGGCAGCCTCATCCTCACCGCGAACTAGAAGAGTCTTCCTTTACG |
| gne_b1 | GAGGAGGGCAGCAAACGGAATGGTGTGTAATTTGGAGCCGATGTC |
| gne_b1 | CGTCAATGAGATGTGAACCCAGCACTAGAAGAGTCTTCCTTTACG |
| gne_b1 | GAGGAGGGCAGCAAACGGAACCACCTCCAGGTCAAAGATGTCCGG |
| gne_b1 | AGTCCGCTCGGTTGCAAGTGGCCACTAGAAGAGTCTTCCTTTACG |
| gne_b1 | GAGGAGGGCAGCAAACGGAAACACGCGGAGTCTTTCTTTCTTTGG |
| gne_b1 | TCAGATAGTAAAGCTCGTCCGACGCTAGAAGAGTCTTCCTTTACG |
| gne_b1 | GAGGAGGGCAGCAAACGGAATTATCTGTAGTTCATTCCCTTGACG |
| gne_b1 | AATCATCAACAGAGGAAGATTGGTCTAGAAGAGTCTTCCTTTACG |
| gne_b1 | GAGGAGGGCAGCAAACGGAAGTCGCGCAAAACTAGTGAGACTCTT |
| gne_b1 | AGCATCTCATCTTTGGCCCACGTATTAGAAGAGTCTTCCTTTACG |
| gne_b1 | GAGGAGGGCAGCAAACGGAATCAATAGGTTTACAACGGAACTCTA |
| gne_b1 | TAGTTGAACATAATTCTGGAAGACATAGAAGAGTCTTCCTTTACG |
| gne_b1 | GAGGAGGGCAGCAAACGGAAAAAGGCTTGGCAACAAATAAGCCAT |
| gpaa1_B2 | AACTCTAGGAAAGAAGGCAATTTTGAAATCATCCAGTAAACCGCC |
| gpaa1_B2 | CCTCGTAAATCCTCATCAAAGTGCAGAAAGACAACAAACCTTTAA |
| gpaa1_B2 | TTTGCTCTTTCACTTGTTTACTTCCAAATCATCCAGTAAACCGCC |
| gpaa1_B2 | CCTCGTAAATCCTCATCAAAAAGAGAATATTCCAGAACATCAGCC |
| gpaa1_B2 | AAAAGCGGATAGACGAGCGACCCGTAAATCATCCAGTAAACCGCC |
| gpaa1_B2 | CCTCGTAAATCCTCATCAAAAGCGCGTGATCGAGGATTCCCTGAG |
| gpaa1_B2 | AGTCCCTCAGGAAGGCTGACCGGCGAAATCATCCAGTAAACCGCC |
| gpaa1_B2 | CCTCGTAAATCCTCATCAAATCCATCAGCTCCTGATAGATGAGGA |
| gpaa1_B2 | GGACTGAGGAGCACCATGGCCAGAGAAATCATCCAGTAAACCGCC |
| gpaa1_B2 | CCTCGTAAATCCTCATCAAACTCACAGATTTGAGCATGTGTGGTG |
| gpaa1_B2 | AGTGTGACGGCCAGGATGAAGCCCAAAATCATCCAGTAAACCGCC |
| gpaa1_B2 | CCTCGTAAATCCTCATCAAAGAGAAGTTAATGAGCGCAGTGCAGC |
| gpaa1_B2 | AGTGCCAGGCCTGCAGTGTAGATGGAAATCATCCAGTAAACCGCC |
| gpaa1_B2 | CCTCGTAAATCCTCATCAAAAGAGCGGTCAGCACCACAGCTTCGG |
| gpaa1_B2 | ACAGCCATGTCCTGCAGGTAAACAGAAATCATCCAGTAAACCGCC |
| gpaa1_B2 | CCTCGTAAATCCTCATCAAAAGTATATACAGCGCCACACCAGTCA |
| gpaa1_B2 | AACACGGACAACACCCCTGGACTGGAAATCATCCAGTAAACCGCC |
| gpaa1_B2 | CCTCGTAAATCCTCATCAAAGGCTGCTCGGCTTCACTGACTCCAT |
| gpaa1_B2 | CCCAGATGAACCCAGAGATCCAACGAAATCATCCAGTAAACCGCC |
| gpaa1_B2 | CCTCGTAAATCCTCATCAAACGGAGCAGTAAGATGACCGCCAGCA |
| gpaa1_B2 | CCGATGGAGACAAACCGAGACAGAGAAATCATCCAGTAAACCGCC |
| gpaa1_B2 | CCTCGTAAATCCTCATCAAAGGCAACAAGTAAAAGAAATATGACT |
| gpaa1_B2 | TTCAGCTTACGGACCATTCCCTCCAAAATCATCCAGTAAACCGCC |
| gpaa1_B2 | CCTCGTAAATCCTCATCAAAAGCCTGCCAATGGTGGTGGCGTCCA |
| gpaa1_B2 | ATGCCGCGGATTGAGGCGGCCTCAAAAATCATCCAGTAAACCGCC |
| gpaa1_B2 | CCTCGTAAATCCTCATCAAATGATATCGGAGGAAGAGGCCGTGAT |
| gpaa1_B2 | TGTTTTAGCACCATCAGCATCATGGAAATCATCCAGTAAACCGCC |
| gpaa1_B2 | CCTCGTAAATCCTCATCAAATGAGCCGCATGGGTGTAGCCCTCAG |
| gpaa1_B2 | AGCTTTCCCTGAATGGTGCACAGAAAAATCATCCAGTAAACCGCC |
| gpaa1_B2 | CCTCGTAAATCCTCATCAAATTGAGCTTCTGACAGAAGGCATAGA |
| gpaa1_B2 | GGCAGCTGACCATTGAGTCCCTCTAAAATCATCCAGTAAACCGCC |
| gpaa1_B2 | CCTCGTAAATCCTCATCAAAACCAAATCCAGGCTGGTGATGACAT |
| gpaa1_B2 | GCAGCCTGAATCGAACCAGCACGACAAATCATCCAGTAAACCGCC |
| gpaa1_B2 | CCTCGTAAATCCTCATCAAATGCAGAGGAGAGTACTCCATTCCTG |
| gpaa1_B2 | TCCAGCCAGGCCTGCATGCCGATCAAAATCATCCAGTAAACCGCC |
| gpaa1_B2 | CCTCGTAAATCCTCATCAAATCGTGCTCATTCACCAGAAAGATGA |
| gpaa1_B2 | TTCCTGAAGTATTGTGCCAGGCCCAAAATCATCCAGTAAACCGCC |
| gpaa1_B2 | CCTCGTAAATCCTCATCAAAAGGATTCCAACGGCCTGATTGTTGG |
| gpaa1_B2 | CTGATCACGAGCGCTTCTGTCCGAGAAATCATCCAGTAAACCGCC |
| gpaa1_B2 | CCTCGTAAATCCTCATCAAAGCTCGAGGCGCTCGCAAAATCCCAT |
| gpaa1_B2 | TATCTCTCCTTGTTCTCATCAGGGAAAATCATCCAGTAAACCGCC |
| gpaa1_B2 | CCTCGTAAATCCTCATCAAAGGCAGTTTTTGAAAGAAGCTCTGCG |
| gpaa1_B2 | TGCATCGCCTTCACCAGCCACTCCAAAATCATCCAGTAAACCGCC |
| gpaa1_B2 | CCTCGTAAATCCTCATCAAAGGCATCCCATCTGCTTTTCGTTTGT |
| gpaa1_B2 | GCCAAAGCTCGTTCTCCTGCAGAAAAAATCATCCAGTAAACCGCC |
| gpaa1_B2 | CCTCGTAAATCCTCATCAAACGCTCCTCCACCATTGTGGAGCCCA |
| gpaa1_B2 | CGCAGTGTGAAGGGCTCGAACGCCAAAATCATCCAGTAAACCGCC |
| gpaa1_B2 | CCTCGTAAATCCTCATCAAACCCATGAACCACACTATCGCTGCCA |
| gpaa1_B2 | GTGTTGAGTCGGGTCAGCAGGTTGGAAATCATCCAGTAAACCGCC |
| gpaa1_B2 | CCTCGTAAATCCTCATCAAAAAGGCTTTTCTGCGATTCGGGTCGG |
| gpaa1_B2 | TACTGCTCTGAGAGAGCTGAAGAAAAAATCATCCAGTAAACCGCC |
| gpaa1_B2 | CCTCGTAAATCCTCATCAAATAGAGATTTCTTGCTGTGTGGATGC |
| gpaa1_B2 | GAAAACTTCAAACATTAAAGCAGAGAAATCATCCAGTAAACCGCC |
| gpaa1_B2 | CCTCGTAAATCCTCATCAAATGATCTGCAGCTGTAGTGCATTCAT |
| gpaa1_B2 | CTCTTTATTTGGGCCTGACGTCCTCAAATCATCCAGTAAACCGCC |
| gpaa1_B2 | CCTCGTAAATCCTCATCAAAACATCTTCATCCAGCAGCTCCTAAT |
| man1b1a_b1 | CTGCGCTGCAGTCGAGAAAGCTGCTTAGAAGAGTCTTCCTTTACG |
| man1b1a_b1 | CTCTGCAGTATCTCCCAGCCCCAGTTAGAAGAGTCTTCCTTTACG |
| man1b1a_b1 | GAGGAGGGCAGCAAACGGAACTATATTTGCTGTCCTTGGTGAATC |
| man1b1a_b1 | CTCTCAACAGTCTCGGGTCTCAGCATAGAAGAGTCTTCCTTTACG |
| man1b1a_b1 | GAGGAGGGCAGCAAACGGAATTGTGTCTGTCTGCAGGCTTCACGT |
| man1b1a_b1 | TGGTTCTGCAGGTTAAAGTGAGCAATAGAAGAGTCTTCCTTTACG |
| man1b1a_b1 | GAGGAGGGCAGCAAACGGAATCTGGACTCAGACCTGTTTCCATCT |
| man1b1a_b1 | CACGTCTCGATCAGCTGAATGGCCATAGAAGAGTCTTCCTTTACG |
| man1b1a_b1 | GAGGAGGGCAGCAAACGGAATCCATGTGATCTGCAGGCAGACCGT |
| man1b1a_b1 | AGTGTTCCAGGAAGAAAACAAACCATAGAAGAGTCTTCCTTTACG |
| man1b1a_b1 | GAGGAGGGCAGCAAACGGAATGATCCATTTTGGGATTAAGACGGC |
| man1b1a_b1 | ACAAAGGTCAGTTTACTAGTAGAAGTAGAAGAGTCTTCCTTTACG |
| man1b1a_b1 | GAGGAGGGCAGCAAACGGAATGTTTCAAAAGATTCTTCTTGACAC |
| man1b1a_b1 | TAGTCCTCCAGCAGCTCAGTCTCCTTAGAAGAGTCTTCCTTTACG |
| man1b1a_b1 | GAGGAGGGCAGCAAACGGAATTGCCGCCCTGAAGCCACTGTTTGA |
| man1b1a_b1 | CTGTCTGCTCGGGCCCCCAGGGTAATAGAAGAGTCTTCCTTTACG |
| man1b1a_b1 | GAGGAGGGCAGCAAACGGAAGCCCCTCGCCGCGTGAATTTGCCAC |
| man1b1a_b1 | ATTGGCACCAGTCCATCACGCTTCCTAGAAGAGTCTTCCTTTACG |
| man1b1a_b1 | GAGGAGGGCAGCAAACGGAATCCAATCTGTGGACCTGTTTGGTCA |
| man1b1a_b1 | TGATACTGCGGGTCCTGTGTGAGCCTAGAAGAGTCTTCCTTTACG |
| man1b1a_b1 | GAGGAGGGCAGCAAACGGAACTGAGCTCTCTGAACTCCAGCTGAA |
| man1b1a_b1 | ACTGTGCTGTCTGTGGTCCAGCGTGTAGAAGAGTCTTCCTTTACG |
| man1b1a_b1 | GAGGAGGGCAGCAAACGGAAGGATGAGCCGTTCCCTTACCAATGT |
| man1b1a_b1 | TTTCACAGTGTGGGACTTCAGCAGTTAGAAGAGTCTTCCTTTACG |
| man1b1a_b1 | GAGGAGGGCAGCAAACGGAAGCCTTGTGATTTACGAAGCTCTGAC |
| man1b1a_b1 | GTGTCGCCACTCAGGTGATATGTGCTAGAAGAGTCTTCCTTTACG |
| man1b1a_b1 | GAGGAGGGCAGCAAACGGAAAAGAGACCTCCTAAAATGCGAATGG |
| man1b1a_b1 | ACGTCCACATTCTTGGAGAAGGATATAGAAGAGTCTTCCTTTACG |
| man1b1a_b1 | GAGGAGGGCAGCAAACGGAATCTGTCTCCACCCACTTCCTGGCCT |
| man1b1a_b1 | AGACCAAGGATCCACATGGTGTCCATAGAAGAGTCTTCCTTTACG |
| man1b1a_b1 | GAGGAGGGCAGCAAACGGAAGCATCTATGAGCGTGAGGCCCAGAC |
| man1b1a_b1 | GTCTTAGATAGTGGCTTCAGCTCATTAGAAGAGTCTTCCTTTACG |
| man1b1a_b1 | GAGGAGGGCAGCAAACGGAATGACCCCATGCAAAAGCCTTATAAC |
| man1b1a_b1 | AAGGCCTCCTGCACCGCTTCCACACTAGAAGAGTCTTCCTTTACG |
| man1b1a_b1 | GAGGAGGGCAGCAAACGGAATCCACTGGCTTCAATGTCCCTGTTT |
| man1b1a_b1 | TCAGCATCAACCACAGTTGCATCCTTAGAAGAGTCTTCCTTTACG |
| man1b1a_b1 | GAGGAGGGCAGCAAACGGAACCCTGAATGGTGGACTGAAGCTCAG |
| man1b1a_b1 | ATCATCGCCCCGCGCCAGCTAATGATAGAAGAGTCTTCCTTTACG |
| man1b1a_b1 | GAGGAGGGCAGCAAACGGAAGTTTTTGCTTCTTTTGCTCCGTCTT |
| man1b1a_b1 | CCTTCTTCTCCATCTTTCAGCTGCCTAGAAGAGTCTTCCTTTACG |
| man1b1a_b1 | GAGGAGGGCAGCAAACGGAATCGGACACGTTCATTCGCTTCTGTA |
| man1b1a_b1 | TTGCTAGGAGGTTTTCTCATAGCTGTAGAAGAGTCTTCCTTTACG |
| man1b1a_b1 | GAGGAGGGCAGCAAACGGAAGGTTTAGGAAGTAATGGAATGCTGG |
| man1b1a_b1 | AAATATGGACGGATATGGCAGTTATTAGAAGAGTCTTCCTTTACG |
| man1b1a_b1 | ATAACATCTGGTTGGACGTCTTGAATAGAAGAGTCTTCCTTTACG |
| man1b1a_b1 | GAGGAGGGCAGCAAACGGAACCAACTCCTGCAGACAGCTTTGGGA |
| man1b1a_b1 | GAGGAGGGCAGCAAACGGAACGACATGCAAGAGTGTCACAAGTTG |
| man1b1a_b1 | GGCCTGAGACCCCGATTATCCACTTTAGAAGAGTCTTCCTTTACG |
| man1b1a_b1 | GAGGAGGGCAGCAAACGGAATCCCAATCCTCCTTCAATGACCGAT |
| man1b1a_b1 | TGCTCCGTGAGACTGGAATAGGACATAGAAGAGTCTTCCTTTACG |
| man1b1a_b1 | GAGGAGGGCAGCAAACGGAAATCCCACATATAAAGAGGAAGGCCA |
| man1b1a_b1 | GAGGAGGGCAGCAAACGGAACATTTCCTCCAGCAGGACTGTCTCC |
| man1b1a_b1 | TAGCCACGGTTGTGTTGTTCTCGAATAGAAGAGTCTTCCTTTACG |
| man1b1a_b1 | GAGGAGGGCAGCAAACGGAAGTCAAAGACACAAAATCCTTCTTCT |
| man1b1a_b1 | ACCAGACCACCACTCAAAGACTCTTTAGAAGAGTCTTCCTTTACG |
| man1b1a_b1 | GAGGAGGGCAGCAAACGGAAATGGGACAAGTTGCTTCTCCTGGTC |
| man1b1a_b1 | GGAGGTCACACGGCCAACACATCCATAGAAGAGTCTTCCTTTACG |
| man1b1a_b1 | GAGGAGGGCAGCAAACGGAACAGAGTGAGTGTAGTATTAGTATAC |
| man1b1a_b1 | AGGAGTGGCTATCAGATCTCTATCTTAGAAGAGTCTTCCTTTACG |
| man1b1a_b1 | GAGGAGGGCAGCAAACGGAACCTCCATCACGCTGCTCTCCACAAA |
| man1b1a_b1 | AACCGCTCATTCATGCTGATCTACATAGAAGAGTCTTCCTTTACG |
| man1b1a_b1 | GAGGAGGGCAGCAAACGGAAATATCCGTAATGTGTTAATCACTAC |
| man1b1a_b1 | TTGTCATTAGTCAAACAGATGTGTGTAGAAGAGTCTTCCTTTACG |
| man1b1a_b1 | GAGGAGGGCAGCAAACGGAATTCAGTGCAATGTTTATATGTGACA |
| man1b1a_b1 | CCAAATAAAGCTCAAGTGCATAAGCTAGAAGAGTCTTCCTTTACG |
| man1b1a_b1 | GAGGAGGGCAGCAAACGGAAAGGTCCTGATGGGTGGCTTATGGTG |
| man1b1a_b1 | GGTCAAAGTCAATATCACTGAAATTTAGAAGAGTCTTCCTTTACG |
| man1b1a_b1 | GAGGAGGGCAGCAAACGGAAGAGTGAGCGCCGGCATTGGTTCCTC |
| man1b1a_b1 | ATGGGCAGCGGATGGGCTTCAGTGTTAGAAGAGTCTTCCTTTACG |
| man1b1a_b1 | GAGGAGGGCAGCAAACGGAAAACACGTATTTATCCAGACTGATGA |
| man1b1a_b1 | AACAGGAGGAAGAGATACTTTAGCGTAGAAGAGTCTTCCTTTACG |
| man1b1a_b1 | GAGGAGGGCAGCAAACGGAATCGCTCAGGAAGAAGCTCTCCATCT |
| man1b1a_b1 | TTTAACGGGTCCCGGACGTTACCTATAGAAGAGTCTTCCTTTACG |
| man1b1a_b1 | GAGGAGGGCAGCAAACGGAAGAAGTGTAGCCACCACTGGGAACTC |
| man1b1b_b3 | CCAACGTGCCAGGCAGGAAACATACTTCCACTCAACTTTAACCCG |
| man1b1b_b3 | GTCCCTGCCTCTATATCTTTAATGGTCCATTTTGGGACTGAAGTG |
| man1b1b_b3 | GATTGTATATTAAATGCGTTTTGCGTTCCACTCAACTTTAACCCG |
| man1b1b_b3 | AACTCTCCTACAAATGTAAGGCCAATTCCACTCAACTTTAACCCG |
| man1b1b_b3 | GTCCCTGCCTCTATATCTTTGGTGTCGATTTCTTCAGCAGGTTCT |
| man1b1b_b3 | GTCCCTGCCTCTATATCTTTCCTGAGAGAGGTGTGTTTGGTTGTG |
| man1b1b_b3 | TTCCACAGCCTGCAGATAGTCCTCCTTCCACTCAACTTTAACCCG |
| man1b1b_b3 | GTCCCTGCCTCTATATCTTTCAATTCCTGCTCGATTCTGCCTCCC |
| man1b1b_b3 | GCTTAAGCAGATACTCATAGTAGCTTTCCACTCAACTTTAACCCG |
| man1b1b_b3 | GTCCCTGCCTCTATATCTTTCAGCCCTGGCGCCTAGCGTGTAAAT |
| man1b1b_b3 | GTGAACAGTCCATTGTTGGTATTGATTCCACTCAACTTTAACCCG |
| man1b1b_b3 | GTCCCTGCCTCTATATCTTTAACATCGGCACAAGCCCATCTTGCT |
| man1b1b_b3 | TTTGTGCACCTGCTTCATCACCTCCTTCCACTCAACTTTAACCCG |
| man1b1b_b3 | GTCCCTGCCTCTATATCTTTCACAGCCAGCTTGTATTTGGGGTCT |
| man1b1b_b3 | GGCTGAGCTCTCTGAACTCCAGCTGTTCCACTCAACTTTAACCCG |
| man1b1b_b3 | GTCCCTGCCTCTATATCTTTTGCTAGTGACCTCGGCTACGGTGCT |
| man1b1b_b3 | CATCGCGGAGGATGAGCCGTGCCCTTTCCACTCAACTTTAACCCG |
| man1b1b_b3 | GTCCCTGCCTCTATATCTTTCCAATATTCACATCAGAATAGGGGA |
| man1b1b_b3 | AGTGCTGAAGGCGGGCATCAGTCTGTTCCACTCAACTTTAACCCG |
| man1b1b_b3 | GTCCCTGCCTCTATATCTTTGCCAATCTCTTTAGCTTTATCCAGG |
| man1b1b_b3 | CTCCTGTGAGATGGTAGGTACTCAGTTCCACTCAACTTTAACCCG |
| man1b1b_b3 | GTCCCTGCCTCTATATCTTTGGCCACCCAGGATACGAATGGTGCT |
| man1b1b_b3 | TTAACATCCACATTCTTATCGAAAGTTCCACTCAACTTTAACCCG |
| man1b1b_b3 | GTCCCTGCCTCTATATCTTTAGCTCTTTGGCCACCCACTCCCTGG |
| man1b1b_b3 | TCTGCTTGTGGCCATACAGGCAGAGTTCCACTCAACTTTAACCCG |
| man1b1b_b3 | CTCTTCTTGTAGACCCAAGATCCACTTCCACTCAACTTTAACCCG |
| man1b1b_b3 | GTCCCTGCCTCTATATCTTTGGTATCTAGAGCATCAATTAGGGTC |
| man1b1b_b3 | GTCCCTGCCTCTATATCTTTTGAGCCTCAGTGTTGAACACGTACT |
| man1b1b_b3 | CAAACCACTCTCCGTATGATTTAGATTCCACTCAACTTTAACCCG |
| man1b1b_b3 | GTCCCTGCCTCTATATCTTTTTGGTTTGAGTTCATCGTGACCCCA |
| man1b1b_b3 | TTGTAACCTTTCCAGGCATGTCTAATTCCACTCAACTTTAACCCG |
| man1b1b_b3 | GTCCCTGCCTCTATATCTTTGCTTCTCTCACAGCCTCCAGTCTAC |
| man1b1b_b3 | TGCTAATGAAGCTGGGTCTCCTTGGTTCCACTCAACTTTAACCCG |
| man1b1b_b3 | GTCCCTGCCTCTATATCTTTTTCTGTGTCTTTGGTGTCAGTATCA |
| man1b1b_b3 | CTTCTTCGATCACAGCTCCACGCCATTCCACTCAACTTTAACCCG |
| man1b1b_b3 | GTCCCTGCCTCTATATCTTTTAATTATAGGCTTTTGTCCCTCAGA |
| man1b1b_b3 | GCATTTTTGAGGTCCAGTCCATGATTTCCACTCAACTTTAACCCG |
| man1b1b_b3 | GTCCCTGCCTCTATATCTTTTGGTCTCCCAACACACCCGAAACAT |
| man1b1b_b3 | TTGCAAAAGTTTCTCCAAACCAAAATTCCACTCAACTTTAACCCG |
| man1b1b_b3 | GTCCCTGCCTCTATATCTTTTCCACGTCTATCGAAATCTTTTCTC |
| man1b1b_b3 | GCTCGGGAAGAATGGACCTGTACTCTTCCACTCAACTTTAACCCG |
| man1b1b_b3 | GTCCCTGCCTCTATATCTTTTTTTCAGGTCAACAACAAATGAGTT |
| man1b1b_b3 | TCCCCATTCATTCCTCTCCAGTGTCTTCCACTCAACTTTAACCCG |
| man1b1b_b3 | GTCCCTGCCTCTATATCTTTATGAGGGTGGGAGATGTGGCGATTC |
| man1b1b_b3 | CAACAGCATCAACGTAAGCAGAATCTTCCACTCAACTTTAACCCG |
| man1b1b_b3 | GTCCCTGCCTCTATATCTTTGCTGCGCTGCAATCTGGACAGCTGT |
| man1b1b_b3 | GATAAGGTTGGGGTCTTCTGAAAACTTCCACTCAACTTTAACCCG |
| man1b1b_b3 | TCCAGCAAGACTGCCGTCTCCACTGTTCCACTCAACTTTAACCCG |
| man1b1b_b3 | GTCCCTGCCTCTATATCTTTTGCTGTTACTGTAGCTGCCGCTTTC |
| man1b1b_b3 | GTCCCTGCCTCTATATCTTTCAGGTAGAAGTATTTGAGCGTTTCC |
| man1b1b_b3 | AGAGATACAAAATCCTTTCTTGTTGTTCCACTCAACTTTAACCCG |
| man1b1b_b3 | GTCCCTGCCTCTATATCTTTGGCAGCATGATTCCCTTTCTCTCTC |
| man1b1b_b3 | ATAAACTCAGTGGCCTGGTCTCTTTTTCCACTCAACTTTAACCCG |
| man1b1b_b3 | GTCCCTGCCTCTATATCTTTCTTGTTTTCCTGGAGTTTATCGACG |
| man1b1b_b3 | GGGAAGCGGAGCTCTGATCGGCTCATTCCACTCAACTTTAACCCG |
| man1b1b_b3 | GTCCCTGCCTCTATATCTTTGTTTCACGACGGTCCGAGGCATCAC |
| man1b1b_b3 | AGAGATCACCGGCGATCACTCTCACTTCCACTCAACTTTAACCCG |
| man1b1b_b3 | GTCCCTGCCTCTATATCTTTCCGAAAACCTCAGTCGACTCTTCAT |
| man1b1b_b3 | TAACGCGCGATTTTCCCGACGACAATTCCACTCAACTTTAACCCG |
| man1b1b_b3 | GTCCCTGCCTCTATATCTTTCTGAGAAGCCAGGAGGATCAAACCG |
| man1b1b_b3 | AGCCATTTTCTCAGCTGATGTGTCATTCCACTCAACTTTAACCCG |
| man1b1b_b3 | GTCCCTGCCTCTATATCTTTTGATTTCCTGGAGACGAATACACTA |
| man1b1b_b3 | AGCTCTCCATCTTGTCTCGAGGACTTTCCACTCAACTTTAACCCG |
| man1b1b_b3 | GTCCCTGCCTCTATATCTTTTGTAGGAAGGGTCACGGACGTTGTT |
| man1b1b_b3 | TAACCTCCGGTTGGAACCCTTGTGTTTCCACTCAACTTTAACCCG |
| man1b1b_b3 | GTCCCTGCCTCTATATCTTTTTGTTAAAGTTCTGCAGGATCTCCC |
| man1b1b_b3 | CTGGTACCTCTTATCCCTGGTCAATTTCCACTCAACTTTAACCCG |
| man1b1b_b3 | GTCCCTGCCTCTATATCTTTGTACAGGTAGAAGAGGCTCTCCACG |
| man1b1b_b3 | GCAGAAGATTGTGGCGGTCCGCAATTTCCACTCAACTTTAACCCG |
| man1b1b_b3 | GTCCCTGCCTCTATATCTTTTCACATCTACATCCTGTGTGCTACC |
| man1b1b_b3 | TTGAAATGGGCTATTTCTGGACTCATTCCACTCAACTTTAACCCG |
| man1b1b_b3 | GTCCCTGCCTCTATATCTTTCCGGTCTCCATCTGTGCATACATCT |
| man1b1b_b3 | CTCTATCAGCTGTTTCGCAAGCTCCTTCCACTCAACTTTAACCCG |
| man1b1b_b3 | GTCCCTGCCTCTATATCTTTATGATCAGCGGGAAGGCCGTAATGC |
| nansa_b2 | ACATGCCGCTCCACAACTTTGGCTCAAATCATCCAGTAAACCGCC |
| nansa_b2 | CCTCGTAAATCCTCATCAAAAGAGCAACTGCCCCGACTGTGATGT |
| nansa_b2 | CATGACCCGAGTAACCGATGGGAATAAATCATCCAGTAAACCGCC |
| nansa_b2 | CCTCGTAAATCCTCATCAAACTGGGAATTCTTTCTGATATTCCGT |
| nansa_b2 | GTTGACGTCCTCAGGTTCAAGTGGAAAATCATCCAGTAAACCGCC |
| nansa_b2 | CCTCGTAAATCCTCATCAAATGCACTTGTGCACTGCAGGATGCAA |
| nansa_b2 | TGCTCCTTGACAATCTTATAAACGTAAATCATCCAGTAAACCGCC |
| nansa_b2 | CCTCGTAAATCCTCATCAAATGCATGGTTTCCATGGACTGCATCC |
| nansa_b2 | CCATGGGGCGTCCTTTTTGGGCAGTAAATCATCCAGTAAACCGCC |
| nansa_b2 | CCTCGTAAATCCTCATCAAATCTTGAGGTAAGGAAAGTTGTTGGT |
| nansa_b2 | AACTTTGAAAAAAGGCACATCGAGTAAATCATCCAGTAAACCGCC |
| nansa_b2 | CCTCGTAAATCCTCATCAAAATGGAGGAATTCTACAGCCATCTCG |
| nansa_b2 | GCAGTGAAGAAGATGCCCACTTGTTAAATCATCCAGTAAACCGCC |
| nansa_b2 | CCTCGTAAATCCTCATCAAAGCGTACTGCTGCAGCTCCCTGTACT |
| nansa_b2 | ACTCCAGGTGTCGCTTATGCTCTCCAAATCATCCAGTAAACCGCC |
| nansa_b2 | CCTCGTAAATCCTCATCAAAATGTTTTCCCCCAGGAGTGTTTGGA |
| nansa_b2 | TTCAAGAGCTTTCTTATTGAATTTAAAATCATCCAGTAAACCGCC |
| nansa_b2 | CCTCGTAAATCCTCATCAAACTCCAGCTCACTCTTCTGAAACTTT |
| nansa_b2 | CCACAGTCCTTTGCCATTTTGATCAAAATCATCCAGTAAACCGCC |
| nansa_b2 | CCTCGTAAATCCTCATCAAATTCTTGGCAATCTCAATGTCTCCCT |
| nansa_b2 | CGATTTCTGCAATAATGAAGCACGGAAATCATCCAGTAAACCGCC |
| nansa_b2 | CCTCGTAAATCCTCATCAAATGTTTCCTCCGATCATGCGGCCGGG |
| nansa_b2 | ATGCATTATATAAACAAGTCTGGCTAAATCATCCAGTAAACCGCC |
| nansa_b2 | TTTAAGTGGCATTCTCACAACTAAAAAATCATCCAGTAAACCGCC |
| nansa_b2 | CCTCGTAAATCCTCATCAAACACAAACAAGACAAGAAAAGCTACA |
| nansa_b2 | CCTCGTAAATCCTCATCAAAATACACAGTATTTGTTGTATTGTTG |
| nansa_b2 | CCTCAAAAACAAAACTTGATTTTAAAAATCATCCAGTAAACCGCC |
| nansa_b2 | CCTCGTAAATCCTCATCAAAAGGGCACTGAAACATGTAACCTTTT |
| nansa_b2 | TCAAGGATCACATGCTTTCCTCAAGAAATCATCCAGTAAACCGCC |
| nansa_b2 | CCTCGTAAATCCTCATCAAATTAGCTTTTTTGCCATAGCTGTCGA |
| nansa_b2 | TAATGCTCTCGTCCTCCTCCACATCAAATCATCCAGTAAACCGCC |
| nansa_b2 | CCTCGTAAATCCTCATCAAATGGTCACTTTCTTCCCCACCAGCTG |
| nansa_b2 | TGGAGCTACACCCTTGGGTTCAGCTAAATCATCCAGTAAACCGCC |
| nansa_b2 | CCTCGTAAATCCTCATCAAATTTCACTGCCAGCATGTCAAGGGTG |
| nansa_b2 | TTTGGAATGGAGGTCTTTGCCACCAAAATCATCCAGTAAACCGCC |
| nansa_b2 | CCTCGTAAATCCTCATCAAAGACTTCCCGAGCTTATCATGGCATG |
| nansa_b2 | GCAGCATGCGTTTCAGTCCAGTGCCAAATCATCCAGTAAACCGCC |
| nansa_b2 | CCTCGTAAATCCTCATCAAAAAGCTCTCTCCACAATGCGGATGGA |
| nansa_b2 | AGCCAGCTCTTCTGGTTCCAGAGACAAATCATCCAGTAAACCGCC |
| nansa_b2 | CCTCGTAAATCCTCATCAAATGCATGATCACTACCTTTCCAGCTC |
| nansb_b3 | CATGGTCCAAACCTGTTTGAGTTTCTTCCACTCAACTTTAACCCG |
| nansb_b3 | GTCCCTGCCTCTATATCTTTTTCTCCTGATGAACACAAAAGGAGA |
| nansb_b3 | CTACCAGTCTTCAAAATATCTTCTGTTCCACTCAACTTTAACCCG |
| nansb_b3 | GTCCCTGCCTCTATATCTTTGTGTTCAACAGACTCATGAAACACA |
| nansb_b3 | GAACTAGAGAATGCTCATTTTTGGCTTCCACTCAACTTTAACCCG |
| nansb_b3 | GTCCCTGCCTCTATATCTTTAACTGTCCCTTTATGTTTTAAATGG |
| nansb_b3 | CCTTTAATCATGGCATCAGTAATTGTTCCACTCAACTTTAACCCG |
| nansb_b3 | GTCCCTGCCTCTATATCTTTGCATCTTTCTCCAAGTTCACCGATA |
| nansb_b3 | ACAAGTTTGAAGATGTTCTCTGGTCTTCCACTCAACTTTAACCCG |
| nansb_b3 | GTCCCTGCCTCTATATCTTTACACCTTGTGGTTCGGCCACTTTCA |
| nansb_b3 | TCGAGGGTTAAAGTCTCACCCATTTTTCCACTCAACTTTAACCCG |
| nansb_b3 | GTCCCTGCCTCTATATCTTTAGTGGTTTCCGGGCCACCACTGATT |
| nansb_b3 | CTGTGGCATGAAGCCTCACAGGGCATTCCACTCAACTTTAACCCG |
| nansb_b3 | GTCCCTGCCTCTATATCTTTATCTGTTTGACTGGTGACCCCATGG |
| nansb_b3 | GCCCGGATGGCTTTCACCAGCTCTGTTCCACTCAACTTTAACCCG |
| nansb_b3 | GTCCCTGCCTCTATATCTTTAGTTCATCAGGCTCCAGCGAGGCCG |
| nansb_b3 | CGTTTCTCAGAGCCTTGTTGCCACGTTCCACTCAACTTTAACCCG |
| nansb_b3 | CCTTTCCAGCTTTTATCCAGAGTCATTCCACTCAACTTTAACCCG |
| nansb_b3 | GTCCCTGCCTCTATATCTTTTGACGCTCCAGGACTTTTGCTCCGA |
| nansb_b3 | GTCCCTGCCTCTATATCTTTTGTAAAATGAACCCACGTGTATACG |
| nansb_b3 | GCTACAGACACATGGATGCCTGTCTTTCCACTCAACTTTAACCCG |
| nansb_b3 | GTCCCTGCCTCTATATCTTTTGTCCGGAGTATCCGATGGGAATGT |
| nansb_b3 | TTCTGGAACTCGGTGATCAGACTGATTCCACTCAACTTTAACCCG |
| nansb_b3 | GTCCCTGCCTCTATATCTTTTTGACGTGCTCTAGCGGCAGCGGAT |
| nansb_b3 | CACTGCAGAAACGTGAAATTAGCGTTTCCACTCAACTTTAACCCG |
| nansb_b3 | GTCCCTGCCTCTATATCTTTTGTTTCTTGACAGTTTGATAAACGC |
| nansb_b3 | TCCATACTCTGCATTCCACTGGAAATTCCACTCAACTTTAACCCG |
| nansb_b3 | GTCCCTGCCTCTATATCTTTACCATAGGACGACCCTTTTTGGCTG |
| nansb_b3 | TAGGGGATGTTGTTGGTGTCGGCTGTTCCACTCAACTTTAACCCG |
| nansb_b3 | GTCCCTGCCTCTATATCTTTGCCACTTTGAAAAACGGCACGTTTA |
| nansb_b3 | AACTCCACCGCCATCTCGTCCATGCTTCCACTCAACTTTAACCCG |
| nansb_b3 | GTCCCTGCCTCTATATCTTTGATGCGGTAAAAAAGATGCCGATGT |
| nansb_b3 | TGCTGTAGTTCTCTGTACTGCTCGTTTCCACTCAACTTTAACCCG |
| nansb_b3 | GTCCCTGCCTCTATATCTTTCTGAACTCTAGATGGTGCTTATGAG |
| nansb_b3 | GGGCCCCAAGCGTGAGGGGATGTGTTTCCACTCAACTTTAACCCG |
| nansb_b3 | GTCCCTGCCTCTATATCTTTAGCCGATCCAGAGCGTTTCTGGTGA |
| nansb_b3 | ATCTCGCTCTTCTGAAACTTTGCACTTCCACTCAACTTTAACCCG |
| nansb_b3 | GTCCCTGCCTCTATATCTTTTCGGCTCCACAGTCCTTGGCCATTC |
| nansb_b3 | TTGGCGATTTCTACATCTCCCTGATTTCCACTCAACTTTAACCCG |
| nansb_b3 | GTCCCTGCCTCTATATCTTTTTCTGTCCGATCTCGGCGATGATAA |
| nansb_b3 | GAGCCCCCAATTTTTCTCCCAGGGCTTCCACTCAACTTTAACCCG |
| nansb_b3 | GTCCCTGCCTCTATATCTTTAGTTCAAACTCGACAGGCATTTTAT |
| nansb_b3 | TTAACTGATCAGTGATAAAGATGCTTTCCACTCAACTTTAACCCG |
| nansb_b3 | GTCCCTGCCTCTATATCTTTAGACGTGTTTCACCTGCGCTGGAAT |
| nansb_b3 | TTGAATAGCCTGCAAAACTAGAGCATTCCACTCAACTTTAACCCG |
| nansb_b3 | GTCCCTGCCTCTATATCTTTTTGAAATGCTTAAGGCCTATTAAGG |
| nansb_b3 | TGTAAGGTCAATAAAAGCAAATGCCTTCCACTCAACTTTAACCCG |
| nansb_b3 | GTCCCTGCCTCTATATCTTTTATCACTGGAAATGAGCGAAACATA |
| nansb_b3 | AACAGTTACAAAGTGTTGAGAGATTTTCCACTCAACTTTAACCCG |
| nansb_b3 | GTCCCTGCCTCTATATCTTTGTTGATATACTTGTGGTGTAGTGCT |
| ogt_b1 | GAGGAGGGCAGCAAACGGAACTGCATTTTGACCAGGTGGTCAGGT |
| ogt_b1 | CGGGCACGCACCTTCTTCAAGAATTTAGAAGAGTCTTCCTTTACG |
| ogt_b1 | GATGAGTTCAGGACAGCCAAGACAGTAGAAGAGTCTTCCTTTACG |
| ogt_b1 | TGTCCACGTCGTACATGCTCCTCTTTAGAAGAGTCTTCCTTTACG |
| ogt_b1 | GTATTGCTGGATATTGGGCTCACCCTAGAAGAGTCTTCCTTTACG |
| ogt_b1 | GAGGAGGGCAGCAAACGGAAATGGTTCTGGGAACTTCTTCTCCTG |
| ogt_b1 | TACAGTCCCAGATACTACAACCTCCTAGAAGAGTCTTCCTTTACG |
| ogt_b1 | GAGGAGGGCAGCAAACGGAACACTGTAAAAATCTGTGTGGTGGCC |
| ogt_b1 | TGGTGACTTGGATCTGGCCTTGGTTTAGAAGAGTCTTCCTTTACG |
| ogt_b1 | GAGGAGGGCAGCAAACGGAATCATGTTAATGATGGCCTCAGCTGC |
| ogt_b1 | AGGGCACCATTACTGTCAGCAACCTTAGAAGAGTCTTCCTTTACG |
| ogt_b1 | TCAGCAACCTCAATGGGAGACGTTGTAGAAGAGTCTTCCTTTACG |
| ogt_b1 | GAGGAGGGCAGCAAACGGAAGCGCAATGCAAACAGCTCATTGCGT |
| ogt_b1 | CCAGGATGTGGATTCCATCTTGATGTAGAAGAGTCTTCCTTTACG |
| ogt_b1 | TGAGCCTCTGCCATGACTTTAACTCTAGAAGAGTCTTCCTTTACG |
| ogt_b1 | AGAGTTGTGCATGCCAGGAATGGATTAGAAGAGTCTTCCTTTACG |
| ogt_b1 | AAGTTGCCATGTCTCTCTGCAATGGTAGAAGAGTCTTCCTTTACG |
| ogt_b1 | GCAATTGCTTCAGGAATGTTCCCAGTAGAAGAGTCTTCCTTTACG |
| ogt_b1 | CTGAATGGCTCGAGTGTAGCATTGCTAGAAGAGTCTTCCTTTACG |
| ogt_b1 | CATCCGCAAATGTAGGGCTGATTCTTAGAAGAGTCTTCCTTTACG |
| ogt_b1 | ACACTGGCAAGGTTTGAATGAGCAGTAGAAGAGTCTTCCTTTACG |
| ogt_b1 | GAGGAGGGCAGCAAACGGAAGCAAATTCTGGGAACACCTCCAGTG |
| ogt_b1 | AATGTTGCCCTGCTCACGCTTGATGTAGAAGAGTCTTCCTTTACG |
| ogt_b1 | GAGGAGGGCAGCAAACGGAAGTGTCAATGGCTAGGTCGATCAGAC |
| ogt_b1 | TGCCTTCAGGTAACAAGCCTTGGCTTAGAAGAGTCTTCCTTTACG |
| ogt_b1 | GAGGAGGGCAGCAAACGGAAACACATATGCTTGAACTGCTCCCTC |
| ogt_b1 | GAGGAGGGCAGCAAACGGAATCTGGTTTGAGACGAAGAGCGTGAC |
| ogt_b1 | ACCTCTCTCCTTGTAAACATTGCCCTAGAAGAGTCTTCCTTTACG |
| ogt_b1 | GAGGAGGGCAGCAAACGGAAGTTAGAGTAGGCCTCAGCTAGCATG |
| ogt_b1 | GAGGAGGGCAGCAAACGGAAACTGGAAGTGGATAGAGGAGAGCAG |
| ogt_b1 | GAGGAGGGCAGCAAACGGAATCTGCTGCCTCAAAATCTCCTGACT |
| ogt_b1 | CCCTTGGAAGGAAAGAACACGTTTGTAGAAGAGTCTTCCTTTACG |
| ogt_b1 | TCTGCAGGTAAAGTTTCTCTAGGTCTAGAAGAGTCTTCCTTTACG |
| ogt_b1 | GAGGAGGGCAGCAAACGGAATGGTGTACTGTTTGGTGTTGAAGAG |
| ogt_b1 | GAGGAGGGCAGCAAACGGAATGCCTGTGGTGTGGCCATTACAGAG |
| ogt_b1 | GAGGAGGGCAGCAAACGGAAGGAGCCACAGGAGAGAAGATGATGC |
| ogt_b1 | CCATATTGTGAGCGGGTGGTCACCATAGAAGAGTCTTCCTTTACG |
| ogt_b1 | GGCTTTCAGATCAATGCCATTCAGATAGAAGAGTCTTCCTTTACG |
| ogt_b1 | TCTTAAGGTGAGGGAACATGTTAGCTAGAAGAGTCTTCCTTTACG |
| ogt_b1 | GAGGAGGGCAGCAAACGGAATCCTGTCTGCAGCTTTTCCATTACA |
| ogt_b1 | GAGGAGGGCAGCAAACGGAAAAGTTGGTGCCGTCATCAGGGCTGA |
| ogt_b1 | GAGGAGGGCAGCAAACGGAACATAAGATGGGAAGTAGGATGGTTG |
| ogt_b1 | GCAATCGGCCTGAACTGGCTTTCAGTAGAAGAGTCTTCCTTTACG |
| ogt_b1 | GAGGAGGGCAGCAAACGGAACCTTTGGATGCTCATAAGCAGGTTT |
| ogt_b1 | GAGGAGGGCAGCAAACGGAATTGCGGAAACCGTGAGAGAGGGGAT |
| ogt_b1 | TGGCAGGCGGTTCTTCTCCAACTGATAGAAGAGTCTTCCTTTACG |
| ogt_b1 | GAGGAGGGCAGCAAACGGAAAGCAACAATGCTCACCAGTTTCTTC |
| ogt_b1 | GAGGAGGGCAGCAAACGGAATGGCCTCCTTATAGTGCATGAGGGC |
| ogt_b1 | GAGGAGGGCAGCAAACGGAAGGCCAGATTGTTAAGGGAGTCAGCG |
| ogt_b1 | TGTAACACTCCTCAGCTTCAGAAACTAGAAGAGTCTTCCTTTACG |
| ogt_b1 | TCCATGGACAACAGCGTGGTTTGGGTAGAAGAGTCTTCCTTTACG |
| ogt_b1 | GAGGAGGGCAGCAAACGGAACAGACTAAGGGCTCGCAGATACCCA |
| ogt_b1 | GAGGAGGGCAGCAAACGGAAAAGCATCCAAAAAGTTTGGGTCCAG |
| ogt_b1 | GAGGAGGGCAGCAAACGGAAAACACACAGCCCAGGTTACTCCAGG |
| ogt_b1 | GAGGAGGGCAGCAAACGGAATTCAAGGCGCCCAAGCGCTTTAAGC |
| ogt_b1 | GCCAGGTTAATGTATCCATCGATGATAGAAGAGTCTTCCTTTACG |
| ogt_b1 | GAGGAGGGCAGCAAACGGAATGGTTCTGTGCTGTCAGCCACGTTC |
| ogt_b1 | TCAGGTGCTCTCGCTGGTCTCCAGATAGAAGAGTCTTCCTTTACG |
| ogt_b1 | GAGGAGGGCAGCAAACGGAAATATCAGTGCCAAGCTTTACTGCAA |
| ogt_b1 | GAGGAGGGCAGCAAACGGAAGAGCTGAGAAGCAGCGACGCGTGAA |
| ogt_b1 | GAGGAGGGCAGCAAACGGAATGACCATCACATTCCATCTTCACAA |
| ogt_b1 | GAGGAGGGCAGCAAACGGAAAATGCGATTGTCAAAAATGTGCCCA |
| ogt_b1 | GAGGAGGGCAGCAAACGGAAGATCTCCAATGAAGAAAGTATTCGG |
| ogt_b1 | AAGCCACATGGCCTGAATGGGCGCTTAGAAGAGTCTTCCTTTACG |
| ogt_b1 | GAGGAGGGCAGCAAACGGAATACAATAAGCATCTGGGAAATCTGG |
| ogt_b1 | GAGGAGGGCAGCAAACGGAACGCCCCCTGAACATCCTGCATCTCT |
| ogt_b1 | GAGGAGGGCAGCAAACGGAATGCCTTTTTCTTTCAGAGCATTAGC |
| ogt_b1 | AGTGAGCTGACCTGTCTAGCCTGCGTAGAAGAGTCTTCCTTTACG |
| ogt_b1 | TGCCTCCACAGCTGCATACAGTGGCTAGAAGAGTCTTCCTTTACG |
| ogt_b1 | TGGGGGTACCAGCCCACAGCACATCTAGAAGAGTCTTCCTTTACG |
| ogt_b1 | TGGCCCACATCTGCAGGGTTGGAGGTAGAAGAGTCTTCCTTTACG |
| ogt_b1 | GAGGAGGGCAGCAAACGGAACAATCTTATAGAGCTGGTTGAAGTT |
| ogt_b1 | ATACAATCTGTAGACAATGAGCAAGTAGAAGAGTCTTCCTTTACG |
| ogt_b1 | GAGGAGGGCAGCAAACGGAATCTTTGTGTATGGAAGCCAAATTAC |
| ogt_b1 | GGCTGCAGCTCAATGGCACGACGGTTAGAAGAGTCTTCCTTTACG |
| ogt_b1 | CTTTCAGAACATTACCCAAGTTGATTAGAAGAGTCTTCCTTTACG |
| ogt_b1 | GCCAGCCAGATCTCACCCTGGGCATTAGAAGAGTCTTCCTTTACG |
| ogt_b1 | ATAGATCAGGATTATACTGAAGAGCTAGAAGAGTCTTCCTTTACG |
| ogt_b1 | GAGGAGGGCAGCAAACGGAAGGCTGGGAAGCGCAGCAGCCAGATC |
| ogt_b1 | GAGGAGGGCAGCAAACGGAATTATCACTGACAATGTAGTCCATAA |
| pigk_b3 | TGAGGACACTCCCCGGATCCCTTTGTTCCACTCAACTTTAACCCG |
| pigk_b3 | GTCCCTGCCTCTATATCTTTACAGGTCAGTACGGTGGCCCGGAGT |
| pigk_b3 | GACTTTGTGGGCACACTTTAAACAGTTCCACTCAACTTTAACCCG |
| pigk_b3 | GTCCCTGCCTCTATATCTTTCATTCATGTTGGCTTGGCTAGCAGG |
| pigk_b3 | CCAGAAACTCCAGCAAGTAAAAAGTTTCCACTCAACTTTAACCCG |
| pigk_b3 | GTCCCTGCCTCTATATCTTTACTTGTCCATCAAATGGACCCCAAT |
| pigk_b3 | GCTGATGTGAAAGTGAGTCCTCCCCTTCCACTCAACTTTAACCCG |
| pigk_b3 | GTCCCTGCCTCTATATCTTTCTTGGCTGCTAGCAAGAGCCATGAT |
| pigk_b3 | AAAACCTCTCATACATGGAGGCCCCTTCCACTCAACTTTAACCCG |
| pigk_b3 | GTCCCTGCCTCTATATCTTTGGCACGTGTCTATGATGAACAACAA |
| pigk_b3 | GTCTCTTTTGCCACATCTGCTCAAATTCCACTCAACTTTAACCCG |
| pigk_b3 | GTCCCTGCCTCTATATCTTTCATCAGCTAACTCCATGTTACTGAT |
| pigk_b3 | CCTGGAACTTCAGGAAGCCGTTTCCTTCCACTCAACTTTAACCCG |
| pigk_b3 | GTCCCTGCCTCTATATCTTTCATGACCTGTCAGGTAAATGAGAAT |
| pigk_b3 | CGTCTGACAGAAGACGTTTGGATCGTTCCACTCAACTTTAACCCG |
| pigk_b3 | GTCCCTGCCTCTATATCTTTGGGTACTGAGAGGGAGACGCCCCGT |
| pigk_b3 | GAAAGTTCTCTACCGTAACCTCATATTCCACTCAACTTTAACCCG |
| pigk_b3 | GTCCCTGCCTCTATATCTTTCGCGGTAATCCACCTCCACGTCGTC |
| pigk_b3 | TGAGCTCCATGTTCTTATGACTGAATTCCACTCAACTTTAACCCG |
| pigk_b3 | GTCCCTGCCTCTATATCTTTCAGTAGCTGGTTTGGGGTTTCTATA |
| pigk_b3 | TATCATCGGCTAGCATCAGAACAATTTCCACTCAACTTTAACCCG |
| pigk_b3 | GTCCCTGCCTCTATATCTTTGACTGTCGGGAATGCCCAGTCTCTT |
| pigk_b3 | ACACTGACAGCGTGTTCGCCACATGTTCCACTCAACTTTAACCCG |
| pigk_b3 | GTCCCTGCCTCTATATCTTTGATAATTAAACCAGAATCTAGAAGT |
| pigk_b3 | CTGCCCAGTTGTTAGTGTGTCCACTTTCCACTCAACTTTAACCCG |
| pigk_b3 | GTCCCTGCCTCTATATCTTTTAAAAAACTGTCCAGCTTTGGTCTC |
| pigk_b3 | TGTATGCGCATGGTATGGAAACAAATTCCACTCAACTTTAACCCG |
| pigk_b3 | GTCCCTGCCTCTATATCTTTGAAGAAGGCATGAATACACTGTAAA |
| pigk_b3 | AATCCATTATCGTTCGTCACCGGTATTCCACTCAACTTTAACCCG |
| pigk_b3 | GTCCCTGCCTCTATATCTTTAGTGACAGTGTAGATTTGCAATGAC |
| pigk_b3 | GCTATTTCTGACCAAACCTATCACATTCCACTCAACTTTAACCCG |
| pigk_b3 | GTCCCTGCCTCTATATCTTTCAAATATTGTGTGTGGAATCTTTTA |
| pigk_b3 | GTTTCAAGCGGTTGGTTCTTTGATCTTCCACTCAACTTTAACCCG |
| pigk_b3 | GTCCCTGCCTCTATATCTTTTGTGGAATCAGAAGATGTGTTTGAG |
| pigk_b3 | CGTAGGTCTGGAAGAACACCAGCAGTTCCACTCAACTTTAACCCG |
| pigk_b3 | GTCCCTGCCTCTATATCTTTTGAGAGTCCAGAGCCCAAGAATGAA |
| pigk_b3 | GGTGCCAGTCCTTCTGCTTGGGTTTTTCCACTCAACTTTAACCCG |
| pigk_b3 | GTCCCTGCCTCTATATCTTTGATGAATAATTTCAGCAACTGGAAG |
| pigk_b3 | ACGTAAAAGTTTCAACTTGTAGATCTTCCACTCAACTTTAACCCG |
| pigk_b3 | GTCCCTGCCTCTATATCTTTCACTTCCAGACCTCTGCATTGGCGA |
| pigk_b3 | ACGTCAGACTGATTGTGTCCTTGGTTTCCACTCAACTTTAACCCG |
| pigk_b3 | GTCCCTGCCTCTATATCTTTTCTCCACTTTCCGGACACTTCCGAA |
| pigt_b3 | CTCAAAACAAAACACTAGCAAAACATTCCACTCAACTTTAACCCG |
| pigt_b3 | GTCCCTGCCTCTATATCTTTAAAGTGATGGACAATCTTTGCTTGT |
| pigt_b3 | AAATCAAAGCAAACACAACCAAAACTTCCACTCAACTTTAACCCG |
| pigt_b3 | GTCCCTGCCTCTATATCTTTAATGAAAGACAATCGAAGCTTTTCT |
| pigt_b3 | AAAACACAGCAAAGAAAGTGACGGATTCCACTCAACTTTAACCCG |
| pigt_b3 | GTCCCTGCCTCTATATCTTTACCTTTGGTTTTGAGAAAAAACAAG |
| pigt_b3 | AACAGTCACTTGGCATCAAATCCTTTTCCACTCAACTTTAACCCG |
| pigt_b3 | GTCCCTGCCTCTATATCTTTTCCCAGAGCCGAAAAGCATCCAAAA |
| pigt_b3 | TAATATGGGCACTTGGGGGCATCAGTTCCACTCAACTTTAACCCG |
| pigt_b3 | GTCCCTGCCTCTATATCTTTCTGCTCTGGGCATTTATGTCAGCAA |
| pigt_b3 | TAGGGCATGCTGAAGTCAGGAGTGGTTCCACTCAACTTTAACCCG |
| pigt_b3 | GTCCCTGCCTCTATATCTTTAGATTTACCAGCAGGGGTTCAGTGT |
| pigt_b3 | TAAACAATGGCTGGTCCAGCGTGAATTCCACTCAACTTTAACCCG |
| pigt_b3 | GTCCCTGCCTCTATATCTTTTGGTGTTCATTGCAACCATACTTGG |
| pigt_b3 | ATCAGGGGTGTATTCAGTCCATTTGTTCCACTCAACTTTAACCCG |
| pigt_b3 | GTCCCTGCCTCTATATCTTTCAGAGCCCTATCAAACTGCACCGTG |
| pigt_b3 | TGAGGACGGTGGCGGTCTTTAGCCGTTCCACTCAACTTTAACCCG |
| pigt_b3 | GTCCCTGCCTCTATATCTTTTGGTAGTGAATGTAGCTGGGCTTGT |
| pigt_b3 | GGTACCAGGGCACTGAATCCATCAGTTCCACTCAACTTTAACCCG |
| pigt_b3 | GTCCCTGCCTCTATATCTTTGTACAGGAAACGCCCGAAACGGGTG |
| pigt_b3 | AGACACATAGCGTTCTGCATGCAGCTTCCACTCAACTTTAACCCG |
| pigt_b3 | GTCCCTGCCTCTATATCTTTCGGACGCAGCATATCACCGCTCTCT |
| pigt_b3 | TCAGCTCTGGTCAAGTCATAGACTGTTCCACTCAACTTTAACCCG |
| pigt_b3 | GTCCCTGCCTCTATATCTTTTATGTCCTGCGATCTCCTAGAACCA |
| pigt_b3 | CCTCAGGGTTGTCTGTGATGTCCACTTCCACTCAACTTTAACCCG |
| pigt_b3 | GTCCCTGCCTCTATATCTTTAGACTTTACTAGAGGAAGCAAGTGG |
| pigt_b3 | GCCACATTCCTAAATGTCCACTTTCTTCCACTCAACTTTAACCCG |
| pigt_b3 | CTTTTTGCCTTGTCCTGAAGTGTAGTTCCACTCAACTTTAACCCG |
| pigt_b3 | GTCCCTGCCTCTATATCTTTATCAAACACCACATTGAGAGTTTGC |
| pigt_b3 | GTCCCTGCCTCTATATCTTTTAGTTTTAGGAGTGTAGCATCTCTG |
| pigt_b3 | CTGATGTGAACAGCTTGGGAGTGGATTCCACTCAACTTTAACCCG |
| pigt_b3 | GTCCCTGCCTCTATATCTTTCTGCTGTGAAAAAGCTTCTCTGATT |
| pigt_b3 | TCCAGGGTGTCAGATTTTCAGTGCATTCCACTCAACTTTAACCCG |
| pigt_b3 | GTCCCTGCCTCTATATCTTTCAATTTCCCGTGGCAGCGTCGCATA |
| pigt_b3 | GGAAGCACTGGGCTGCACTGTGTTGTTCCACTCAACTTTAACCCG |
| pigt_b3 | GTCCCTGCCTCTATATCTTTCAGATCAATGAAGTTCAGAGATGCA |
| pigt_b3 | ACGTCGGTGACGGTGTGATGGAACCTTCCACTCAACTTTAACCCG |
| pigt_b3 | GTCCCTGCCTCTATATCTTTACCCAGAGCTCTGCTCCAGGAGGAG |
| pigt_b3 | TGAAGGAGATGTGCAGCTCACGCACTTCCACTCAACTTTAACCCG |
| pigt_b3 | GTCCCTGCCTCTATATCTTTAAAACTTGGAAAGCACTTGGCCCAG |
| pigt_b3 | GAAGTCAACATCCCAGAGTGTGCGGTTCCACTCAACTTTAACCCG |
| pigt_b3 | GTCCCTGCCTCTATATCTTTCTGGAAACTGGCATAGATGTCGCCG |
| pigt_b3 | GGTCTAACAGACGGGGTGTTTTGTGTTCCACTCAACTTTAACCCG |
| pigt_b3 | GTCCCTGCCTCTATATCTTTTCATCTCCTGTGTATGGTTCCACAT |
| pigt_b3 | CATTATCTCGCGGCGCGTCTGAAGCTTCCACTCAACTTTAACCCG |
| pigt_b3 | GTCCCTGCCTCTATATCTTTTTTGCTTTTCTCCTGTGGTGTCGTT |
| pigt_b3 | ACATAAGGCTTTTTTCCATGTATGTTTCCACTCAACTTTAACCCG |
| pigt_b3 | GTCCCTGCCTCTATATCTTTCTATCAGTAGGCCATACTACTACTT |
| pigt_b3 | GACAGTCTTCGACTCTGACAAAACCTTCCACTCAACTTTAACCCG |
| pigt_b3 | GTCCCTGCCTCTATATCTTTAAAAGATGGTCAATCGTGTCTTATG |
| pigu_b1 | GAGCTCTCAGTCCAGTGAAAGAAGGTAGAAGAGTCTTCCTTTACG |
| pigu_b1 | GAGGAGGGCAGCAAACGGAAATGTTCTCGGCGAAGATATGCGTAG |
| pigu_b1 | TGCCAGCGTAAATCCATAGATGCCATAGAAGAGTCTTCCTTTACG |
| pigu_b1 | ACATGAGGAACACTGGATGCTCCTTTAGAAGAGTCTTCCTTTACG |
| pigu_b1 | GAGGAGGGCAGCAAACGGAAGTTTGATGGAGAGCGGGATGGTGTA |
| pigu_b1 | AAGAGTCCGATGTTTGGAGTGAGGTTAGAAGAGTCTTCCTTTACG |
| pigu_b1 | AGTCCCAGGATCCCAAGAGGAAGAATAGAAGAGTCTTCCTTTACG |
| pigu_b1 | GAGGAGGGCAGCAAACGGAAAGAGTCCAGTGATCACGATCAGACT |
| pigu_b1 | GAGGAGGGCAGCAAACGGAAGCAAACAGTGTCAGAGGATACATGG |
| pigu_b1 | GAGGAGGGCAGCAAACGGAATGACTTTGCCACACATGACAGGATC |
| pigu_b1 | GAGGAGGGCAGCAAACGGAAGGGATGTAGAACATCTCCTTAGGAG |
| pigu_b1 | CCAAACAGTCAGCAGGATACCGTTCTAGAAGAGTCTTCCTTTACG |
| pigu_b1 | GAGGAGGGCAGCAAACGGAAAGCCACTGCTGTGATTCCATCTGCT |
| pigu_b1 | GAGGAGGGCAGCAAACGGAAAGAAGGAGTGTTAAAGGAGCCGCCA |
| pigu_b1 | ACCAGGAGAATCTGTCCAAAGTTGATAGAAGAGTCTTCCTTTACG |
| pigu_b1 | GAGGAGGGCAGCAAACGGAATCGATAAAGGTGACTCCAGGCTGGC |
| pigu_b1 | GATCTGGAAGACGCAAATGAAGAAGTAGAAGAGTCTTCCTTTACG |
| pigu_b1 | GAGGAGGGCAGCAAACGGAAGCGGAAGTGCTCAAACATCTCTGCG |
| pigu_b1 | GAACGCATACTGAAGAGTGAAGAACTAGAAGAGTCTTCCTTTACG |
| pigu_b1 | GAGGAGGGCAGCAAACGGAAGAAGCTGCTCCTCCTCAGGTTCACT |
| pigu_b1 | CTCTGCAGGAAGAACAGCAGTGCAGTAGAAGAGTCTTCCTTTACG |
| pigu_b1 | GAGGAGGGCAGCAAACGGAAAGGCACTACCTTTCAATGTACAGAG |
| pigu_b1 | GAGGAGGGCAGCAAACGGAACCAACAAGGCAAGACCCTCCACAAC |
| pigu_b1 | GAGGAGGGCAGCAAACGGAAACTTTAACACTAGAGTGGCTTCAGT |
| pigu_b1 | CTTTAGGTAGAGGCCGTGCGTCAGGTAGAAGAGTCTTCCTTTACG |
| pigu_b1 | GAGGAGGGCAGCAAACGGAAAGCAACGTGATGGCATAGTAGAAAT |
| pigu_b1 | GAGGAGGGCAGCAAACGGAAGTACAGGGAAGAGCGCAGAGCATGC |
| pigu_b1 | TGACACCAGAAAGATGTTTCTCAGATAGAAGAGTCTTCCTTTACG |
| pigu_b1 | TAGAGGGCGACATCTCCCACAGTGGTAGAAGAGTCTTCCTTTACG |
| pigu_b1 | GAGGAGGGCAGCAAACGGAATAGGACTTAAAGATGGAGATGATGG |
| pigu_b1 | CCAAAGCTAACAATATCCCACTTAGTAGAAGAGTCTTCCTTTACG |
| pigu_b1 | GATAATAGCATTGTTCAGCCCACAATAGAAGAGTCTTCCTTTACG |
| pigu_b1 | ATTCAACGGCGACACCACCTCGACCTAGAAGAGTCTTCCTTTACG |
| pigu_b1 | ACGCCTCTTATTGTAACAGCTACTATAGAAGAGTCTTCCTTTACG |
| pigu_b1 | GAGGAGGGCAGCAAACGGAAGGAACAGAAAGTATGAATCCATAGA |
| pigu_b1 | AACAGATAAAACATGGCAACCTTGATAGAAGAGTCTTCCTTTACG |
| pigu_b1 | GAGGAGGGCAGCAAACGGAAAAGAGGTATATCATAAGGGGTGTTT |
| pigu_b1 | CTCCTGAGTAAGGCGAGACCCCCAGTAGAAGAGTCTTCCTTTACG |
| pigu_b1 | GAGGAGGGCAGCAAACGGAACTCGGAAATTATTTCTGCCAAATTG |
| pigu_b1 | GAGGAGGGCAGCAAACGGAAATTCCAAGGCATATTTCTGTTTTCT |
| pigu_b1 | GTTATAAATCTGGACTGAGAGATAATAGAAGAGTCTTCCTTTACG |
| pigu_b1 | ATCTCTGCATAGTCAACCAAAAAATTAGAAGAGTCTTCCTTTACG |
| pmm2_b1 | GAGGAGGGCAGCAAACGGAAGCTGAAAAAGAGCTCCCTGCAGATC |
| pmm2_b1 | TTCAGGAGATTTGACCTCATGGCCGTAGAAGAGTCTTCCTTTACG |
| pmm2_b1 | CAAACAGTAGCGCTTATCCCAGCCTTAGAAGAGTCTTCCTTTACG |
| pmm2_b1 | GAAAGCTAGTCCCTTCCCAGCAAACTAGAAGAGTCTTCCTTTACG |
| pmm2_b1 | GAGGAGGGCAGCAAACGGAAGATTCTTTCCTGTTGGCTGCAGCTT |
| pmm2_b1 | GAGGAGGGCAGCAAACGGAATCCAAACCTGTACGCAACCAAACCG |
| pmm2_b1 | TCCTCCAACCACTCCGACTTTCACATAGAAGAGTCTTCCTTTACG |
| pmm2_b1 | GGTGGCTTTTTGACGAGCTGCTGTTTAGAAGAGTCTTCCTTTACG |
| pmm2_b1 | ATCCACGGTAGAGCCAGACATTGTGTAGAAGAGTCTTCCTTTACG |
| pmm2_b1 | CGTGTTGTCGGAAGTGTATTGAAGCTAGAAGAGTCTTCCTTTACG |
| pmm2_b1 | AGAGGTTGTGTTCAGGGTCTTTTCTTAGAAGAGTCTTCCTTTACG |
| pmm2_b1 | GAGGAGGGCAGCAAACGGAATGTTCTGGGGTCTACAAAGATTTCG |
| pmm2_b1 | TCCAGGCATTGTTTTGTCTCCAAAGTAGAAGAGTCTTCCTTTACG |
| pmm2_b1 | GAGGAGGGCAGCAAACGGAAATGGATGTGCTGATAGGAGTCTTTC |
| pmm2_b1 | AGAGACATTCAGCATTCCATTCCGATAGAAGAGTCTTCCTTTACG |
| pmm2_b1 | CTGAATATTCTGCACAGAGTGAAGCTAGAAGAGTCTTCCTTTACG |
| pmm2_b1 | CACGTAATCCACTCTGTCAATCACATAGAAGAGTCTTCCTTTACG |
| pmm2_b1 | GAGGAGGGCAGCAAACGGAAAGTCCCGTCCACGTCAAACAGACAC |
| pmm2_b1 | GAGGAGGGCAGCAAACGGAACAGTTCATTCATTCAGCAGTGCAAC |
| pmm2_b1 | GAGGAGGGCAGCAAACGGAACAGAGCTGTACGTCAGAGTGAGCGA |
| pmm2_b1 | ACAACATCTGCCAACAGCAGACAAATAGAAGAGTCTTCCTTTACG |
| pmm2_b1 | GTATGGGCCACGTTGTTATTGTAGTTAGAAGAGTCTTCCTTTACG |
| pmm2_b1 | GAGGAGGGCAGCAAACGGAAACAGTGTAGTCTTCGGTATACTTCA |
| pmm2_b1 | GAGGAGGGCAGCAAACGGAAATCTCCAAGCTGCTCTTTGATTTTG |
| pmm2_b1 | GAAGTTTTATGAAGACTCAGTGGCATAGAAGAGTCTTCCTTTACG |
| pmm2_b1 | GAGGAGGGCAGCAAACGGAATGGAAACACATCAAAACTGATCTGT |
| pmm2_b1 | CTTTGAAAGGTAATTGAGGCAGAAATAGAAGAGTCTTCCTTTACG |
| pmm2_b1 | GAGGAGGGCAGCAAACGGAAGATGAATTCCTGCAAAATGTCTTCT |
| pmm2_b1 | GAGGAGGGCAGCAAACGGAACTGTTTCAGTTTAGACAGAAACTCA |
| pmm2_b1 | CTCCTTCTTATCAAGTTCAAAGAATTAGAAGAGTCTTCCTTTACG |
| pmm2_b1 | GAGGAGGGCAGCAAACGGAAATGACAATGCGACCAATTCAAATTT |
| pmm2_b1 | TTTCATTCATAGTTTCACGATTCATTAGAAGAGTCTTCCTTTACG |
| pmm2_b1 | GAGGAGGGCAGCAAACGGAATTCTTTTAAAACGGAGACAAATGTT |
| pmm2_b1 | GAGGAGGGCAGCAAACGGAATTCAATAAATGTGCCTCTTTTTTTA |
| pmm2_b1 | GAGGAGGGCAGCAAACGGAAACTTCATAAATAAGAAGAATAACCT |
| pmm2_b1 | GAGGAGGGCAGCAAACGGAAATTTGGAAACTTTAGAAATCAATAT |
| pmm2_b1 | TTTTTTTTTTTTTTTTGGTTTTTTTTAGAAGAGTCTTCCTTTACG |
| pmm2_b1 | GAGGAGGGCAGCAAACGGAATTTTTTTTTTTTTTTTTTTTTTTTC |
| pmm2_b1 | TTTTTTTTTTTTTTTTTTTTTTTTTTAGAAGAGTCTTCCTTTACG |
| pmm2_b1 | GAGGAGGGCAGCAAACGGAATTTTTTTTTTTTTTTTTTTTTTTTT |
| slc35a1_b2 | TCTCCCAGTGCACCTCCTGATGTATAAATCATCCAGTAAACCGCC |
| slc35a1_b2 | CCTCGTAAATCCTCATCAAATGGTCAGTTTCACAGGTCCTGGATC |
| slc35a1_b2 | CCTCGTAAATCCTCATCAAAGTAGATGGACACACACACCAGCAGA |
| slc35a1_b2 | CGCAGAGAAGCCCTTCATGATGTTAAAATCATCCAGTAAACCGCC |
| slc35a1_b2 | CCTCGTAAATCCTCATCAAAAGTGTATTTGACCACCACAGACGTG |
| slc35a1_b2 | CCTCGTAAATCCTCATCAAACCAGGGTGTGTATCCGTAGAAGAAG |
| slc35a1_b2 | GTTTCTGACCCACAGAGACGTGTCTAAATCATCCAGTAAACCGCC |
| slc35a1_b2 | CCTCGTAAATCCTCATCAAAGCTCTTCAGGACCTTCTCGAAGTAC |
| slc35a1_b2 | CCTCGTAAATCCTCATCAAACATGTCTGTGTTGTGTCGGTGACTC |
| slc35a1_b2 | GTTCTGTCACACTGAGATCAGCTTAAAATCATCCAGTAAACCGCC |
| slc35a1_b2 | CCTCGTAAATCCTCATCAAATGTGTCGTCTTGTTCTGCTCCAGCC |
| slc35a1_b2 | AGTGTCCTGTTTGGGCAGACCGTACAAATCATCCAGTAAACCGCC |
| slc35a1_b2 | ACTCGCCAAAAAGACCACCAGGCAGAAATCATCCAGTAAACCGCC |
| slc35a1_b2 | CCTCGTAAATCCTCATCAAACATGAGGGTCACAGCAATCCCAGAC |
| slc35a1_b2 | CCTCGTAAATCCTCATCAAACATGAAGCCCAGGAACGGGTTTTGC |
| slc35a1_b2 | CCTCGTAAATCCTCATCAAACAGCTTCAGCAGCTCTTTGGGACTC |
| slc35a1_b2 | CCTCGTAAATCCTCATCAAAATCTCCGGTCTCTCTGACCAGCATG |
| slc35a1_b2 | CGTCAACGTGATCTGCAGGCCGAAGAAATCATCCAGTAAACCGCC |
| slc35a1_b2 | CCTCGTAAATCCTCATCAAACATCAGCACCGTACACAGCGCCGTA |
| slc35a1_b2 | CTGGTATGTCACCTGGTAGACGGCGAAATCATCCAGTAAACCGCC |
| slc35a1_b2 | CCTCGTAAATCCTCATCAAAGTCCAGGTTGCTCAGGGCCACAAAG |
| slc35a1_b2 | CAGCTTGATGATCTCGGCCAGGCAGAAATCATCCAGTAAACCGCC |
| slc35a1_b2 | CCTCGTAAATCCTCATCAAAGGCGGTGGTGGAGAAGTAGAGCTCT |
| slc35a1_b2 | GGTGTACCTGAGGGCCACGGTGTATAAATCATCCAGTAAACCGCC |
| slc35a1_b2 | GAAGAGCACACTCACCGACTCGCTCAAATCATCCAGTAAACCGCC |
| slc35a1_b2 | TCGGGCCCCATCCGTCATGTAGACGAAATCATCCAGTAAACCGCC |
| slc35a1_b2 | CCTCGTAAATCCTCATCAAACAGCGTGACTCCCGCACACAGCATG |
| slc35a1_b2 | GATGGCGTAGACCACTGACGGGACGAAATCATCCAGTAAACCGCC |
| slc35a1_b2 | CCTCGTAAATCCTCATCAAAGGCGGCGATCAGGGTCATCACAGTC |
| slc35a1_b2 | CTGCAGGCGGCTCAGAGAGCGGTTCAAATCATCCAGTAAACCGCC |

**Supplementary Table 3: Probe sequences used for HCR in this study.**

HCR probes were designed as oligonucleotide pools. The sequences in each pool are listed in this table.
